# Development of a PET radioligand for α2δ-1 subunit of calcium channels for imaging neuropathic pain

**DOI:** 10.1101/2022.03.09.483673

**Authors:** Yu-Peng Zhou, Yang Sun, Kazue Takahashi, Vasily Belov, Nick Andrews, Clifford J. Woolf, Pedro Brugarolas

**Author notes:** **Correspondence:** Pedro Brugarolas, PhD, 55 Fruit St, Bulfinch 051, Boston, MA 02114. The Salk Institute for Biological Studies, San Diego, California (Nick Andrews).

## Abstract

Neuropathic pain affects 7-10% of the adult population. Being able to accurately monitor biological changes underlying neuropathic pain will improve our understanding of neuropathic pain mechanisms and facilitate the development of novel therapeutics. Positron emission tomography (PET) is a noninvasive molecular imaging technique that can provide quantitative information of biochemical changes at the whole-body level by using radiolabeled ligands. One important biological change underlying the development of neuropathic pain is the overexpression of α2δ-1 subunit of voltage-dependent calcium channels (the target of gabapentin). Thus, we hypothesized that a radiolabeled form of gabapentin may allow imaging changes in α2δ-1 for monitoring the underlying pathophysiology of neuropathic pain. Here, we report the development of two ^18^F-labeled derivatives of gabapentin (*trans-*4-[^18^F]fluorogabapentin and *cis-*4-[^18^F]fluorogabapentin) and their evaluation in healthy rats and a rat model of neuropathic pain (spinal nerve ligation model). Both isomers were found to selectively bind to the α2δ-1 receptor with *trans-*4-[^18^F]fluorogabapentin having a higher affinity. Both tracers displayed around 1.5- to 2-fold increased uptake in injured nerves over the contralateral uninjured nerves when measured by gamma counting *ex vivo*. Although the small size of the nerves and the signal from surrounding muscle prevented visualizing these changes using PET, this work demonstrates that fluorinated derivatives of gabapentin retain binding to α2δ-1 and that their radiolabeled forms can be used to detect pathological changes *in vitro* and *ex vivo*. Furthermore, this work confirms that α2δ-1 is a promising target for imaging specific features of neuropathic pain.

## Introduction

Defined by the International Association for the Study of Pain (IASP), neuropathic pain is the “pain caused by a lesion or disease of the somatosensory nervous system”.^1^ An epidemiological study suggests that the prevalence rate for neuropathic pain is between 6.9% and 10%.^2^ The clinical assessment of pain is mainly based on self-report^3^ whereas the preclinical assessment relies on measuring physiological/behavioral response to evoking stimuli.^4, 5^ Pain imaging using electroencephalography (EEG), magnetic resonance imaging (MRI), or positron emission tomography (PET) can provide additional information of structure, function, or function underlying the pain condition, which can help us better understand the mechanisms and biochemical processes contributing to the pathological pain^6-10^ and help us develop nonopioid analgesics.^11, 12^ Compared with EEG and MRI, PET can be used to visualize biochemical events *in vivo* at a whole-body dimension and to quantify the density and occupancy of the neuroreceptors or therapeutic targets with high sensitivity and moderate spatial resolution.^13^ PET tracers such as [^18^F]FDG, [^11^C]diprenorphine/[^18^F]FDPN, 6-[^18^F]fluoro-L-DOPA, [^11^C]ABP688, and [^18^F]FTC-146 have been used to image glucose uptake,^14-16^ opioid receptor density,^17-19^ dopaminergic activity,^20^ glutamate receptor 5 (mGluR5) binding level,^21^ and the expression of sigma 1 receptor^22-24^ under different pain conditions. However, the development of neuropathic pain involves other pathways and receptors^25^ which have not been sufficiently studied by PET imaging. Therefore, developing novel tracers for other targets specific to neuropathic pain is pressingly needed.

The α2δ subunit is one of the auxiliary components of voltage-gated calcium channels (VGCCs). α2δ subunits have four gene types, which encode subunits α2δ-1, α2δ-2, α2δ-3, and α2δ-4. α2δ-1 subunit is mainly expressed in muscles, central nervous system (CNS), peripheral nervous system (PNS), and endocrine tissues.^26^ Multiple studies have shown that the expression of α2δ-1 is highly increased (3-17 fold) in the nerves, dorsal root ganglia and spinal cords of different animal models of neuropathic pain.^27-37^ In addition, transgenic mice overexpressing α2δ-1 display a phenotype of hyperalgesia^38^ and α2δ-1 knock-out mice show reduced sensitivity to mechanical and cold stimuli.^39^ Remarkably, the increased expression of α2δ-1 in the spinal dorsal horn of neuropathic pain animals can be reversed by treatment with gabapentinoids.^30, 40, 41^ Taken together, these results demonstrate that α2δ-1 is a robust biomarker of neuronal injury associated with neuropathic pain. Despite the potential value of neuroimaging α2δ-1 receptor by PET, this has not been achieved, mainly due to the lack of a suitable PET tracer. In 2019, Yang *et al*. communicated at the SNMMI annual meeting the development of a copper-64 labeled antibody for the α2δ-1 receptor.^42^ This antibody-based PET tracer showed promise for cancer imaging, however its ability to detect changes in nerves or spinal cord due to neuropathic pain has not been reported.

Gabapentinoids are ligands of α2δ receptors and the first-line treatment of neuropathic pain recommended by the International Association for the Study of Pain (IASP) Neuropathic Pain Special Interests Group (NeuPSIG) and European Federation of the Neurological Societies (EFNS).^25^ FDA approved gabapentinoids include gabapentin (commercial name Neurontin) and pregabalin (commercial name Lyrica) (**Scheme 1**). Given the efficacy of these compounds and their selectivity towards α2δ receptors, we hypothesized that radiolabeled gabapentinoids may serve as PET tracers for neuropathic pain imaging. In particular, gabapentin appears to have suitable properties including high affinity towards α2δ-1 receptor, high metabolic stability, being brain penetrant and is amenable to chemical modifications.^43, 44^ Furthermore, fluorine-18 labeled tracers are generally preferred over carbon-11 tracers as their longer half-life (110 min *vs*. 20 min) allows for remote production and regional distribution. Therefore, we set out to develop novel fluorinated derivatives of gabapentin and their ^18^F-labeled versions. In this paper, we describe the synthesis, and the evaluation of two novel radiotracers in healthy rats and rats post spinal nerve ligation (SNL), a model of peripheral neuropathic pain.

**Scheme 1.**
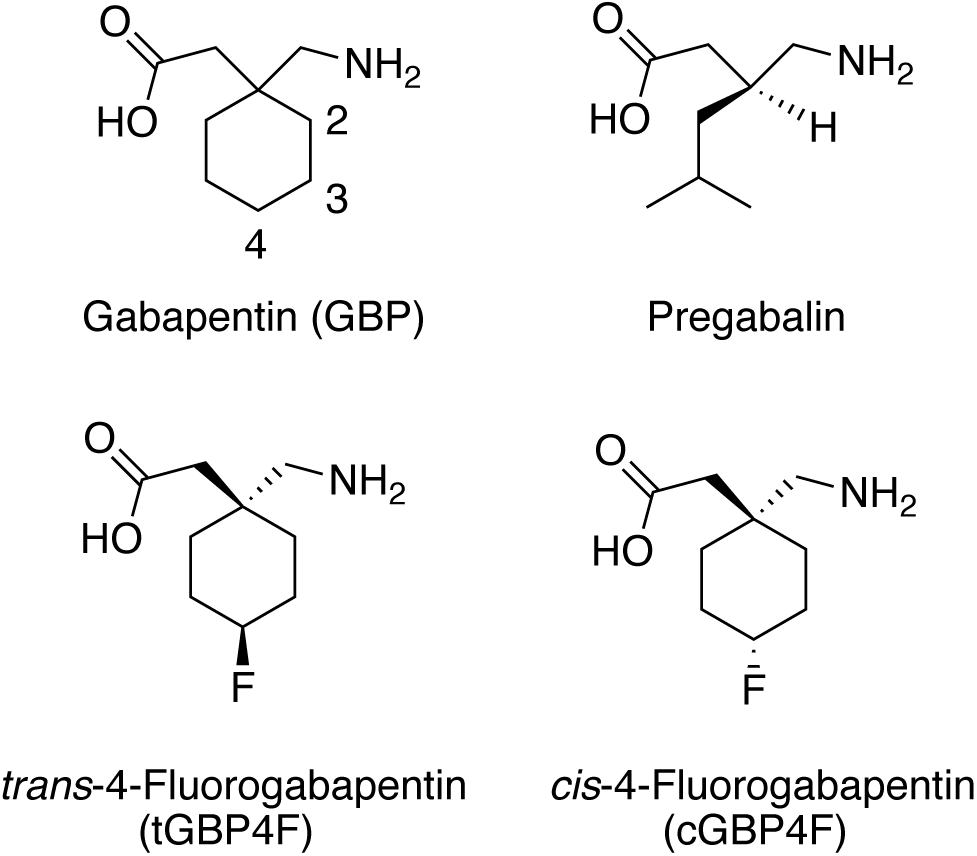
Chemical structures of gabapentin, pregabalin, and 4-fluorogabapentin.

## RESULTS AND DISCUSSION

### Validation of gabapentin as a basis for a neuropathic pain tracer

As described above, the expression of α2δ-1 increases in animal models of peripheral neuropathic pain.^27-37^ Before investing effort in generating new ^18^F-labeled ligands and testing them *in vivo*, we decided to test whether the increased expression in α2δ-1 in the spinal cord of mice with neuropathic pain would result in a measurable increase in gabapentin binding. To do this, we incubated spinal cord sections obtained from spared nerve injury (SNI) mice, a well-established model of neuropathic pain in which two of the three distal branches at the foot level of the sciatic nerve are transected and ligated,^45^ with tritium labeled gabapentin ([^3^H]GBP). Incubation of longitudinal spinal cord sections showed a 245% increase in [^3^H]GBP binding in the spinal cord of SNI animals compared to sham operated mice (**Fig. 1A**). Autoradiography of axial spinal cord sections showed a binding pattern consistent with the previously reported expression of α2δ-1^46^ and a 160% higher binding in SNI animals than controls (**Fig. 1B**). Furthermore, closer inspection of the signal in SNI sections showed 38% higher binding in the dorsal horn ipsilateral to the injured nerve than in the dorsal horn of the contralateral side (**Fig. 1C**).

**Figure 1.**
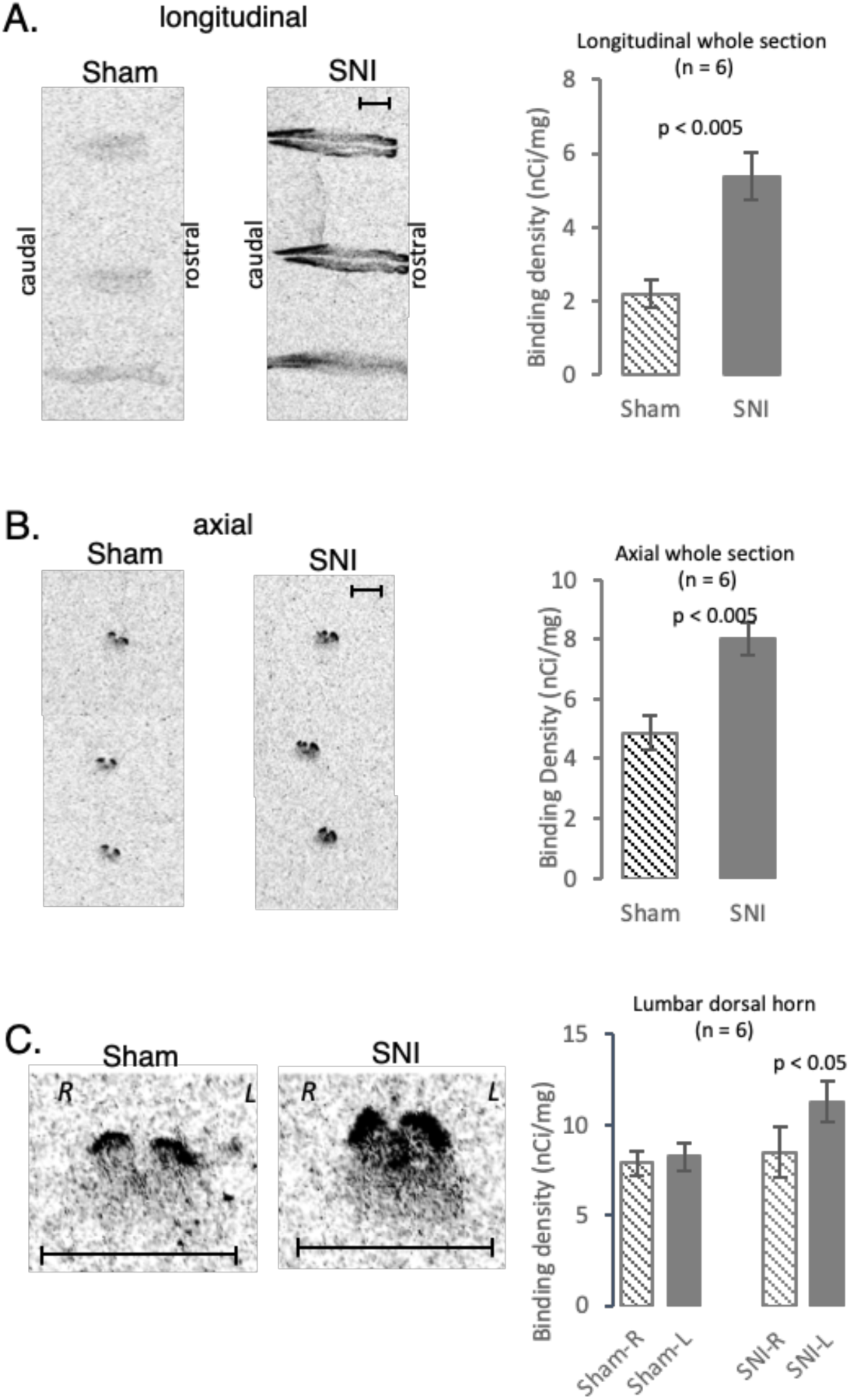
Binding of [^3^H]GBP in the spinal cord of nerve injured mice. **A**. Representative *in vitro* [^3^H]GBP autoradiography of longitudinal spinal cord sections of spared nerve injury (SNI) mice and sham controls. SNI animals showed a 245% increase in [^3^H]GBP binding in whole spinal cord sections. **B**. Representative *in vitro* [^3^H]GBP autoradiography of axial spinal cord sections of spared nerve injury (SNI) mice and sham controls. SNI animals showed a 160 % increase in [^3^H]GBP binding in whole spinal cord sections. **C**. Representative zoom in *in vitro* [^3^H]GBP autoradiography images of axial spinal cord sections of spared nerve injury (SNI) mice and sham controls. SNI animals showed a 138% higher [^3^H]GBP binding in the dorsal horn ipsilateral to the injury than the contralateral side.

### Chemistry and Radiochemistry

Encouraged by the ability of [^3^H]GBP to detect changes in a neuropathic pain model, we decided to pursue fluorinated derivatives of gabapentin. Fluorinated derivatives of gabapentin have not been previously described. Previous structure-activity relationship (SAR) studies of gabapentin alkyl derivatives have indicated that the *trans-*4-substituted, (1S,3R)-, and (1R,3S)-3-substituted gabapentin (substitution = methyl or ethyl) show similar binding affinities to the α2δ-1 compared to the parent drug gabapentin.^47^ Based on these SAR, we hypothesized that addition of a fluorine atom to the 4-position would retain binding to the target. The methods to synthesize 4-fluorogabapentin both with nonradioactive fluorine-19 and radioactive fluorine-18 are discussed here.

**Scheme 2.**
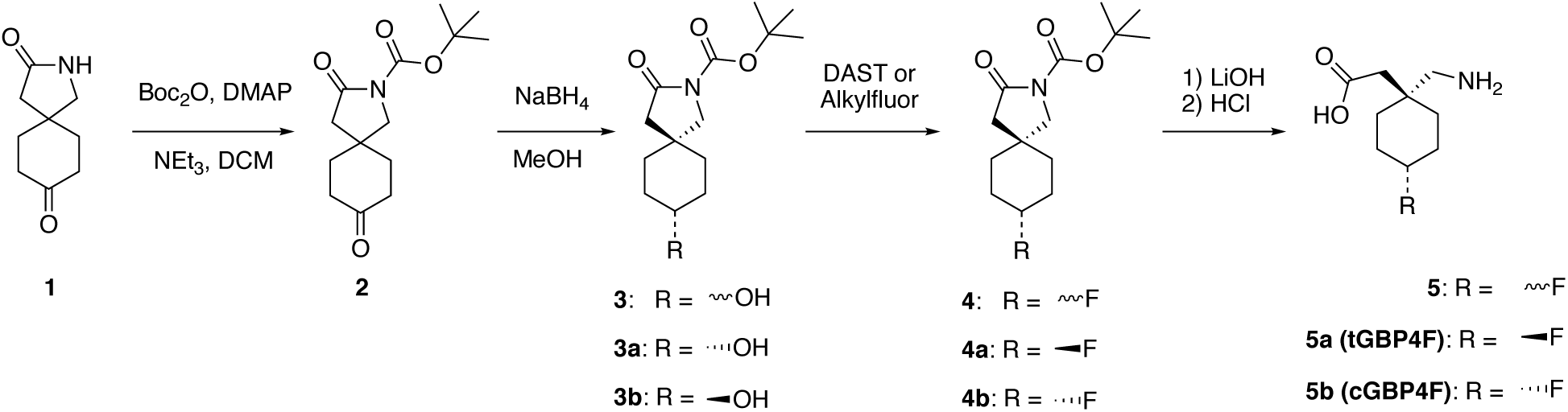
Synthesis of *trans*-4-fluorogabapentin (**5a**, tGBP4F) and *cis*-4-fluorogabapentin (**5b**, cGBP4F).

As shown in **Scheme 2**, the synthesis of nonradioactive 4-fluorogabapentin starts from the commercially available precursor 2-azaspiro[4.5]decane-3,8-dione (**1**). First, the amide -NH is protected by *tert-*butyloxycarbonyl (Boc) by the reaction of **1** with di-*tert*-butyl dicarbonate (Boc_2_O). The subsequent reduction of the Boc-protected intermediate **2** with excess amount of sodium borohydride (NaBH_4_) in methanol (MeOH) yielded the *trans*- and *cis*-mixture of compound **3** in 97% chemical yield. The isolation of *trans*- (**3b**) and *cis*- (**3a**) isomers was achieved by reverse-phase semi-preparative HPLC. Deoxyfluorination of **3a** and **3b** using DAST (diethylaminosulfur trifluoride) or AlkylFluor yielded the fluorinated Boc-protected gabapentin lactam **4a** and **4b**, respectively. AlkylFluor showed better deoxyfluorination efficiency (40-50%) than DAST (10%), however, it also resulted in more side products. Finally, the hydrolysis and deprotection of **4a** and **4b** in one pot yielded the final products *trans*-4-fluorogabapentin (**5a**, tGBP4F) and *cis*-4-fluorogabapentin (**5b**, cGBP4F) in 70-80 % yield.

In order to prepare the ^18^F-labeled products, cis/trans mixture **3** or isomerically pure **3a**/**3b** were reacted with MsCl (methanesulfonyl chloride) in dichloromethane in the presence of NEt_3_ to give the mesylated compounds **6** and **6a**/**6b** (Scheme 3). Stereochemically pure **6a** and **6b** can be obtained both through direct mesylation of the stereochemically pure precursors (**3a**/**3b**) or by the semi-prep reverse-phase HPLC purification of compound **6**. The stereochemical assignment of **6a** and **6b** was determined based on the large vicinal H-H coupling^48^ of 4-position proton and 3/5-position proton of **6a** [H_4_ δ = 4.77 ppm (tt, *J*_*1*_ = 4 Hz, *J*_*2*_ = 8 Hz, 1 H)] and small vicinal H-H coupling of 4-position proton of **6b** [H_4_ δ = 4.80 ppm (tt, *J*_*1*_ = 5 Hz, *J*_*2*_ = 5 Hz, 1 H)] as well as the NOESY H-H coupling of lactam ring proton with axial proton of cyclohexyl ring (NMR spectra section, **Supporting Information**). Both assignment methods provided consistent and unequivocal results. The stereochemical assignments of **3a**/**3b, 4a**/**4b**, and **5a**/**5b** were based on the assignment of **6a**/**6b** after confirming experimentally that these transformations were stereoselective.

**Scheme 3.**
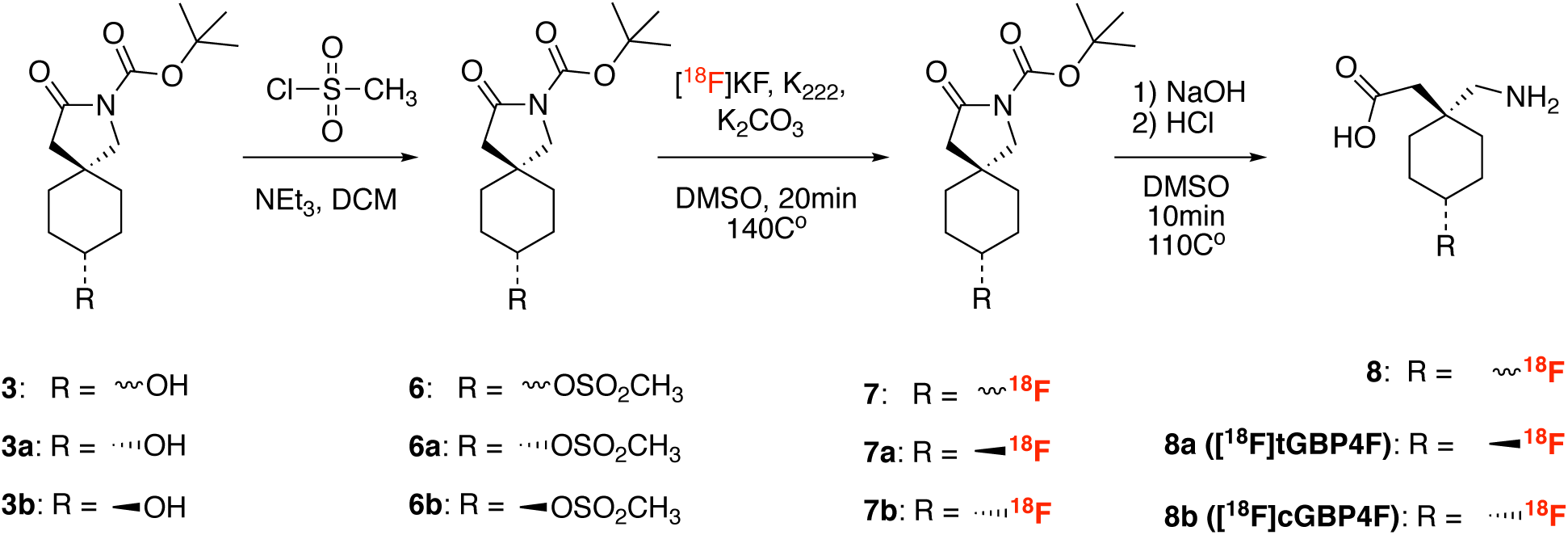
Synthesis of *trans*-[^18^F]4-fluorogabapentin ([^18^F]tGBP4F, **8a**) and *cis*-[^18^F]4-fluorogabapentin ([^18^F]cGBP4F, **8b**).

Fluorine-18 labeled gabapentin analogues were synthesized by nucleophilic substitution of [^18^F]F^-^ of the mesylate precursors **6** or **6a**/**6b**. As shown in **Scheme 3**, labeling of precursor **6** led to the isomer mixture [^18^F]fluorogabapentin (**8**, [^18^F]GBP4F) which was further purified to obtain the stereochemically pure *trans*-4-[^18^F]fluorogabapentin (**8a**, [^18^F]tGBP4F) and *cis*-4-[^18^F]fluorogabapentin (**8b**, [^18^F]cGBP4F). On the other hand, labeling of the stereochemically pure precursors **6a** and **6b** yielded stereochemically pure *trans*-4-[^18^F]fluorogabapentin (**8a**, [^18^F]tGBP4F) and *cis*-4-[^18^F]fluorogabapentin (**8b**, [^18^F]cGBP4F) directly (**Scheme 3**).

Labeling of precursor **6** with 18.5-55.5 GBq (0.5–1.5 Ci) of [^18^F]KF generated [^18^F]GBP4F (**8**) in 5.8 ± 1.8 % (n = 9) decay corrected radiochemical yield and >99% of radiochemical purity in ∼120 min of synthesis and purification time. The ratio of [^18^F]tGBP4F to [^18^F]cGBP4F was *ca*. 2:1. Labeling of precursor **6a** with 11.1-37 GBq (0.3–1.0 Ci) of [^18^F]KF generated [^18^F]tGBP4F (**8a**) in 4.3 ± 2.2 % (n = 13) decay corrected radiochemical yield and >99% of radiochemical purity. Labeling of precursor **6b** generated [^18^F]cGBP4F (**8b**) in 1.4 ± 1.2 % (n = 3) decay corrected radiochemical yield and >99% of radiochemical purity. Due to the low UV-Vis absorbance of the 4-fluorogabapentin, the identity of the radioactive compounds was confirmed by coinjection with reference standards on analytical radioHPLC and monitoring by evaporative light-scattering (ELS) detection (**Fig. S1**, Supporting Information).

### *In vitro* evaluations

In order to assess the target binding of the newly synthesized α2δ-1 ligands tGBP4F and cGBP4F, competitive radioligand binding assays to rat spinal cord sections were carried out and analyzed via quantitative *in vitro* autoradiography.

As shown in **Fig. 2A**, the autoradiography of free [^18^F]tGBP4F and [^18^F]cGBP4F showed an accumulation of radioactivity in the dorsal horn of the spinal cord sections, consistent with immunohistochemistry (**Fig. S2**, Supporting Information). This was also consistent with previously reported α2δ-1 immunohistochemistry results^46^ and the [^3^H]gabapentin autoradiography results (**Fig. 1**). As shown in **Fig. 2A**, >90% binding of [^18^F]tGBP4F and [^18^F]cGBP4F could be displaced by 10 μM of tGBP4F/cGBP4F or 10 μM of GBP, which demonstrates high specific binding.

**Figure 2.**
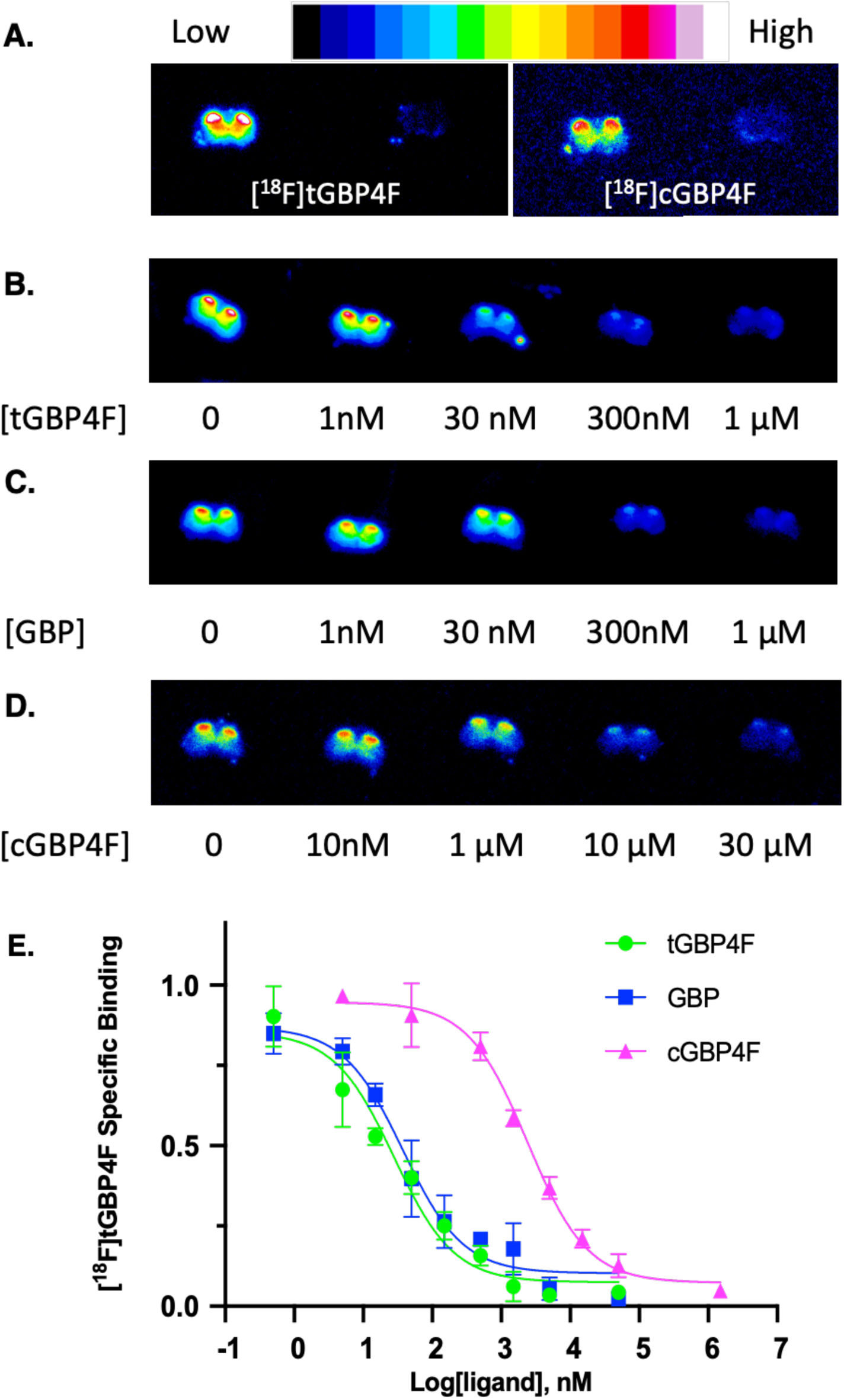
Binding of tGBP4F and cGBP4F to α2δ-1 receptors. **A:** Comparison of autoradiography of [^18^F]tGBP4F (left: baseline; right: blocking w/ 10 μM tGBP4F) and [^18^F]cGBP4F (left: baseline; right: blocking w/ 10 μM cGBP4F) in healthy rat spinal cord slices. Areas of high 4-[^18^F]fluorogabapentin binding correlate with published IHC results of high α2δ-1 expression area.^46^ **B-D:** Quantitative autoradiography of [^18^F]tGBP4F with increasing concentrations of tGBP4F (**B**), GBP (**C**), and cGBP4F (**D**) in healthy rat spinal cord slices. **E:** Competition binding curve for the inhibition of the [^18^F]tGBP4F binding radioligand binding by increasing concentrations of tGBP4F, GBP, and cGBP4F.

To assess the relative potency of the gabapentin derivatives, the half maximal inhibitory concentration (IC_50_) of tGBP4F, cGBP4F and GBP was measured by competitive radioligand binding against [^18^F]tGBP4F. Rat spinal cord sections were incubated with [^18^F]tGBP4F and increasing concentrations of non-radioactive competitors tGBP4F, GBP, and cGBP4F. As shown in **Fig. 2B-2D**, the [^18^F]tGBP4F binding was gradually inhibited by the non-radioactive competitors. The IC_50_ values of each competitors were calculated accordingly (**Fig. 2E**, IC_50_: tGBP4F = 28 ± 15 nM, n = 4; GBP = 38 ± 16 nM, n = 4; cGBP4F = 2415 ± 535 nM, n = 4). These results indicated that *trans-*isomer tGBP4F is slightly more potent than parent drug gabapentin and that the *cis*-isomer cGBP4F is significantly less potent than the *trans-*isomer. This observation is consistent with previously reported structure-activity relationship studies of alkyl derivatives of gabapentin, in which *trans-*4-substituted gabapentin (substitution = methyl or ethyl) showed slightly lower binding affinity than GBP whereas the *cis-*4-substituted gabapentin (substitution = methyl) showed significantly lower affinity.^47^

### *In vivo* evaluation in healthy rats

Being able to produce the ^18^F-labeled versions and encouraged by the selective and potent binding of [^18^F]tGBP4F and [^18^F]cGBP4F, we set out to evaluate these novel radioligands in live rats. A 60 minute dynamic whole body PET scans followed by a 15 minute CT scan was carried out on tracer-injected rats using a microPET/CT scanner. As shown in **Fig. 3A** and **Sup. Fig. S3**, both radiotracers showed wide distribution throughout the body with highest signal in bladder and kidneys. In the brain, [^18^F]tGBP4F showed a standard uptake value (SUV) of 0.7 at 10 min which decreased to 0.4 by 50 min. In contrast, [^18^F]cGBP4F brain SUV was 0.7 at 10 min and increased to 0.9 by 50 min. Higher SUV values and similar trend over time was observed in muscle tissue (**Fig. 3B**). In the kidneys, the signal from [^18^F]tGBP4F and [^18^F]cGBP4F decreased from SUV = 17 at 10 min to SUV = 8 at 50 min and from 14 to 12, respectively. Consistently, the signal in the bladder for [^18^F]tGBP4F and [^18^F]cGBP4F increased from 30 to 105 SUV and 11 to 26 SUV within the first 60 min, respectively. Taken together, these results showed limited brain uptake of both radioligands and faster clearance of [^18^F]tGBP4F compared to [^18^F]cGBP4F.

**Figure 3.**
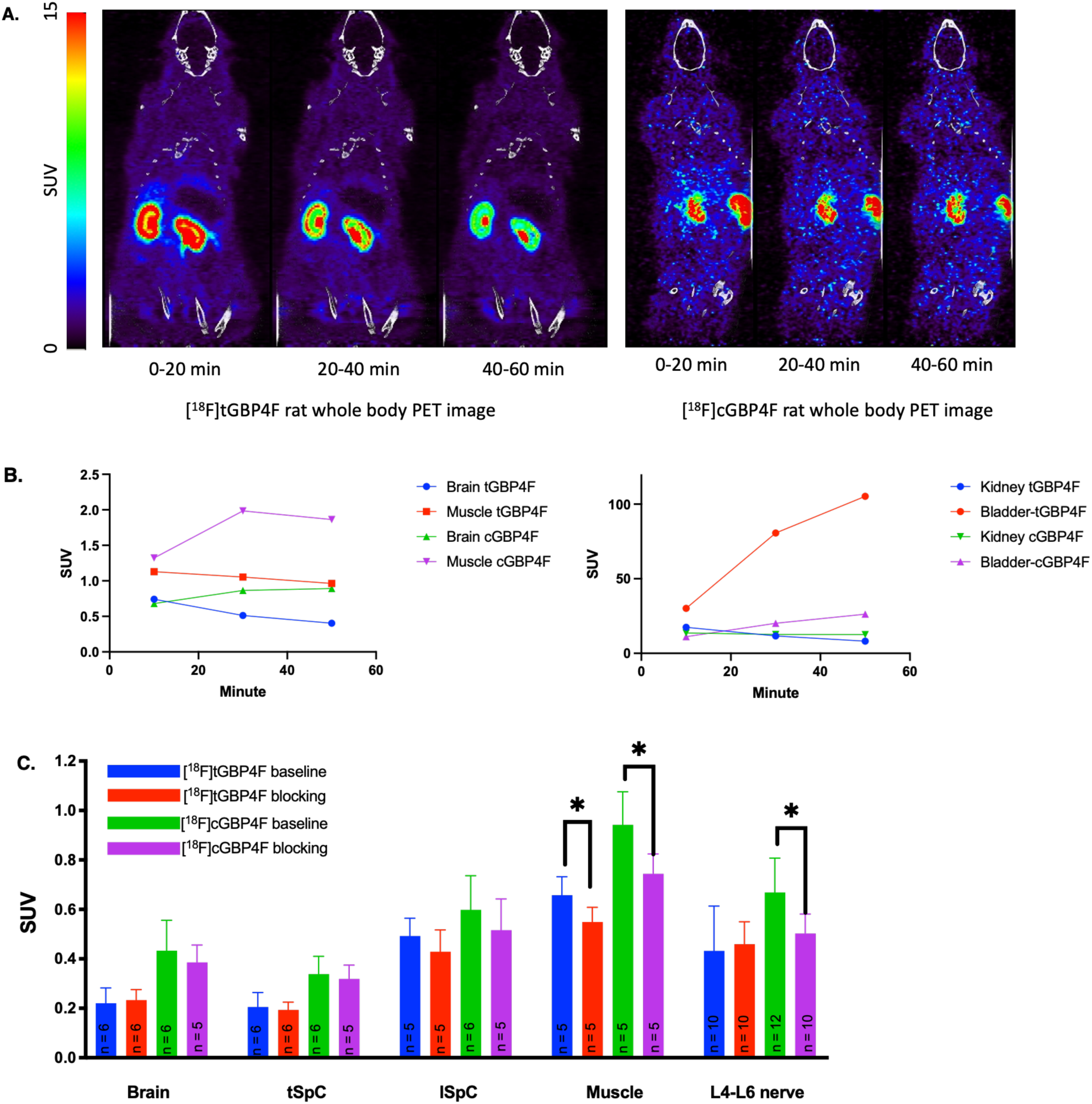
[^18^F]tGBP4F and [^18^F]tGBP4F in rats. **A:** Whole body PET imaging of healthy rats (coronal view, summed images at 0−20 min, 20−40 min, and 40−60 min intervals) with [^18^F]tGBP4F (left) and [^18^F]cGBP4F (right); **B:** PET imaging based time-activity curves of [^18^F]tGBP4F and [^18^F]cGBP4F in healthy rats. **C:** Biodistribution of [^18^F]tGBP4F and [^18^F]cGBP4F in brain, thoracic spinal cord (tSpC), lumbar spinal cord (lSpC), muscle, and L4-L6 spinal nerves of healthy rats (blocking dose: 30mg/kg of gabapentin, 30 min pre-injection) at 75 min post tracer injection (data obtained by the gamma counting of the dissected tissues, *: p < 0.05).

Due to the relatively small size of the rat spinal cord and spinal nerves, it is very difficult to accurately measure the SUV of these tissues using microPET imaging. To accurately measure the SUV in these tissues, rats were injected with the tracer and euthanized 75 min after injection. Their tissues were immediately dissected and the radioactivity in the tissues was measured by gamma counting. SUVs were calculated accordingly. This was done both under baseline (tracer-only) condition as well as 30 min after IP injection of 30 mg/kg of GBP (blocking) to assess specific binding *in vivo*. As shown in **Fig 3C** and consistent with the previous PET results, [^18^F]tGBP4F showed lower SUV than [^18^F]cGBP4F across tissues. The signal in the thoracic spinal cord was similar to that of the brain and the signal in the lumbar spinal cord was 1.7-2.4-fold greater. The signal in the spinal nerves was similar to that of the lumbar spinal cord. Finally, the signal in muscle (where α2δ-1 receptors are also highly expressed) was 10-20% greater than the signal in the nerves and lumbar spinal cord. Under blocking conditions, the signals of [^18^F]tGBP4F and [^18^F]cGBP4F in the muscle showed a -21% (n = 5, p < 0.05) and -16% (n = 5, p < 0.05) decrease, respectively. In the L4-L6 spinal nerves, [^18^F]cGBP4F also showed a -21% decrease (n = 10, p < 0.05) but this decrease was not observed with [^18^F]tGBP4F. Finally, the signals of both tracers in the brain and spinal cord did not show significant difference between baseline and blocking condition possibly due to the lower brain and spinal cord tracer uptake compared to muscle and peripheral nerves.

### *In vivo* and *ex vivo* evaluation in animal model of neuropathic pain (SNL rats)

To check if the tracers could detect increases in α2δ-1 expression under a neuropathic pain condition, both tracers were evaluated in rats after a spinal nerve ligation (SNL). The ligation was performed by unilaterally ligating the L5 and L6 spinal nerves.^49^ One-hour dynamic PET scans with [^18^F]tGBP4F and [^18^F]cGBP4F were carried out using a microPET/CT scanner ∼2 weeks after surgery. PET imaging showed similar results to healthy rats and given the low CNS permeability and high background signal from surrounding muscle and the nearby bladder, no measurable changes at the ligation sites could be detected (**Fig. S4**, Supporting Information). Therefore, biodistribution and *ex vivo* autoradiography experiments were carried out to monitor tracer binding in SNL rat brains, spinal cords and spinal nerves. The rats were euthanized 75 min after the tracer injection. Selected tissues and the L4-L6 spinal nerves on both the ligated side and unligated side were collected, weighed, and measured in a gamma counter for activity. Brain, thoracic spinal cord (tSpC), lumbar spinal cord (lSpC), and control (unligated) L4-L6 spinal nerves showed similar SUVs as healthy rats (**Fig 4A**). Compared to control nerves, [^18^F]tGBP4F showed around a 2-fold higher uptake in ligated nerves (205 ± 51%, n = 5, **Fig. 4C**). Similarly, [^18^F]cGBP4F showed around a 1.6-fold higher uptake in ligated *vs*. contralateral control nerves (166 ± 22 %, n = 4, **Fig. 4E**). Consistent results were obtained for both tracers when the activity was measured using *ex vivo* autoradiography instead of gamma counting (**Fig. 4B and 4D**). Importantly, the SUV of ligated nerves was around 11% higher than the surrounding muscle for [^18^F]cGBP4F and 20% higher for [^18^F]tGBP4F suggesting that in larger animals and humans, where the size of the spinal cord and the spinal nerves is larger than the resolution of PET, these tracers may serve to detect injured nerves.

**Figure 4.**
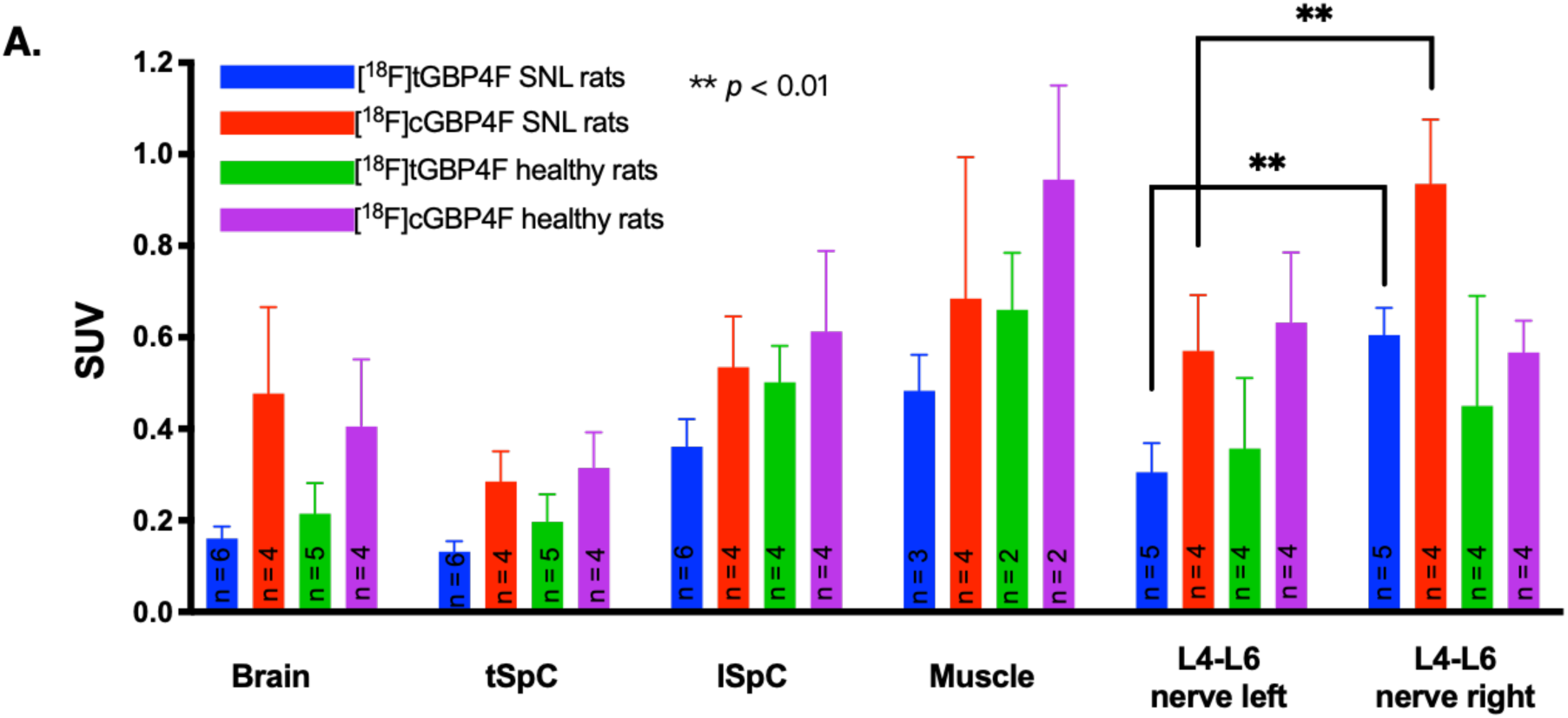
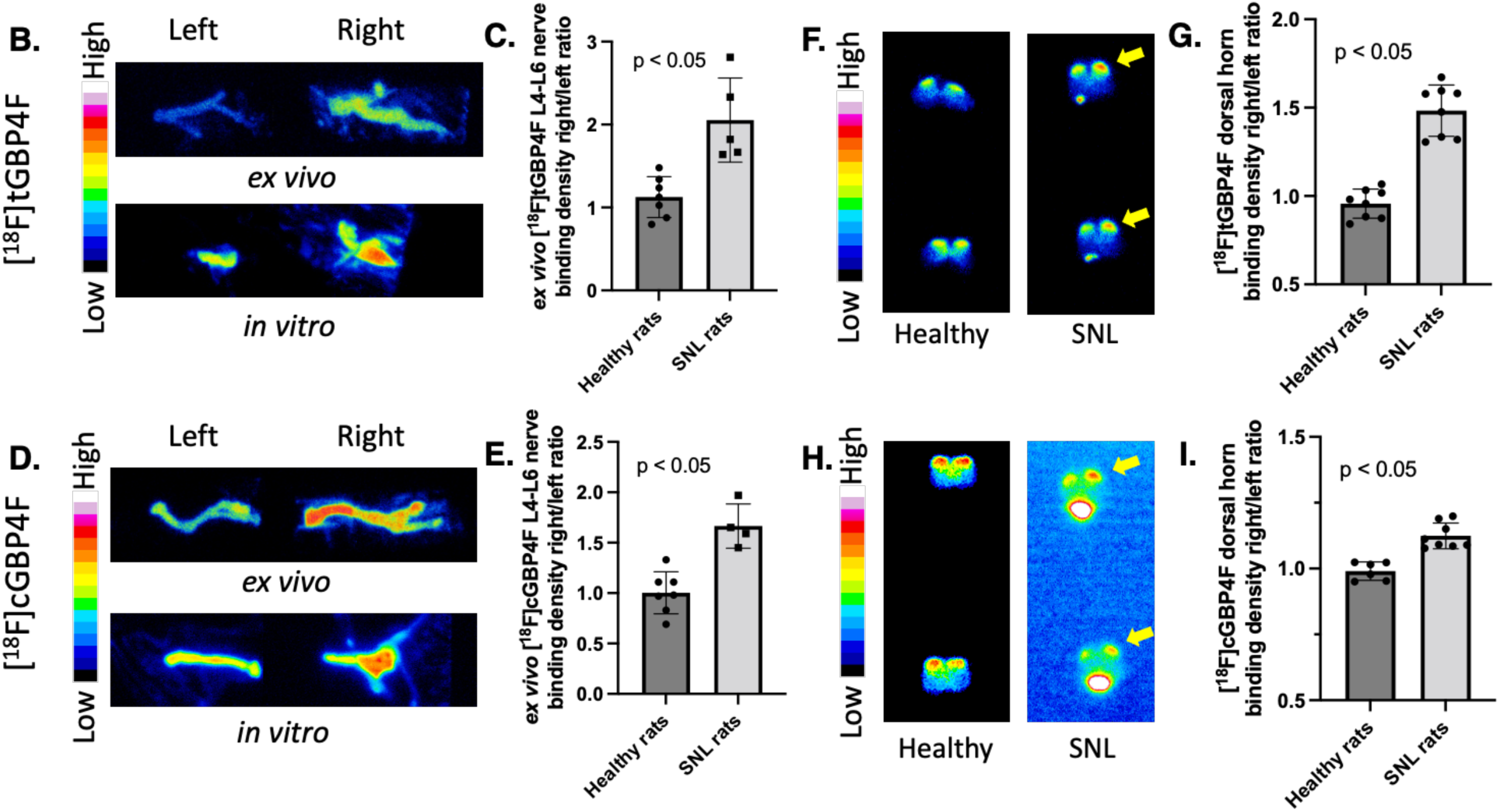
**A:** Biodistribution of [^18^F]tGBP4F and [^18^F]cGBP4F in select organs/tissues of SNL rats at 75 min post tracer injection (data obtained by the gamma counting of the dissected organs/tissues). **B:** Representative *ex vivo* and *in vitro* autoradiography of [^18^F]tGBP4F with SNL rat spinal nerves (10-15 days post-surgery) indicates an increase of [^18^F]tGBP4F accumulation to ligated nerves (right nerve) compared to non-ligated nerves (left nerve). **C:** The gamma counting shows 205 ± 51% increase of [^18^F]tGBP4F accumulation of SNL nerves (right nerves) compared to healthy nerves (left nerves). **D:** Representative *ex vivo* and *in vitro* autoradiography of [^18^F]cGBP4F with SNL rat nerves indicates an increase of [^18^F]cGBP4F accumulation to ligated nerves (right nerves) compared to non-ligated nerves (left nerves). **E:** The gamma counting shows 166 ± 22% increase of [^18^F]cGBP4F binding to SNL nerves (right nerves) compared to healthy nerves (left nerves).). **F:** Representative *in vitro* autoradiography of [^18^F]tGBP4F with SNL and healthy rat spinal cord slices. **G:** Quantitative autoradiography data shows 148 ± 15% (n = 8) increase of [^18^F]tGBP4F accumulation in unilateral side of dorsal horn of SNL rat spinal cord section. **H:** Representative *in vitro* autoradiography of [^18^F]tGBP4F with SNL and healthy rat spinal cord slices. **I:** Quantitative autoradiography data shows 112 ± 5% (n = 8) increase of [^18^F]tGBP4F accumulation in unilateral side of dorsal horn of SNL rat spinal cord section.

*In vitro* autoradiography with both tracers in explanted nerves also showed 1.5-2 fold higher binding in the injured than control nerves (**Fig. 4B and 4D**). This finding confirms that the increased uptake measured *ex vivo* is due to higher binding and not due to greater blood flow. Overall, the increase binding of the tracers in injured nerves is consistent with the literature on increased expression of the α2δ-1 receptor in spinal nerve ligation animals^31, 33, 41, 50-54^.

In addition, *in vitro* quantitative autoradiography with both tracers in spinal cord sections of SNL rats showed higher binding in the dorsal horn ipsilateral to the injured nerve (**Fig. 4F-4I**). These observations are supported by the immunohistochemistry staining (**Fig S5**, Supporting Information) and consistent with the overexpression of α2δ-1 receptors throughout the nerve as well as the ipsilateral dorsal root ganglia (DRG) and spinal cord of the SNL model as previously reported.^29, 46, 51^

## CONCLUSION

In conclusion, two novel fluorine derivatives of gabapentin, namely *trans*-4-fluorogabapentin (tGBP4F) and *cis*-4-fluorogabapentin (cGBP4F), were synthesized and characterized. Both compounds bind to the α2δ-1 receptor. tGBP4F shows a slightly higher IC_50_ compared to gabapentin suggesting it also has potential as a therapeutic. Fluorine-18 versions, *trans*-[^18^F]4-fluorogabapentin ([^18^F]tGBP4F) and *cis*-[^18^F]4-fluorogabapentin ([^18^F]cGBP4F) were also developed for α2δ-1 receptor scanning. Although the chemical structures of the two isomers, [^18^F]tGBP4F and [^18^F]cGBP4P are highly similar, their performance is significantly different (**Table 1**). In terms of similarities, both tracers show specific binding to α2δ-1 receptor *in vitro* as evidenced by the fact that both can be displaced by non-radioactive GBP (**Fig 2**). In terms of differences, the radiolabeling yield of [^18^F]tGBP4F is almost 3-fold higher than [^18^F]cGBP4F. In addition, [^18^F]tGBP4F shows about 100-fold higher affinity towards α2δ-1 receptor than [^18^F]cGBP4F (even higher than gabapentin) (**Fig 2**). Furthermore, *in vivo* evaluation shows that [^18^F]tGBP4F has a faster clearance than [^18^F]cGBP4F, which is generally advantageous for PET tracers (**Fig 3**). Moreover, both tracers show increased binding in injured nerves with [^18^F]tGBP4F being higher than [^18^F]cGBP4F when measured by *ex vivo* gamma counting and *ex vivo* autoradiography (**Fig 4**). Due to the small size of rat spinal nerves, and the relatively high signal in surrounding tissues, these changes were not observable by PET. However, we speculate that those changes might be observable in larger animals and humans.

**Table 1.** Comparison of [^18^F]tGBP4F) and [^18^F]cGBP4F.

| | Radiosynthesis | $\alpha 2\delta$-1 affinity | Pharmacokinetics | CNS permeability |
| --- | --- | --- | --- | --- |
| [<sup>18</sup>F]tGBP4F | high yield | high | slow | moderate-low |
| [<sup>18</sup>F]cGBP4F | low yield | low | slower | moderate |

Overall, these tracers provide novel useful neuroimaging tools for studying/detecting neuropathic pain in the laboratory. Future efforts to develop tracers for α2δ-1 receptor should focus on tracers with high nerve uptake and low background signal. In many chronic pain patients, it is not possible to identify if their pain is neuropathic in origin and these tools may assist in identifying such a component, whether it changes over time and may contribute to selection of the most appropriate therapeutic intervention.

## EXPERIMENTAL SECTION

### Chemistry and radiochemistry

#### Materials and characterization

The NMR spectra were recorded on Bruker spectrometers AV300 (^1^H, 300 MHz; ^13^C, 75 MHz) referenced to residual solvent signals as internal standards (^1^H NMR: CDCl_3_ 7.26 ppm and ^13^C NMR: CDCl_3_ 77.2 ppm). Concentrated solutions of samples in deuterated solvent were sealed off in a NMR tube for measurements. The HR-MS analysis was carried out on a Thermo Scientific Dionex UltiMate 3000 UHPLC coupled to a Thermo Q Exactive Plus mass spectrometer system (Thermo Fisher Scientific Inc, Waltham, MA) equipped with a HESI-II electrospray ionization (ESI) source. Commercially available reagents were purchased from SIGMA-Aldrich, Acros, Alfa-Assar or TCI and used as received. All compounds are >95% pure by HPLC analysis. Tritiated gabapentin (1-(Amino-[3H]-methyl)-[2,3,5,6-^3^H]-cyclohexane acetic acid) with molar activity of 90-120 Ci/mmol was purchased from American Radiolabeled Chemicals.

**2:** To a 30 mL vial containing solid **1** (167 mg, 1 mmol), di-tert-butyl dicarbonate (Boc_2_O, 260 mg, 1.2 mmol), and 4-dimethylaminopyridine (DMAP, 25 mg, 0.2 mmol), dichloromethane (DCM, 10 mL) and triethylamine (NEt3, 0.1 mL) were added via syringe. The mixture was stirred at room temperature for 2 hours. All volatiles were removed under reduced pressure. Residue was then washed with water and then extracted with DCM. Desired product (370 mg, 73% yield) was obtained after flash column as colorless oil. ^1^H NMR (300 MHz, CDCl_3_, 298 K): δ(ppm) = 1.55 (s, 9 H), 1.95 (t, *J* = 6.7 Hz, 4 H), 2.38−2.43 (m, 4 H), 2.58 (s, 2 H), 3.68 (s, 2 H). ^13^C NMR (75 MHz, CDCl_3_, 298 K): δ(ppm) = 28.01, 34.24, 35.65, 37.67, 44.07, 55.96, 83.40, 169.28, 171.93, 209.00. HR-MS:268.1543 for C_14_H_21_NO_4_ + H^+^ ([M]+H, theoretical calcd: 268.1541).

**3:** To a 30 mL vial containing **2** (360 mg, 1.4 mmol) in 10 mL of methanol, NaBH_4_ (120 mg, 3.2 mmol) was added in two portions. After stirring at room temperature for 2 hours, 1 mL of saturated NH_4_Cl solution was added into the reaction mixture. Volatiles were then removed under reduced pressure. Residue was extracted with DCM and gave the desired product (350 mg, 97% yield) as colorless oil. ^1^H NMR (300 MHz, CDCl_3_, 298 K): δ(ppm) = 1.42−1.48 (m, 4 H), 1.52−1.53 (s × 2, two isomers, 9 H), 1.73−1.87 (m, 4 H), 2.36−2.43 (s × 2, 2 H), 3.47−3.55 (s × 2, 2 H), 3.48 (s, 1 H), 3.71−3.76 (m, 1 H). ^13^C NMR (75 MHz, CDCl_3_, 298 K): δ(ppm) = 173.08, 172.95, 150.21, 82.94, 68.89, 68.28, 57.55, 55.84, 50.82, 45.76, 44.11, 34.24, 34.00, 33.29, 32.84, 31.20, 30.94, 28.44, 28.00.

**3a:** Compound **3** was dissolved in acetonitrile/water (1:1) mix solution and then purified by semi-prep HPLC giving colorless oil of the desired product. ^1^H NMR (300 MHz, CDCl3, 298 K): δ(ppm) = 1.3−1.87 (m, 17 H), 2.42 (s, 2 H), 3.47 (s, 2 H), 3.65−3.75 (m, 1 H). ^13^C NMR (75 MHz, CDCl_3_, 298 K): δ(ppm) = 173.15, 150.40, 83.12, 69.07, 57.75, 44.30, 34.43, 33.48, 31.37, 28.61, 28.18. HR-MS:292.1518 for C_14_H_23_NO_4_ + Na^+^ ([M]+Na, theoretical calcd: 292.1519).

**3b:** Compound **3** was dissolved in acetonitrile/water (1:1) mix solution and then purified by semi-prep HPLC giving colorless oil of the desired product. ^1^H NMR (300 MHz, CDCl3, 298 K): δ(ppm) = 1.33−1.97 (m, 17 H), 2.36 (s, 2 H), 3.55 (s, 2 H), 3.72−3.78 (m, 1 H). ^13^C NMR (75 MHz, CDCl_3_, 298 K): δ(ppm) = 173.28, 150.39, 83.15, 68.47, 56.02, 45.96, 34.18, 33.04, 31.13, 28.63, 28.20. HR-MS: 292.1517 for C_14_H_23_NO_4_ + Na^+^ ([M]+Na, theoretical calcd: 292.1519).

**4a:** To a 30 mL vial containing **3a** (25 mg, 0.1 mmol) in dichloromethane (DCM, 10 mL), diethylaminosulfur trifluoride (DAST, 40 mg, 0.25 mmol) was added via syringe at 0 °C dropwise, forming a yellow solution after stirring for 0.5 hour. Volatiles were removed under reduced pressure. Residue was dissolved in acetonitrile/water (1:1) mix solution and then purified by semi-prep HPLC affording colorless oil of the desired product (2 mg, 10% yield). ^1^H NMR (300 MHz, CDCl_3_, 298 K): δ(ppm) = 1.46−1.53 (m, 11 H), 1.66−1.90 (m, 6 H), 2.39 (s, 2 H), 3.52 (s, 2 H), 4.59−4.80 (m, 1 H). ^13^C NMR (75 MHz, CDCl_3_, 298 K): δ(ppm) = 172.81, 150.12, 89.87, 87.61, 83.01, 57.38, 43.96, 34.08, 34.07, 31.19, 31.12, 28.43, 28.02, 28.00, 27.75. ^19^F NMR (282 MHz, CDCl_3_, 298 K): δ(ppm) = −176.73. HR-MS: 294.1474 for C_14_H_22_FNO_4_ + Na^+^ ([M]+Na, theoretical calcd: 294.1476).

**4b:** To a 30 mL vial containing **3b** (25 mg, 0.1 mmol) in dichloromethane (DCM, 10 mL), diethylaminosulfur trifluoride (DAST, 40 mg, 0.25 mmol) was added via syringe at 0 °C dropwise, forming a yellow solution after stirring for 0.5 hour. Volatiles were removed under reduced pressure. Residue was dissolved in acetonitrile/water (1:1) mix solution and then purified by semi-prep HPLC affording colorless oil of the desired product (2 mg, 10% yield). ^1^H NMR (300 MHz, CDCl_3_, 298 K): δ(ppm) = 1.47−1.58 (m, 11 H), 1.73−1.88 (m, 6 H), 2.41 (s, 2 H), 3.52 (s, 2 H), 4.55−4.78 (m, 1 H). ^13^C NMR (75 MHz, CDCl_3_, 298 K): δ(ppm) = 172.82, 150.46, 90.46, 88.19, 83.24, 56.05, 45.73, 34.18, 34.17, 31.80, 31.71, 28.36, 28.20, 28.09. ^19^F NMR (282 MHz, CDCl_3_, 298 K): δ(ppm) = −176.34. HR-MS: 294.1474 for C_14_H_22_FNO_4_ + Na^+^ ([M]+Na, theoretical calcd: 294.1476).

**5a** (tGBP4F): To a 30 mL vial containing **4a** (2 mg, 0.01 mmol) in tetrahydrofuran (THF, 1 mL), 0.1 mL of saturated LiOH solution was added via syringe dropwise. After stirring at room temperature for 3 hours, excess LiOH was quenched by adding 1 mL of saturated NH_4_Cl solution into the reaction mixture. Intermediate was obtained by extracting the mixture using ethyl acetate. Intermediate was then transferred into a new 30 mL vial and 1 mL of trifluoroacetic acid (TFA) was added by syringe. After stirring at room temperature for 0.5 hour, clean solution was transfer into another 30 mL vial. Volatiles were removed under reduced pressure. Residue was extracted with H_2_O and then purified by semi-prep HPLC giving desired product. ^1^H NMR (300 MHz, D_2_O, 298 K): δ(ppm) = 1.46−1.54 (m, 2 H), 1.59−1.68 (m, 2 H), 1.79−1.92 (m, 4 H), 2.46 (s, 2 H), 3.05 (s, 2 H), 4.71−5.00 (m, 1 H). ^13^C NMR (75 MHz, D_2_O, 298 K): δ(ppm) = 180.26, 91.95, 89.78, 48.46, 44.03, 33.56, 28.18, 28.12, 26.10, 25.83. ^19^F NMR (282 MHz, D_2_O, 298 K): δ(ppm) = −174.80. HR-MS: 190.1236 for C_9_H_17_FNO_4_ + H^+^ ([M]+H, theoretical calcd: 190.1238).

**5b** (cGBP4F): To a 30 mL vial containing **4b** (2 mg, 0.01 mmol) in tetrahydrofuran (THF, 1 mL), 0.1 mL of saturated LiOH solution was added via syringe dropwise. After stirring at room temperature for 3 hours, excess LiOH was quenched by adding 1 mL of saturated NH_4_Cl solution into the reaction mixture. Intermediate was obtained by extracting the mixture using ethyl acetate. Intermediate was then transferred into a new 30 mL vial and 1 mL of trifluoroacetic acid (TFA) was added by syringe. After stirring at room temperature for 0.5 hour, clean solution was transfer into another 30 mL vial. Volatiles were removed under reduced pressure. Residue was extracted with H_2_O and then purified by semi-prep HPLC giving desired product. ^1^H NMR (300 MHz, D_2_O, 298 K): δ(ppm) = 1.39−1.48 (m, 2 H), 1.69−1.96 (m, 6 H), 2.48 (s, 2 H), 3.04 (s, 2 H), 4.66−4.90 (m, 1 H). ^13^C NMR (75 MHz, D_2_O, 298 K): δ(ppm) = 180.11, 92.79, 90.61, 47.24, 45.63, 33.53, 29.30, 29.20, 26.51, 26.25. ^19^F NMR (282 MHz, D_2_O, 298 K): δ(ppm) = −171.12. HR-MS: 190.1236 for C_9_H_17_FNO_4_ + H^+^ ([M]+H, theoretical calcd: 190.1238).

**6a:** To a 30 mL vial containing **3a** (25 mg, 0.1 mmol) and triethylamine (NEt_3_, 0.05 mL) in dichloromethane (DCM, 10 mL), methanesulfonyl chloride (MsCl, 14 mg, 0.13 mmol) was added via syringe dropwise. The mixture was stirred at room temperature for 6 hours. All volatiles were removed under reduced pressure. Residue was then extracted with DCM. Desired product (27 mg, 72% yield) was obtained after flash column as colorless oil. ^1^H NMR (300 MHz, CDCl_3_, 298 K): δ(ppm) = 1.47−1.57 (m, 11 H), 1.73−1.84 (m, 4 H), 1.93−2.01 (m, 2 H), 2.43 (s, 2 H), 3.02 (s, 3 H), 3.50 (s, 2 H), 4.77 (tt, *J*_*1*_ = 4 Hz, *J*_*2*_ = 8 Hz, 1 H). ^13^C NMR (75 MHz, CDCl_3_, 298 K): δ(ppm) = 172.31, 150.12, 83.19, 78.47, 38.78, 33.85, 32.35, 28.57, 28.01, 28.00. HR-MS: 370.1291 for C_15_H_25_NO_6_S + Na^+^ ([M]+Na, theoretical calcd: 370.1295).

**6b:** To a 30 mL vial containing **3b** (25 mg, 0.1 mmol) and triethylamine (NEt_3_, 0.05 mL) in dichloromethane (DCM, 10 mL), methanesulfonyl chloride (MsCl, 14 mg, 0.13 mmol) was added via syringe dropwise. The mixture was stirred at room temperature for 6 hours. All volatiles were removed under reduced pressure. Residue was then extracted with DCM. Desired product (25 mg, 71% yield) was obtained after flash column as colorless oil. ^1^H NMR (300 MHz, CDCl_3_, 298 K): δ(ppm) = 1.48−1.57 (m, 11 H), 1.74−1.91 (m, 6 H), 2.39 (s, 2 H), 3.02 (s, 3 H), 3.55 (s, 2 H), 4.80 (tt, *J*_*1*_ = 5 Hz, *J*_*2*_ = 5 Hz, 1 H). ^13^C NMR (75 MHz, CDCl_3_, 298 K): δ(ppm) = 172.61, 150.26, 83.35, 78.52, 56.69, 44.82, 38.90, 34.02, 31.99, 28.59, 28.20. HR-MS: 370.1292 for C_15_H_25_NO_6_S + Na^+^ ([M]+Na, theoretical calcd: 370.1295).

**8a** ([^18^F]tGBP4F) and **8b** ([^18^F]cGBP4F): [^18^F]KF is made by bombardment of a H_2_^18^O with a 16.5 MeV proton beam in a GE PETtrace cyclotron (GE Healthcare, Waukesha, WI, USA) using the ^18^O(p, n)^18^F nuclear reaction. [^18^F]KF is loaded onto a QMA cartridge and then eluted with kryptofix-2.2.2.(K_222_)/K_2_CO_3_ water/acetonitrile solution (1:1). The mixture is dried under nitrogen flow and then ∼0.5 mg of precursor **6a** or **6b** dissolved in 0.3 mL DMSO is added. The ^18^F labeled Boc protected gabapentin lactam (**7a** or **7b**) is obtained by the reaction [^18^F]KF and **6a** or **6b** in the presence of K_222_/K_2_CO_3_ at 140°C for 20 min. ^18^F-labeled gabapentin **8a** ([^18^F]tGBP4F) or **8b** ([^18^F]cGBP4F) is obtained by the hydrolysis and deprotection of **7a** or **7b** using NaOH (2N) and HCl (3N), respectively. Reversed-phase HPLC is used for the purification of **8a** ([^18^F]tGBP4F) or **8b** ([^18^F]cGBP4F) using XBridge 5um C_18_ semiprep-column (10 × 250 mm). The mobile phase is NaH_2_PO_4_ EtOH-H_2_O (10mM, v/v = 5:95) solution at flow rate of 4 mL/min. All the radiosynthesis were carried out using GE TRACERLab FX2N radiosynthesis platform (GE Healthcare, Waukesha, WI, USA) in hot cell.

### Autoradiography

#### [^3^H]gabapentin autoradiography

Rat brains and mouse spinal cords from spared nerve injury (SNI) mice were used for tritium autoradiography, 20μm thick sections were incubated with 10 nCi of [^3^H]gabapentin in PBS for 30 min in the presence or absence of cold competitors (5 μM gabapentin, pregabalin) and then washed with ice cold PBS (3x 1 min). The slides were dried under a stream of nitrogen and apposed onto a tritium sensitive imaging plate (^3^H-Fujifilm imaging plates; Bas-TR2025, Fuji Photo Film, Japan) for 14 days in the dark at - 20 °C. Afterwards, the plate was digitized using a Typhoon 9000 phosphorimager. The tritium autoradiography data were analyzed by measuring the mean pixel intensity of the region of interest using the freehand selection tool in ImageJ software.

#### *In vitro* Autoradiography

20 μm thick rat spinal cord sections were incubated with 4-5 μCi of [^18^F]tGBP4F and [^18^F]cGBP4F in PBS for 30 min in the presence or absence nonradioactive competitors (0-5 μM of gabapentin (GBP), tGBP4F, and cGBP4F) and then washed with cold PBS (3 × 1 min). The slides were air dried and exposed onto an imaging screen. The screen was digitized using a Typhoon 9000 phosphorimager after 16 h of exposure. The autoradiography data were processed analyzed by ImageJ software. Dorsal horn areas were selected as region of interests (ROIs) and ROIs were draw manually and mean pixel intensities were obtained from ImageJ software. For the competition binding experiment, the inhibition rate of each nonradioactive competitors was calculated accordingly and loaded to Prism software for IC_50_ calculation.

#### *Ex vivo* autoradiography of spinal nerve

Tracer injected rats were euthanized and dissected. The L4-L6 spinal nerves were collected and placed on a plastic plate. The plate was exposed on imaging screen for 16 h at -20 °C. The screen was then digitized using a Typhoon 9000 phosphorimager. The autoradiography data were processed analyzed by ImageJ software.

### Immunohistochemistry

Cryosections were air dried and fixed in neutral formalin for 10 min at room temperature (RT) then rinsed in phosphate buffered saline (PBS). Antigen retrieval was performed by incubating the sections in 20 mM Tris, pH 8.0, 10 mM EDTA at 95°C for 1 h. The sections were blocked with 0.3% H_2_O_2_ in water for 15 min, then rinsed in PBS, then blocked with PBS, 0.1% BSA, 0.05% Triton X-114 for 1 h at RT, followed by incubation with mouse anti-α2δ-1 antibody (Clone 20A, Sigma, St Louis, MO) at 1:50 for 1 h. After rinsing in PBS, the sections were incubated with a biotinylated goat-anti-mouse IgG (Jackson Immuno Research, West Grove, PA) at 1:100 for 1 h at RT followed by Vectastain elite ABC-HEP (Vector Labs, Burlingame, CA) for 30 min. The reaction was visualized by incubating with DAB chromogen. The reaction was stopped by water, air dried, and then mounted with permount.

### Animal experiments

#### Compliance

Animal experiments were approved by the Animal Care and Use Committee at the Massachusetts General Hospital (MGH) and Boston Children’s Hospital (BCH).

#### Mice

Male, 6-8 weeks old C57Bl6J mice (Jackson’s Laboratories, Maine, USA) were used for preparing the spared nerve injury (SNI) mice, which was done by lesioning the tibial and common peroneal nerve branches.^55, 56^ The spinal cord tissue was collected and cryosectioned for tritium autoradiography experiment.

#### Rats

6-8 weeks old Sprague-Dawley male and female healthy rats (Charles River, USA) were used in this study. 7 Weeks old male Sprague Dawley rats were used for preparing the rat model of neuropathic pain (spinal nerve ligation rats), which were obtained from Charles River through the preconditioned animal model service. Spinal nerve ligation was done by are tightly ligating the right L5 and L6 spinal nerves with a silk ligature.^49^ The SNL rats were held at Charles River for 7 days post-surgery before shipping to the MGH animal facility. The spinal nerve ligation rats were kept at the MGH animal facility for an additional one to two weeks before the PET imaging and biodistribution study.

#### microPET/CT imaging of rats

Rats were scanned on a SuperArgus 2R microPET/CT scanner (SEDECAL S.A., Madrid, Spain). The scanner works in a 3D list mode and features a 60 mm radial field of view (FOV) and 47 mm axial FOV. The image resolution was 1.1 mm. The energy window was set to 250-700 keV and the timing window was 1.6 ns. Rats were tail vein-injected a corresponding radiotracer (100-300 μCi in 0.2-0.4 mL of saline) with or without Intraperitoneal pre-injection of gabapentin (30 mg/kg) under anesthesia (2% isoflurane with oxygen flow of 1.5L/min). Each rat underwent a whole-body (consisting of 3 or 4 bed positions depending on animal size) dynamic scan for 60 minutes. To this end, each bed position was sequentially scanned for 5 min, and the whole-body passages were repeated 3 or 4 times. The images of adjacent bed positions were stitched together automatically using a 6-slices overlap and the whole-body data sets were stacked into temporal frames. 15-minutes CT scan was acquired after each PET imaging acquisition. The CT acquisition settings were: tube voltage: 45 kV, tube current: 250 microA, resolution: standard, number of shots: 2. CT images were automatically registered with PET images and were used both for anatomical reference and attenuation correction of the PET images. Besides tissue attenuation, corrections for isotope decay, detectors dead-time, and random coincidences were applied during raw data histogramming and image reconstruction. The data were reconstructed into the 175×175 pixels matrix with the pixel size of 0.39×0.39 mm^2^ and the slice thickness of 0.78 mm. 2-Dimensional ordered-subset expectation maximization (OSEM2D) algorithm with 2 iterations and 16 subsets was utilized for image reconstruction. Rat PET imaging data were analyzed by AMIDE (Amide’s a medical imaging data examiner) software (open-source software, Los Angeles, CA, USA).^57^ Volumes of interests (VOIs) were drawn manually under the guide of CT and PET images.

#### Biodistribution study of rats

Corresponding radiotracer (100-300 μCi in 0.2-0.4 mL of saline) was administrated *via* tail vein injection under anesthesia (2% isoflurane with oxygen flow of 1.5L/min). The animals were kept under anesthesia (2% isoflurane with oxygen flow of 1.5L/min) for 75 min before euthanasia by intraperitoneal injection of 0.2 mL of euthasol solution. The blood was collected by cardiac puncture and other tissues were collected by dissection. Tissue and blood samples were weighed, and the radioactive concentrations were measured using a Wallac Wizard gamma counter. The biodistributions at different tissue were expressed as standard uptake value (SUV).

#### SUV calculation

SUV of selected organ or tissue was calculated following the equation 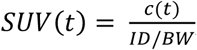, where SUV is the standardized uptake value at time *t*, c(t) is the radioactivity concentration (measured by PET imaging or by gamma counting) of selected organ corrected to *t* = 0, ID is the injected dose corrected to *t* = 0, BW is the body weight of the animal.

## ACKNOWLEDGEMETNS

We thank David Lee and Timothy Beaudoin at the MGH Gordon PET cyclotron facility for producing fluorine-18. We thank Jin Hong for 2-D NMR measurement and data analysis. We thank Jennifer X. Wang for high resolution mass spectra measurement and data analysis.

## AUTHOR CONTRIBUTIONS

YPZ contributed to the study design, developed the synthesis method and synthesized the radiotracers, performed the autoradiography and animal dissection, assisted with the rats PET imaging and biodistribution experiments, processed and analyzed the autoradiography and rat PET imaging data; YS performed the gamma counting and processed and analyzed the biodistribution data, assisted with the rats PET imaging. KT prepared the rats for PET imaging and biodistribution experiments, assisted with the rats PET imaging, and performed the immunohistochemistry experiment. VB performed the rat PET imaging experiments; NA dissected the spinal cord tissues from the SNI mice; CJW contributed to the study design and supervised the SNI mice experiment; PB contributed to the study design, performed the tritium autoradiography, and supervised the entire project. YPZ, and PB wrote the manuscript and all authors reviewed and approved it.

## REFERENCES

1. Jensen, T. S.; Baron, R.; Haanpää, M.; Kalso, E.; Loeser, J. D.; Rice, A. S. C.; Treede, R.-D., A new definition of neuropathic pain. PAIN 2011, 152 (10), 2204–2205.

2. van Hecke, O.; Austin, S. K.; Khan, R. A.; Smith, B. H.; Torrance, N., Neuropathic pain in the general population: A systematic review of epidemiological studies. PAIN 2014, 155 (4), 654–662.

3. Davis, K. D.; Aghaeepour, N.; Ahn, A. H.; Angst, M. S.; Borsook, D.; Brenton, A.; Burczynski, M. E.; Crean, C.; Edwards, R.; Gaudilliere, B.; Hergenroeder, G. W.; Iadarola, M. J.; Iyengar, S.; Jiang, Y.; Kong, J.-T.; Mackey, S.; Saab, C. Y.; Sang, C. N.; Scholz, J.; Segerdahl, M.; Tracey, I.; Veasley, C.; Wang, J.; Wager, T. D.; Wasan, A. D.; Pelleymounter, M. A., Discovery and validation of biomarkers to aid the development of safe and effective pain therapeutics: challenges and opportunities. Nat. Rev. Neurol. 2020, 16 (7), 381–400.

4. Mogil, J. S., Animal models of pain: progress and challenges. Nat. Rev. Neurosci.2009, 10 (4), 283–294.

5. Turner, P. V.; Pang, D. S. J.; Lofgren, J. L. S., A Review of Pain Assessment Methods in Laboratory Rodents. Comp. Med. 2019, 69 (6), 451–467.

6. Fisher, A. S.; Lanigan, M. T.; Upton, N.; Lione, L. A., Preclinical Neuropathic Pain Assessment; the Importance of Translatability and Bidirectional Research. Front. Pharmacol. 2021, 11 (2308).

7. Agha, H.; McCurdy, C. R., In vitro and in vivo sigma 1 receptor imaging studies in different disease states. RSC Med. Chem. 2021, 12 (2), 154–177.

8. Da Silva, J. T.; Seminowicz, D. A., Neuroimaging of pain in animal models: a review of recent literature. PAIN Reports 2019, 4 (4), e732.

9. Morton, D. L.; Sandhu, J. S.; Jones, A. K., Brain imaging of pain: state of the art. J Pain Res 2016, 9, 613–624.

10. Woolf, C. J.; Mannion, R. J., Neuropathic pain: aetiology, symptoms, mechanisms, and management. The Lancet 1999, 353 (9168), 1959–1964.

11. Woolf, C. J., Capturing Novel Non-opioid Pain Targets. Biol.l Psychiatry 2020, 87 (1), 74–81.

12. Yekkirala, A. S.; Roberson, D. P.; Bean, B. P.; Woolf, C. J., Breaking barriers to novel analgesic drug development. Nat.Rev. Drug Discov. 2017, 16 (8), 545–564.

13. Piel, M.; Vernaleken, I.; Rösch, F., Positron Emission Tomography in CNS Drug Discovery and Drug Monitoring. J. Med. Chem. 2014, 57 (22), 9232–9258.

14. Kim, J.; Ryu, S. B.; Lee, S. E.; Shin, J.; Jung, H. H.; Kim, S. J.; Kim, K. H.; Chang, J. W., Motor cortex stimulation and neuropathic pain: how does motor cortex stimulation affect pain-signaling pathways? J. Neurosurg. 2016, 124 (3), 866–876.

15. Jones, A. K. P.; Brown, W. D.; Friston, K. J.; Qi, L. Y.; Frackowiak, R. S. J., Cortical and subcortical localization of response to pain in man using positron emission tomography. Proc. Royal So. B. 1991, 244 (1309), 39–44.

16. Phelps, M. E., Positron computed tomography studies of cerebral glucose metabolism in man: Theory and application in nuclear medicine. Semin. Nucl. Med. 1981, 11 (1), 32–49.

17. Thompson, S. J.; Pitcher, M. H.; Stone, L. S.; Tarum, F.; Niu, G.; Chen, X.; Kiesewetter, D. O.; Schweinhardt, P.; Bushnell, M. C., Chronic neuropathic pain reduces opioid receptor availability with associated anhedonia in rat. PAIN 2018, 159 (9), 1856–1866.

18. Willoch, F.; Schindler, F.; Wester, H. J.; Empl, M.; Straube, A.; Schwaiger, M.; Conrad, B.; Tölle, T. R., Central poststroke pain and reduced opioid receptor binding within pain processing circuitries: a [11C]diprenorphine PET study. PAIN 2004, 108 (3), 213–220.

19. Jones, A. K. P.; Watabe, H.; Cunningham, V. J.; Jones, T., Cerebral decreases in opioid receptor binding in patients with central neuropathic pain measured by [11C]diprenorphine binding and PET. Eur. J. Pain 2004, 8 (5), 479–485.

20. Wood, P. B.; Patterson, J. C.; Sunderland, J. J.; Tainter, K. H.; Glabus, M. F.; Lilien, D. L., Reduced Presynaptic Dopamine Activity in Fibromyalgia Syndrome Demonstrated With Positron Emission Tomography: A Pilot Study. J. Pain 2007, 8 (1), 51–58.

21. Chung, G.; Kim, C. Y.; Yun, Y.-C.; Yoon, S. H.; Kim, M.-H.; Kim, Y. K.; Kim, S. J., Upregulation of prefrontal metabotropic glutamate receptor 5 mediates neuropathic pain and negative mood symptoms after spinal nerve injury in rats. Sci. Rep. 2017, 7 (1), 9743.

22. Yoon, D.; Cipriano, P.; Carroll, I.; Curtin, C.; Roh, E.; Wilson, T.; Biswal, S., Sigma-1 receptor PET/MRI for identifying nociceptive sources of radiating low back pain. J. Nucl. Med. 2020, 61 (supplement 1), 178–178.

23. Shen, B.; Behera, D.; James, M. L.; Reyes, S. T.; Andrews, L.; Cipriano, P. W.; Klukinov, M.; Lutz, A. B.; Mavlyutov, T.; Rosenberg, J.; Ruoho, A. E.; McCurdy, C. R.; Gambhir, S. S.; Yeomans, D. C.; Biswal, S.; Chin, F. T., Visualizing Nerve Injury in a Neuropathic Pain Model with [18F]FTC-146 PET/MRI. Theranostics 2017, 7 (11), 2794–2805.

24. James, M. L.; Shen, B.; Zavaleta, C. L.; Nielsen, C. H.; Mesangeau, C.; Vuppala, P. K.; Chan, C.; Avery, B. A.; Fishback, J. A.; Matsumoto, R. R.; Gambhir, S. S.; McCurdy, C. R.; Chin, F. T., New Positron Emission Tomography (PET) Radioligand for Imaging σ-1 Receptors in Living Subjects. J. Med. Chem. 2012, 55 (19), 8272–8282.

25. Kremer, M.; Salvat, E.; Muller, A.; Yalcin, I.; Barrot, M., Antidepressants and gabapentinoids in neuropathic pain: Mechanistic insights. Neuroscience 2016, 338, 183–206.

26. Dolphin, A. C., The α2δ subunits of voltage-gated calcium channels. Biochimi. Biophys. Acta 2013, 1828 (7), 1541–1549.

27. Luo, Z. D.; Chaplan, S. R.; Higuera, E. S.; Sorkin, L. S.; Stauderman, K. A.; Williams, M. E.; Yaksh, T. L., Upregulation of Dorsal Root Ganglion α2δ Calcium Channel Subunit and Its Correlation with Allodynia in Spinal Nerve-Injured Rats. J. Neurosci. 2001, 21 (6), 1868–1875.

28. Newton, R. A.; Bingham, S.; Case, P. C.; Sanger, G. J.; Lawson, S. N., Dorsal root ganglion neurons show increased expression of the calcium channel α2δ-1 subunit following partial sciatic nerve injury. Mol. Brain Res. 2001, 95 (1), 1–8.

29. Melrose, H. L.; Kinloch, R. A.; Cox, P. J.; Field, M. J.; Collins, D.; Williams, D., [3H]pregabalin binding is increased in ipsilateral dorsal horn following chronic constriction injury. Neurosci. Lett. 2007, 417 (2), 187–192.

30. Xiao, W.; Boroujerdi, A.; Bennett, G. J.; Luo, Z. D., Chemotherapy-evoked painful peripheral neuropathy: Analgesic effects of gabapentin and effects on expression of the alpha-2-delta type-1 calcium channel subunit. Neuroscience 2007, 144 (2), 714–720.

31. Boroujerdi, A.; Kim, H. K.; Lyu, Y. S.; Kim, D.-S.; Figueroa, K. W.; Chung, J. M.; Luo, Z. D., Injury discharges regulate calcium channel alpha-2-delta-1 subunit upregulation in the dorsal horn that contributes to initiation of neuropathic pain. PAIN 2008, 139 (2), 358–366.

32. Boroujerdi, A.; Zeng, J.; Sharp, K.; Kim, D.; Steward, O.; Luo, D. Z., Calcium channel alpha-2-delta-1 protein upregulation in dorsal spinal cord mediates spinal cord injury-induced neuropathic pain states. PAIN 2011, 152 (3), 649–655.

33. Nieto-Rostro, M.; Sandhu, G.; Bauer, C. S.; Jiruska, P.; Jefferys, J. G. R.; Dolphin, A. C., Altered expression of the voltage-gated calcium channel subunit α2δ-1: A comparison between two experimental models of epilepsy and a sensory nerve ligation model of neuropathic pain. Neuroscience 2014, 283, 124–137.

34. D’Arco, M.; Margas, W.; Cassidy, J. S.; Dolphin, A. C., The Upregulation of α2δ-1 Subunit Modulates Activity-Dependent Ca2+ Signals in Sensory Neurons. The Journal of Neuroscience 2015, 35 (15), 5891–5903.

35. Fornasari, D., Pharmacotherapy for Neuropathic Pain: A Review. Pain and Therapy 2017, 6 (1), 25–33.

36. Kusuyama, K.; Tachibana, T.; Yamanaka, H.; Okubo, M.; Yoshiya, S.; Noguchi, K., Upregulation of calcium channel alpha-2-delta-1 subunit in dorsal horn contributes to spinal cord injury-induced tactile allodynia. Spine J. 2018, 18 (6), 1062–1069.

37. Nieto-Rostro, M.; Ramgoolam, K.; Pratt, W. S.; Kulik, A.; Dolphin, A. C., Ablation of α<sub>2</sub>δ-1 inhibits cell-surface trafficking of endogenous N-type calcium channels in the pain pathway in vivo. PNAS. 2018, 115 (51), E12043–E12052.

38. Li, C. Y.; Zhang, X. L.; Matthews, E. A.; Li, K. W.; Kurwa, A.; Boroujerdi, A.; Gross, J.; Gold, M. S.; Dickenson, A. H.; Feng, G.; Luo, Z. D., Calcium channel alpha2delta1 subunit mediates spinal hyperexcitability in pain modulation. Pain 2006, 125 (1-2), 20–34.

39. Patel, R.; Bauer, C. S.; Nieto-Rostro, M.; Margas, W.; Ferron, L.; Chaggar, K.; Crews, K.; Ramirez, J. D.; Bennett, D. L. H.; Schwartz, A.; Dickenson, A. H.; Dolphin, A. C., α2δ-1 Gene Deletion Affects Somatosensory Neuron Function and Delays Mechanical Hypersensitivity in Response to Peripheral Nerve Damage. The Journal of Neuroscience 2013, 33 (42), 16412–16426.

40. Morimoto, S.-i.; Ito, M.; Oda, S.; Sugiyama, A.; Kuroda, M.; Adachi-Akahane, S., Spinal Mechanism Underlying the Antiallodynic Effect of Gabapentin Studied in the Mouse Spinal Nerve Ligation Model. J. Pharmacol. Sci. 2012, 118 (4), 455–466.

41. Bauer, C. S.; Nieto-Rostro, M.; Rahman, W.; Tran-Van-Minh, A.; Ferron, L.; Douglas, L.; Kadurin, I.; Sri Ranjan, Y.; Fernandez-Alacid, L.; Millar, N. S.; Dickenson, A. H.; Lujan, R.; Dolphin, A. C., The Increased Trafficking of the Calcium Channel Subunit α2δ-1 to Presynaptic Terminals in Neuropathic Pain Is Inhibited by the α2δ Ligand Pregabalin. J. Neurosci. 2009, 29 (13), 4076–4088.

42. Guo, X.; Zhu, H.; Yang, Z., Novel 64Cu-NOTA-1B50-1 targeting α2δ-1 for specific cancer stem cell imaging: A preliminary experiments on Hepatocellular Carcinoma. J. Nucl. Med. 2019, 60 (supplement 1), 610.

43. Bockbrader, H. N.; Wesche, D.; Miller, R.; Chapel, S.; Janiczek, N.; Burger, P., A Comparison of the Pharmacokinetics and Pharmacodynamics of Pregabalin and Gabapentin. Clin.Pharmacokinet. 2010, 49 (10), 661–669.

44. Bryans, J. S.; Davies, N.; Gee, N. S.; Dissanayake, V. U. K.; Ratcliffe, G. S.; Horwell, D. C.; Kneen, C. O.; Morrell, A. I.; Oles, R. J.; O’Toole, J. C.; Perkins, G. M.; Singh, L.; Suman-Chauhan, N.; O’Neill, J. A., Identification of Novel Ligands for the Gabapentin Binding Site on the α2δ Subunit of a Calcium Channel and Their Evaluation as Anticonvulsant Agents. J. Med. Chem. 1998, 41 (11), 1838–1845.

45. Cichon, J.; Sun, L.; Yang, G., Spared Nerve Injury Model of Neuropathic Pain in Mice. Bio-protocol 2018, 8 (6), e2777.

46. Taylor, C. P.; Garrido, R., Immunostaining of rat brain, spinal cord, sensory neurons and skeletal muscle for calcium channel alpha2-delta (α2-δ) type 1 protein. Neuroscience 2008, 155 (2), 510–521.

47. Bryans, J. S.; Davies, N.; Gee, N. S.; Dissanayake, V. U.; Ratcliffe, G. S.; Horwell, D. C.; Kneen, C. O.; Morrell, A. I.; Oles, R. J.; O’Toole, J. C.; Perkins, G. M.; Singh, L.; Suman-Chauhan, N.; O’Neill, J. A., Identification of novel ligands for the gabapentin binding site on the alpha2delta subunit of a calcium channel and their evaluation as anticonvulsant agents. J. Med. Chem. 1998, 41 (11), 1838–45.

48. Liu, W.; Huang, X.; Cheng, M.-J.; Nielsen, R. J.; Goddard, W. A.; Groves, J. T., Oxidative Aliphatic C-H Fluorination with Fluoride Ion Catalyzed by a Manganese Porphyrin. Science 2012, 337 (6100), 1322–1325.

49. Chung, J. M.; Kim, H. K.; Chung, K., Segmental Spinal Nerve Ligation Model of Neuropathic Pain. In Pain Research: Methods and Protocols, Luo, Z. D., Ed. Humana Press: Totowa, NJ, 2004; pp 35–45.

50. Nguyen, D.; Deng, P.; Matthews, E. A.; Kim, D.-S.; Feng, G.; Dickenson, A. H.; Xu, Z. C.; Luo, Z. D., Enhanced pre-synaptic glutamate release in deep-dorsal horn contributes to calcium channel alpha-2-delta-1 protein-mediated spinal sensitization and behavioral hypersensitivity. Mol. Pain 2009, 5 (1), 6.

51. Bauer, Claudia S.; Rahman, W.; Tran-Van-Minh, A.; Lujan, R.; Dickenson, Anthony H.; Dolphin, Annette C., The anti-allodynic α2δ ligand pregabalin inhibits the trafficking of the calcium channel α2δ-1 subunit to presynaptic terminals in vivo. Biochem. Soc. Trans. 2010, 38 (2), 525–528.

52. Zhou, C.; Luo, Z. D., Electrophysiological characterization of spinal neuron sensitization by elevated calcium channel alpha-2-delta-1 subunit protein. Eur. J. Pain 2014, 18 (5), 649–658.

53. Chang, E.; Chen, X.; Kim, M.; Gong, N.; Bhatia, S.; Luo, Z. D., Differential effects of voltage-gated calcium channel blockers on calcium channel alpha-2-delta-1 subunit protein-mediated nociception. Eur. J. Pain 2015, 19 (5), 639–648.

54. Zhou, C.; Luo, Z. D., Nerve injury-induced calcium channel alpha-2-delta-1 protein dysregulation leads to increased pre-synaptic excitatory input into deep dorsal horn neurons and neuropathic allodynia. Eur. J. Pain 2015, 19 (9), 1267–1276.

55. Bennett, G. J.; Xie, Y.-K., A peripheral mononeuropathy in rat that produces disorders of pain sensation like those seen in man. PAIN 1988, 33 (1), 87–107.

56. Ho Kim, S.; Mo Chung, J., An experimental model for peripheral neuropathy produced by segmental spinal nerve ligation in the rat. PAIN 1992, 50 (3), 355–363.

57. Loening, A. M.; Gambhir, S. S., AMIDE: A Free Software Tool for Multimodality Medical Image Analysis. Mol. Imag. 2003, 2 (3), 15353500200303133.

