## Supporting Information for "Development of a PET radioligand for α2δ-1 subunit of calcium channels for imaging neuropathic pain"

***Correspondence:**

Pedro Brugarolas, PhD

55 Fruit St

Bulfinch 051

Boston, MA 02114

orcid.org/0000-0002-7455-2743

### Contents


| Topics | Pages |
|---|---|
| **Fig S1.** HPLC chromatography of [$^{18}$F]GBP4F coinjection | 3 |
| **Fig S2.** Immunohistochemistry staining of α2δ-1receptor on spinal cord section of healthy rat and high resolution autoradiography of [$^{18}$F]tGBP4F and [$^{18}$F]cGBP4F in healthy rat spinal cord slices | 4 |
| **Fig S3.** Whole body PET imaging of healthy rats (sagittal view, summed images at 0−20 min, 20−40 min, and 40−60 min intervals) with [$^{18}$F]tGBP4F and [$^{18}$F]cGBP4F. | 5 |
| **Fig S4.** Whole body PET imaging of SNL rats (coronal view-left and sagittal view-right, summed images of 0−60 min intervals) with [$^{18}$F]tGBP4F. | 5 |
| **Fig S5.** Immunohistochemistry staining of α2δ-1receptor on spinal cord section of SNL rat. | 6 |
| NMR spectra of new chemical compounds | 7-32 |
| High resolution mass spectra of new chemical compounds | 33-35 |
| HPLC chromatography | 36-39 |

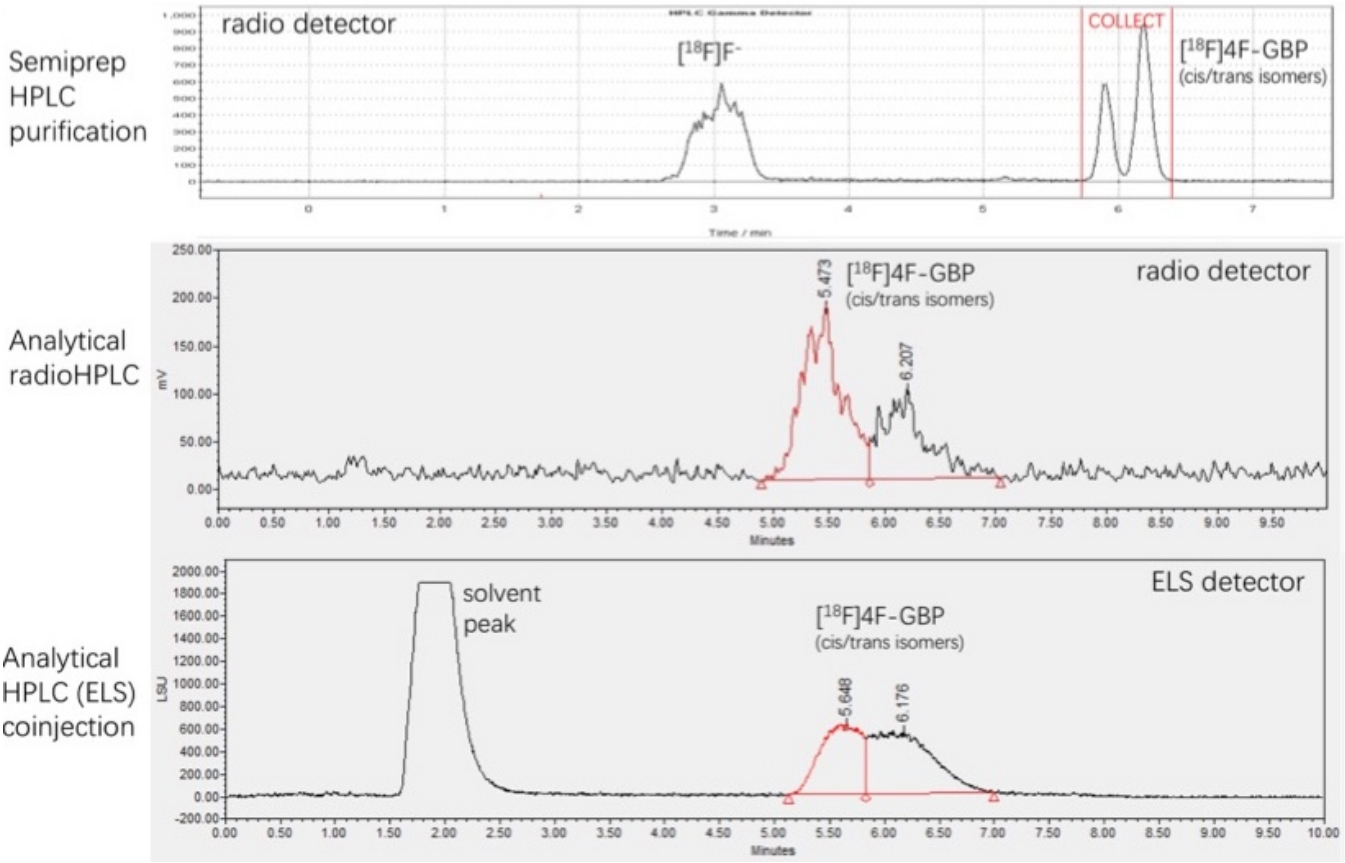


**Fig S1.** HPLC chromatography of [$^{18}$F]GBP4F coinjection (top: Semiprep HPLC; middle: analytic radioHPLC; bottom: analytic ELS-HPLC)

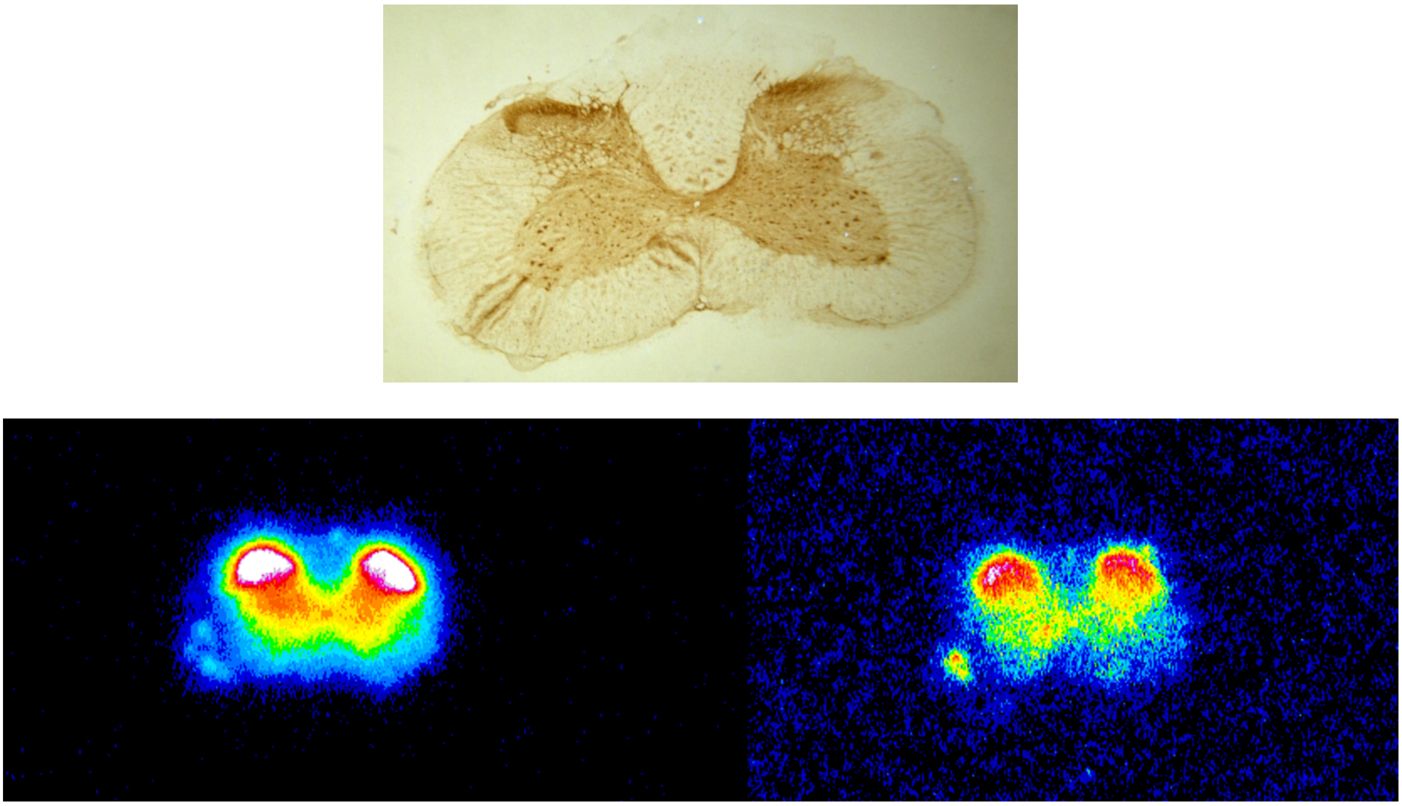

**Fig S2. Top**: Immunohistochemistry staining of α2δ-1receptor on spinal cord section of healthy rat. Approx. dimension 5 x 3.3 mm. **Bottom**: High resolution autoradiography of [$^{18}$F]tGBP4F (left) and [$^{18}$F]cGBP4F (right) in rat spinal cord slices from healthy rat.

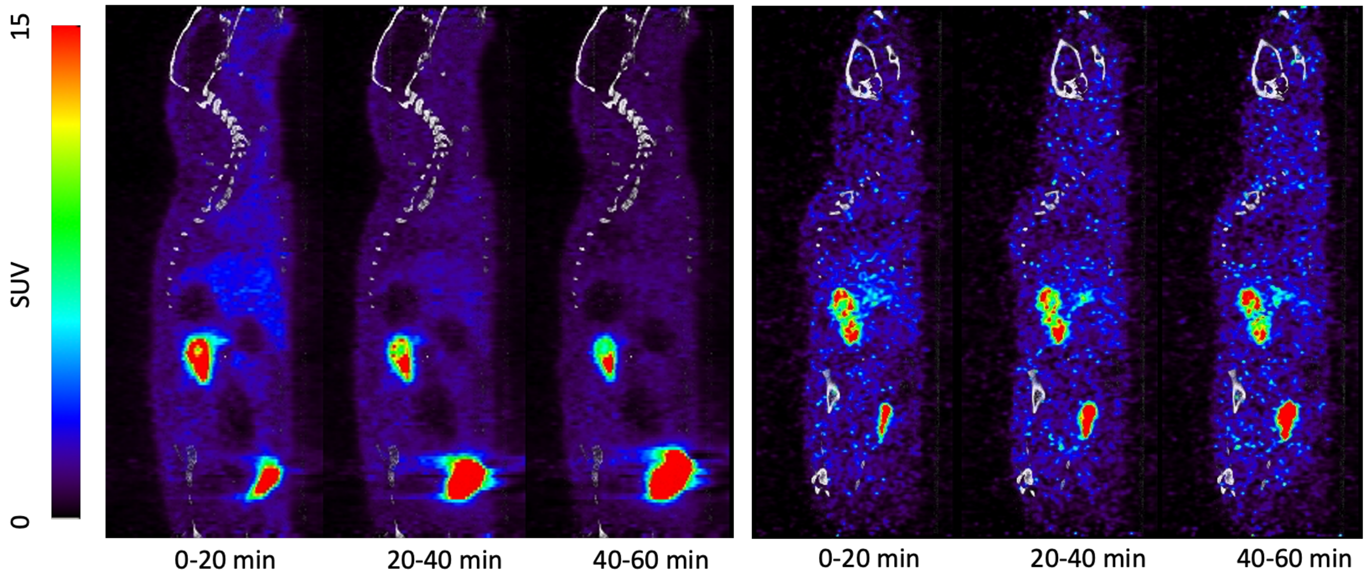


**Fig S3.** Whole body PET imaging of healthy rats (sagittal view, summed images at 0−20 min, 20−40 min, and 40−60 min intervals) with [$^{18}$F]tGBP4F (left) and [$^{18}$F]cGBP4F (right).

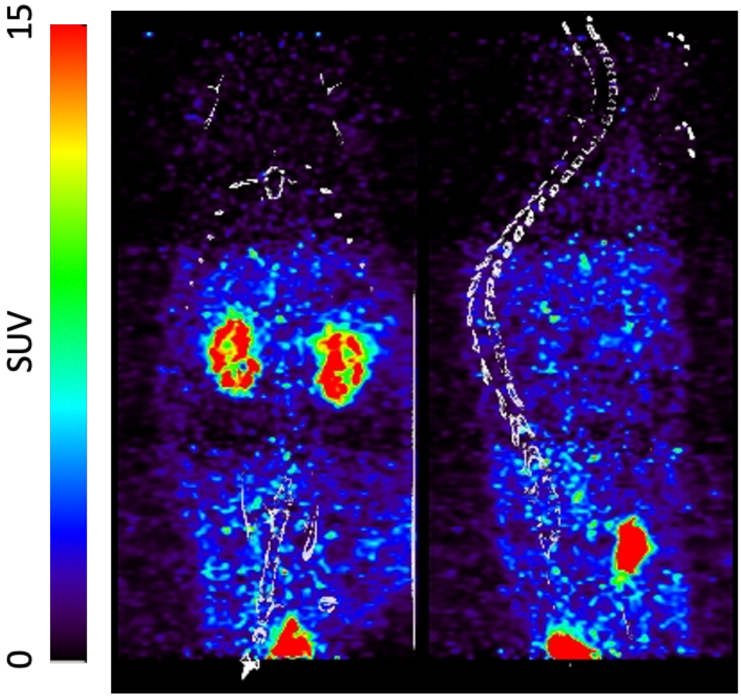


**Fig S4.** Whole body PET imaging of SNL rats (coronal view-left and sagittal view-right, summed images of 0−60 min intervals) with [$^{18}$F]tGBP4F.

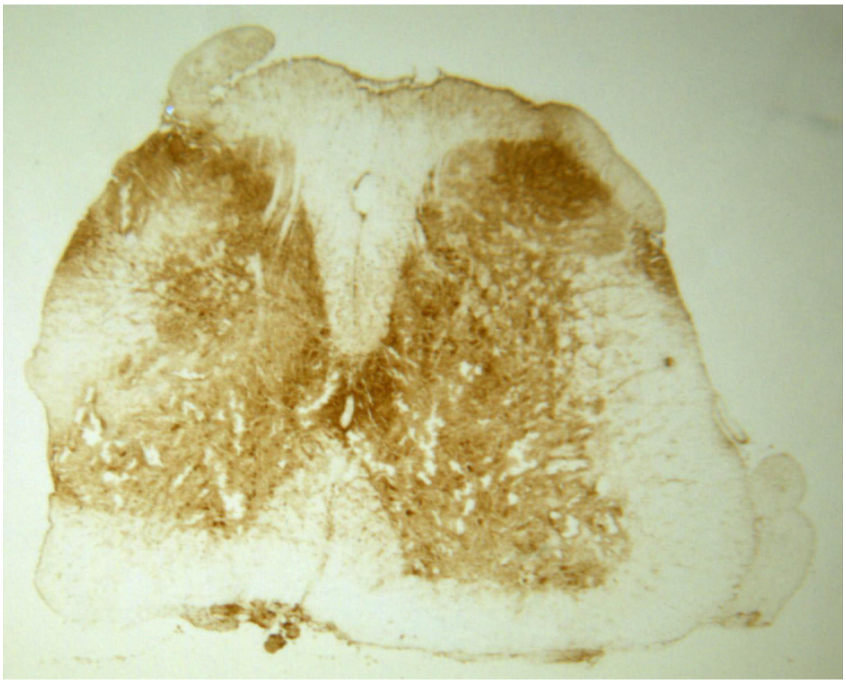

**Fig S5.** Immunohistochemistry staining of α2δ-1receptor on spinal cord section of SNL rat. Right side is the affected side. Approx. dimensions 5 x 4 mm.

### 1. NMR spectra

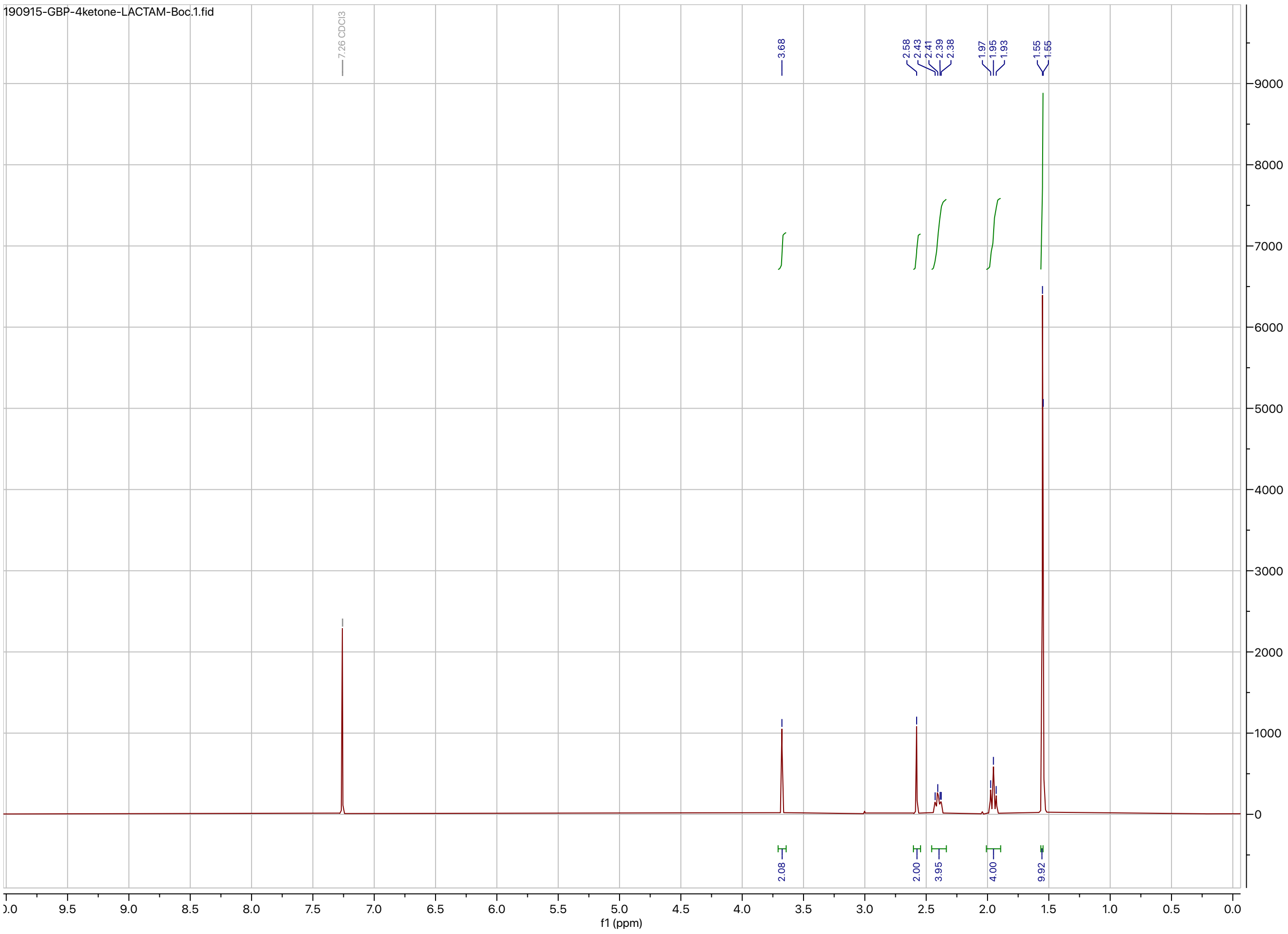


[1]H NMR spectrum of **2**.

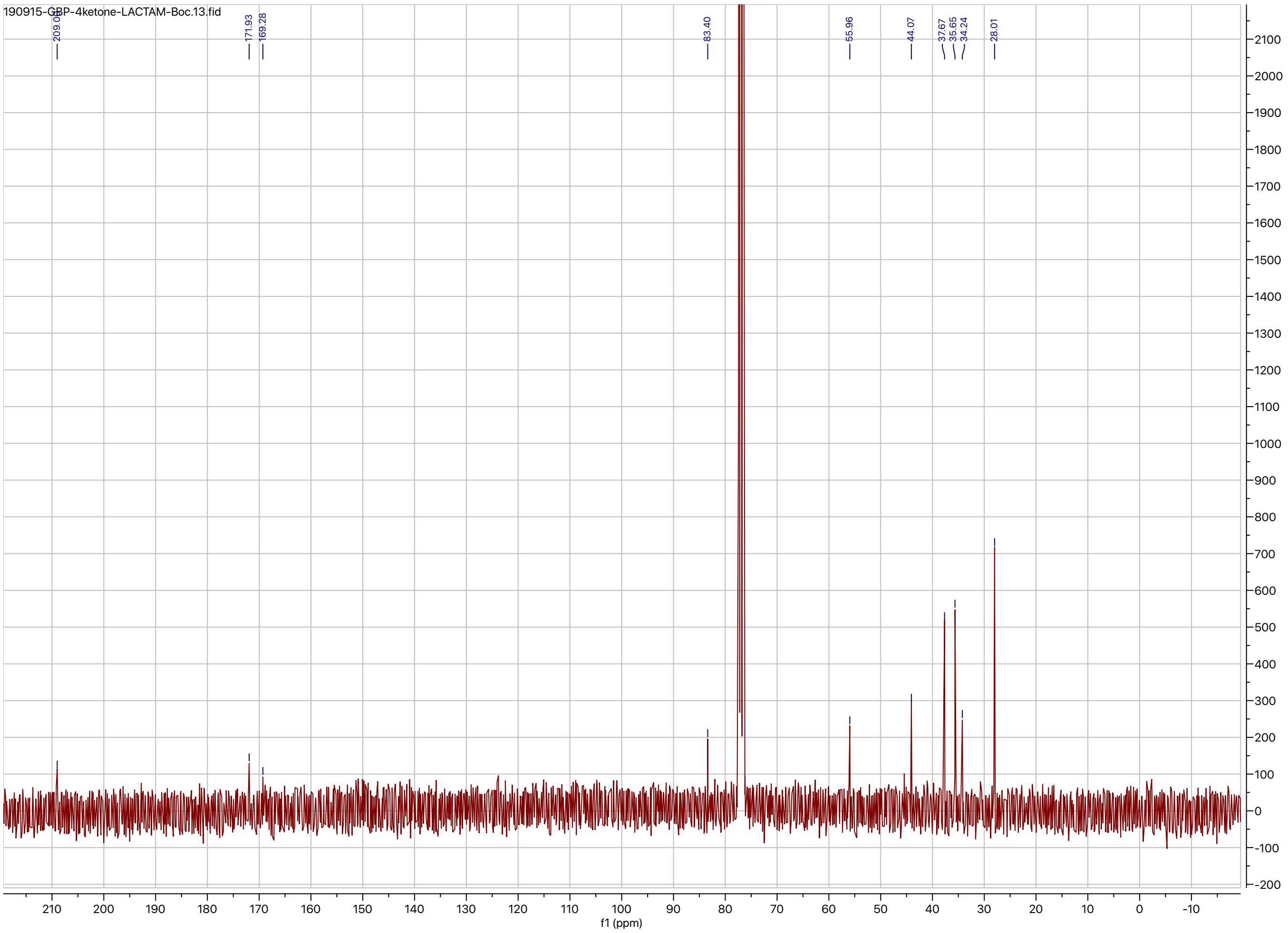


$^{13}$C NMR spectrum of **2**.

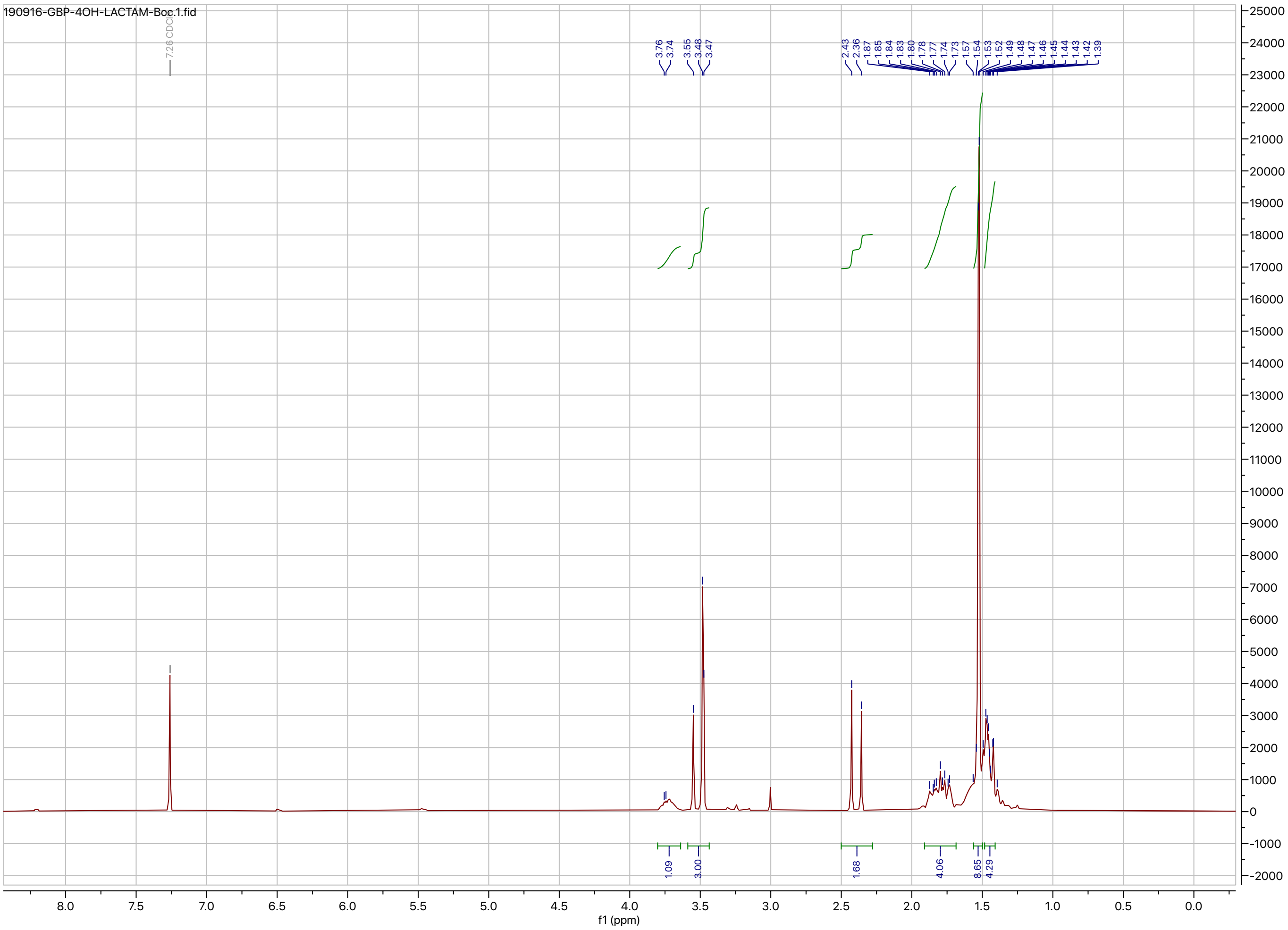


[1]H NMR spectrum of **3**.

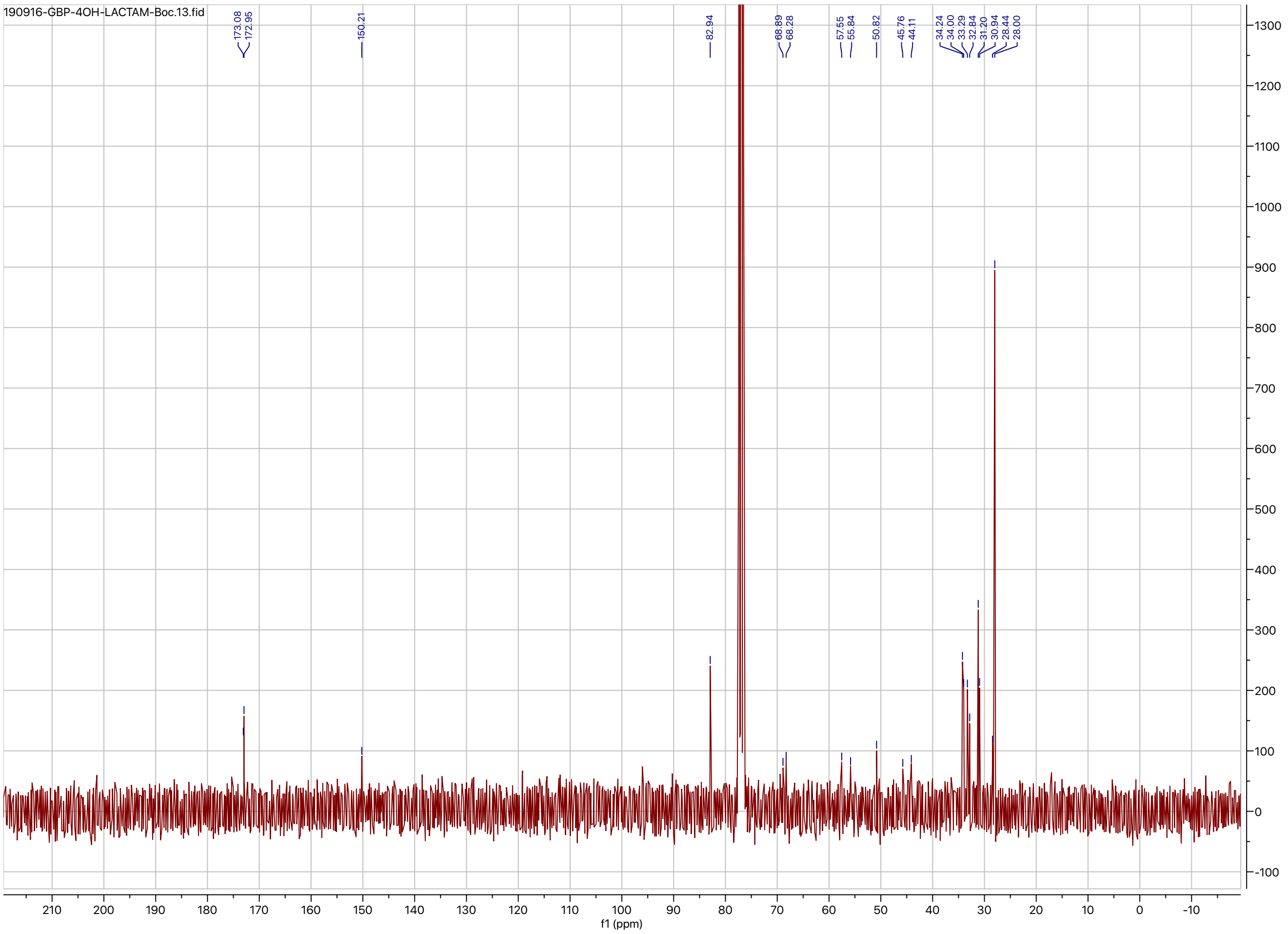


$^{13}C$ NMR spectrum of **3**.

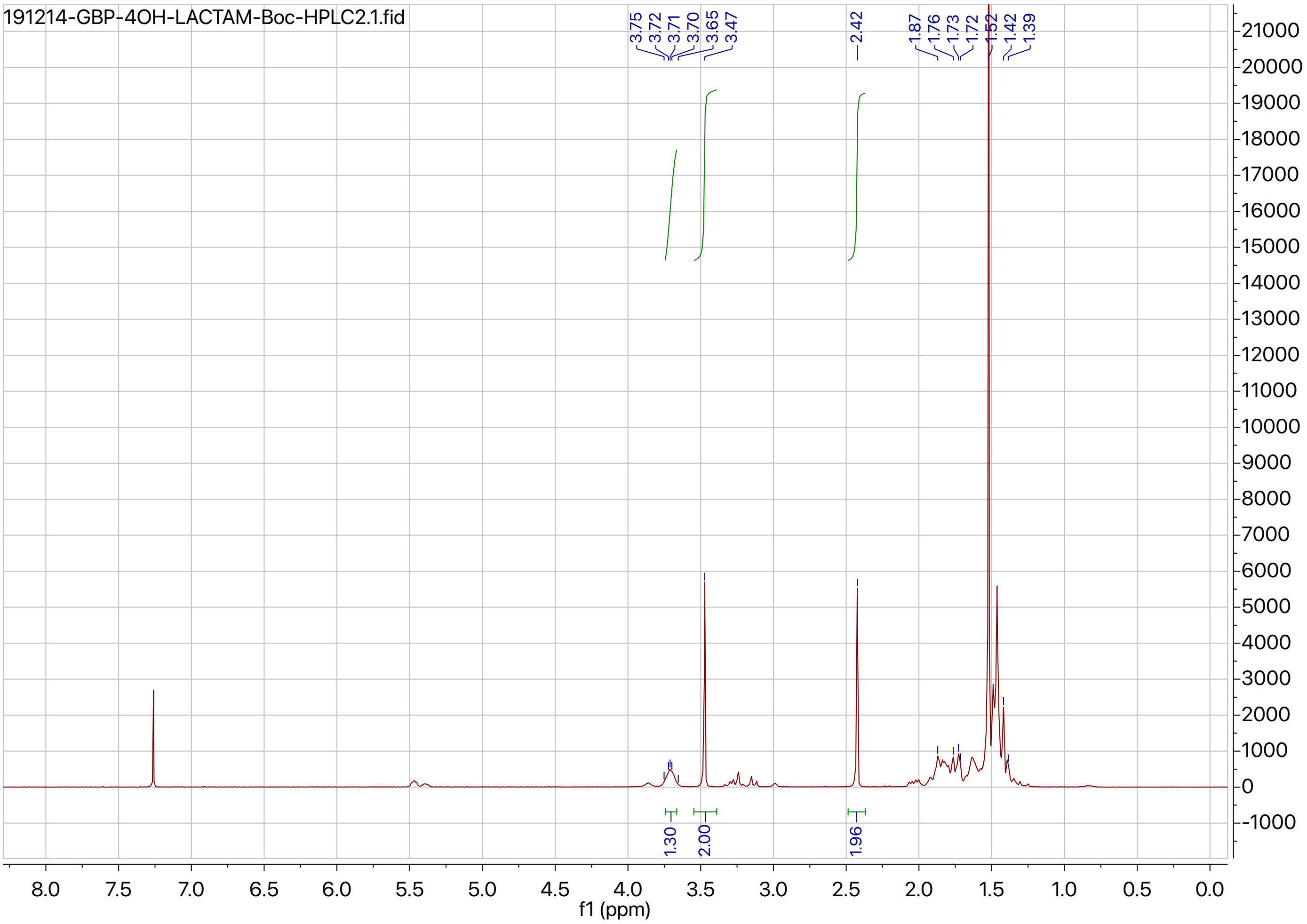


[1]H NMR spectrum of **3a**.

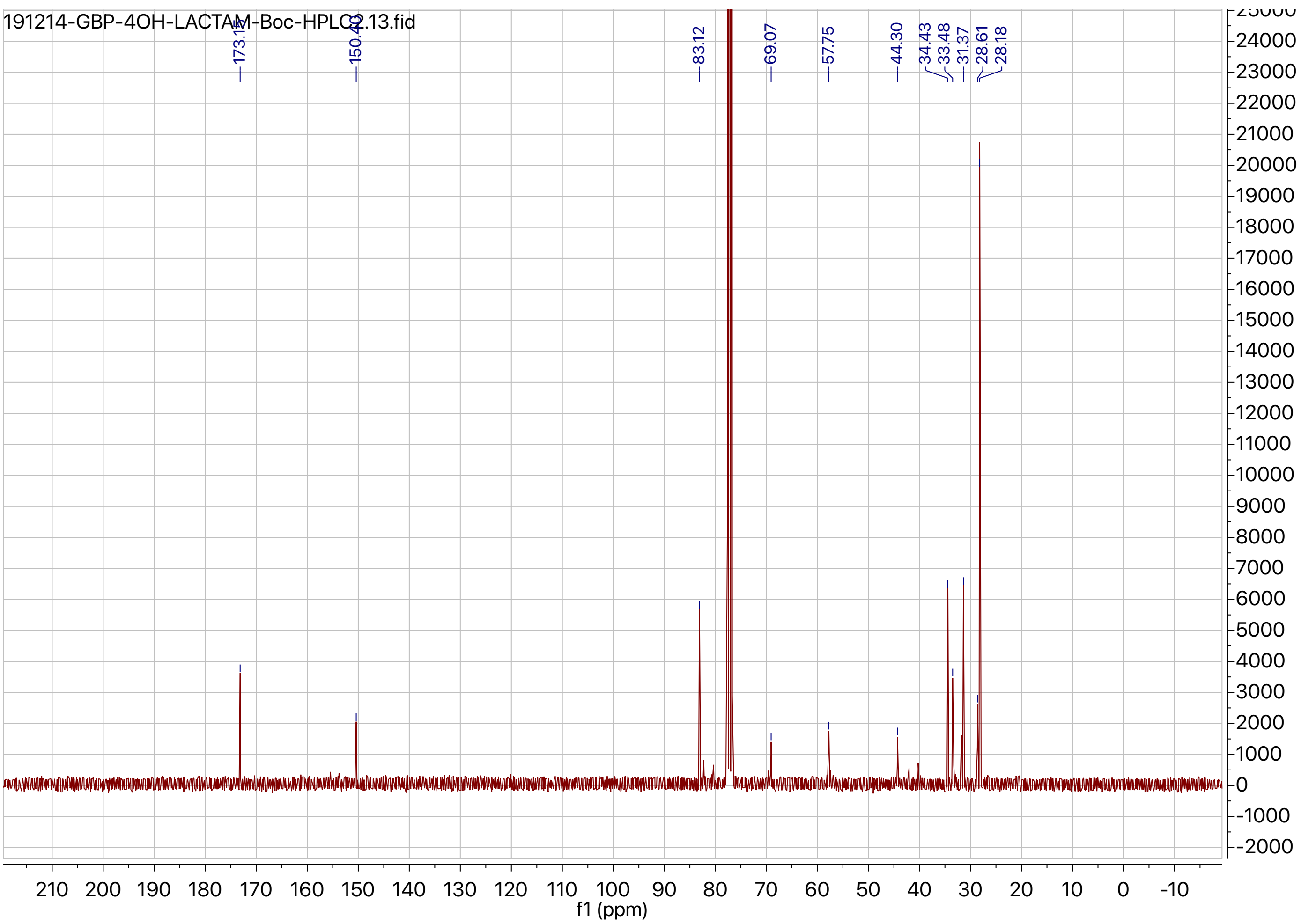


$^{13}$C NMR spectrum of **3a**.

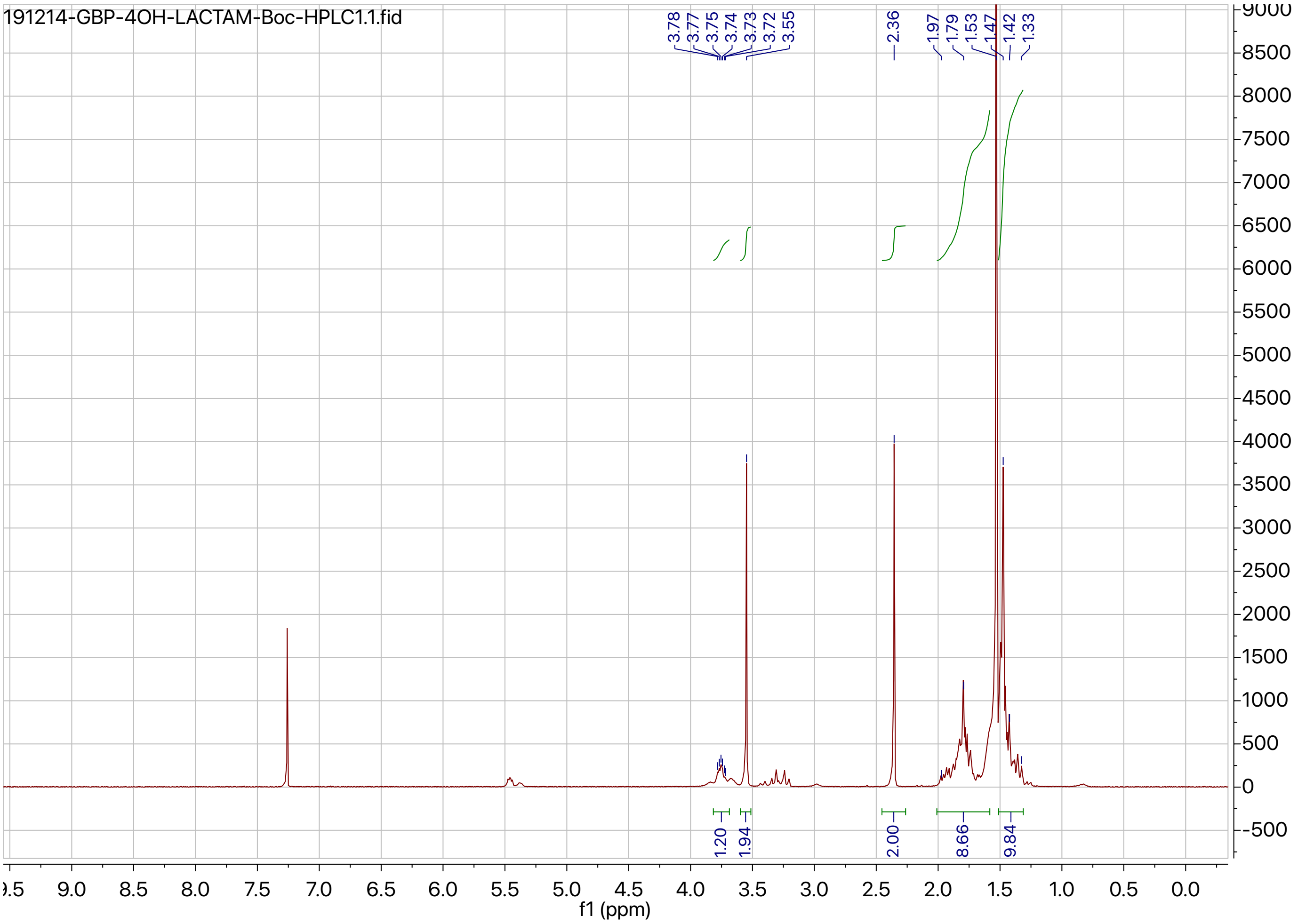


[1]H NMR spectrum of **3b**.

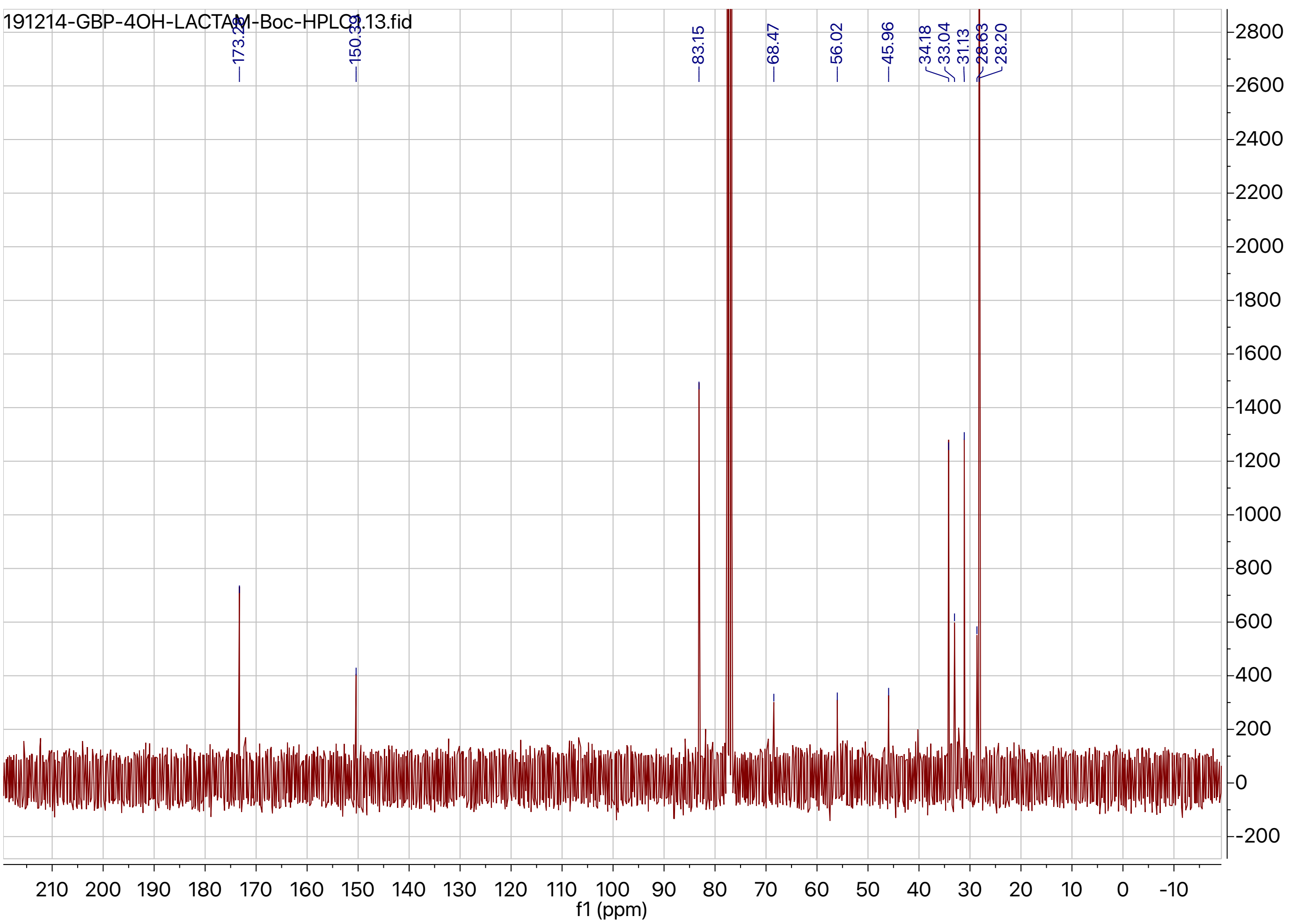


$^{13}$C NMR spectrum of **3b**.

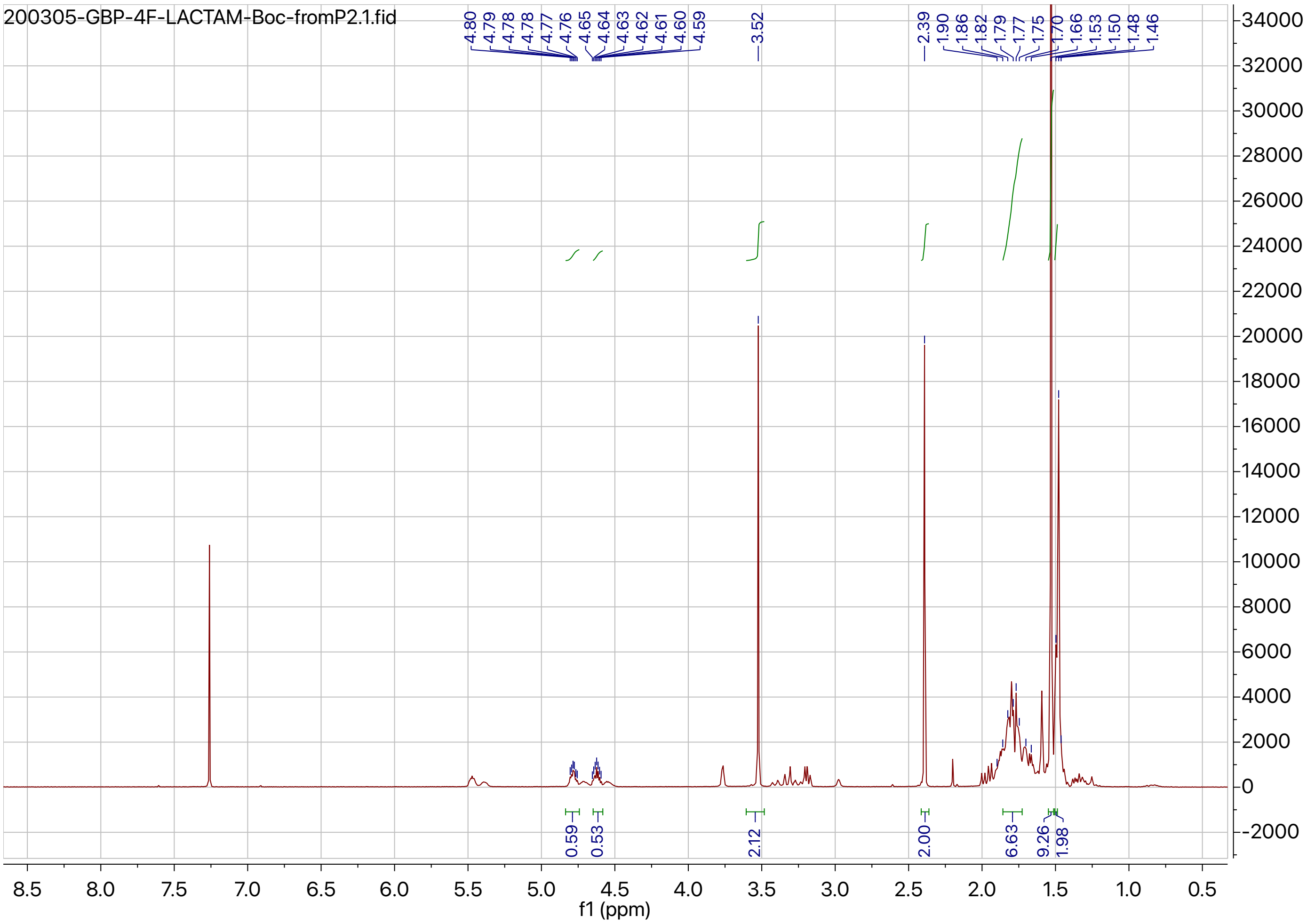


[1]H NMR spectrum of **4a**.

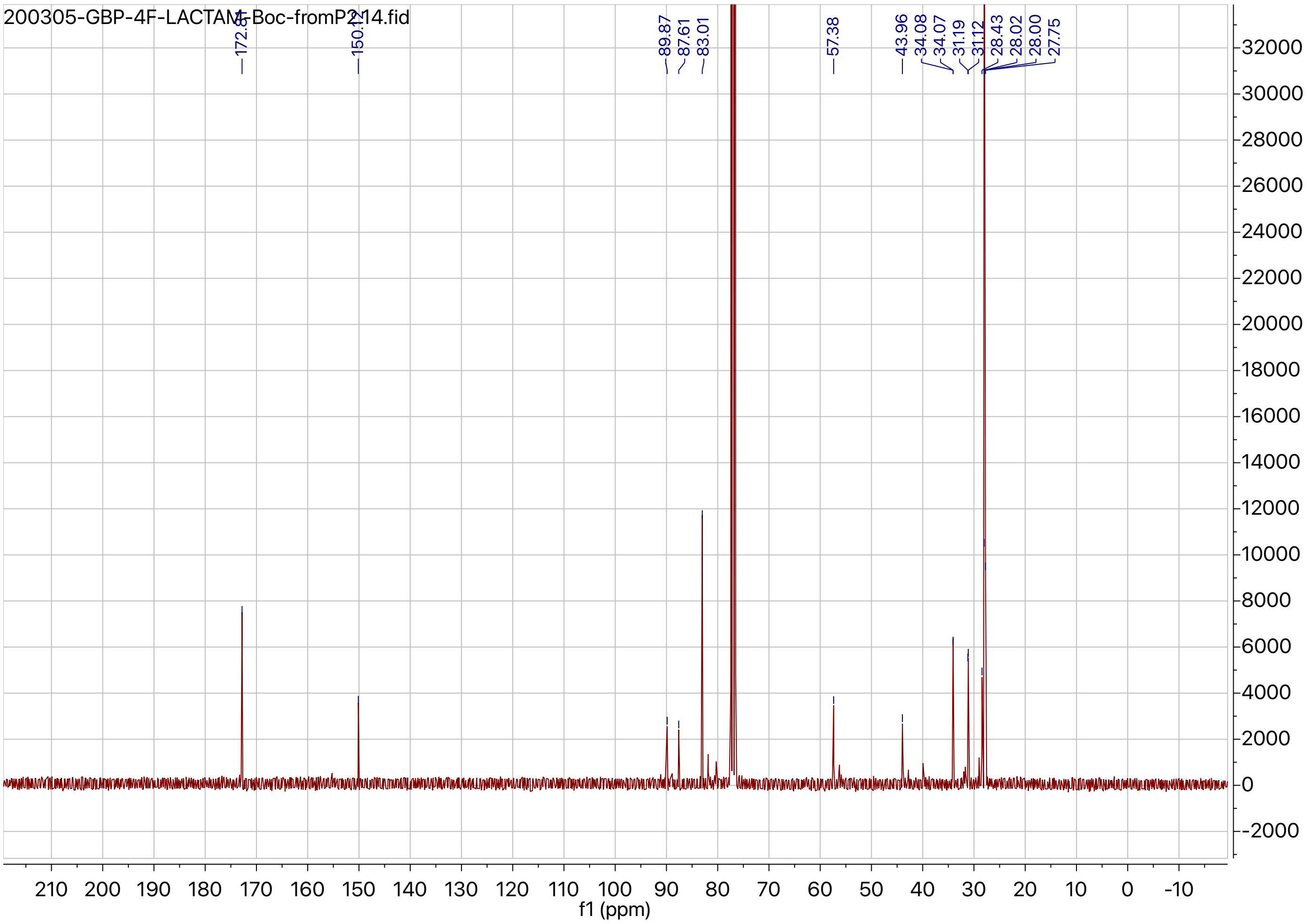


$^{13}$C NMR spectrum of **4a**.

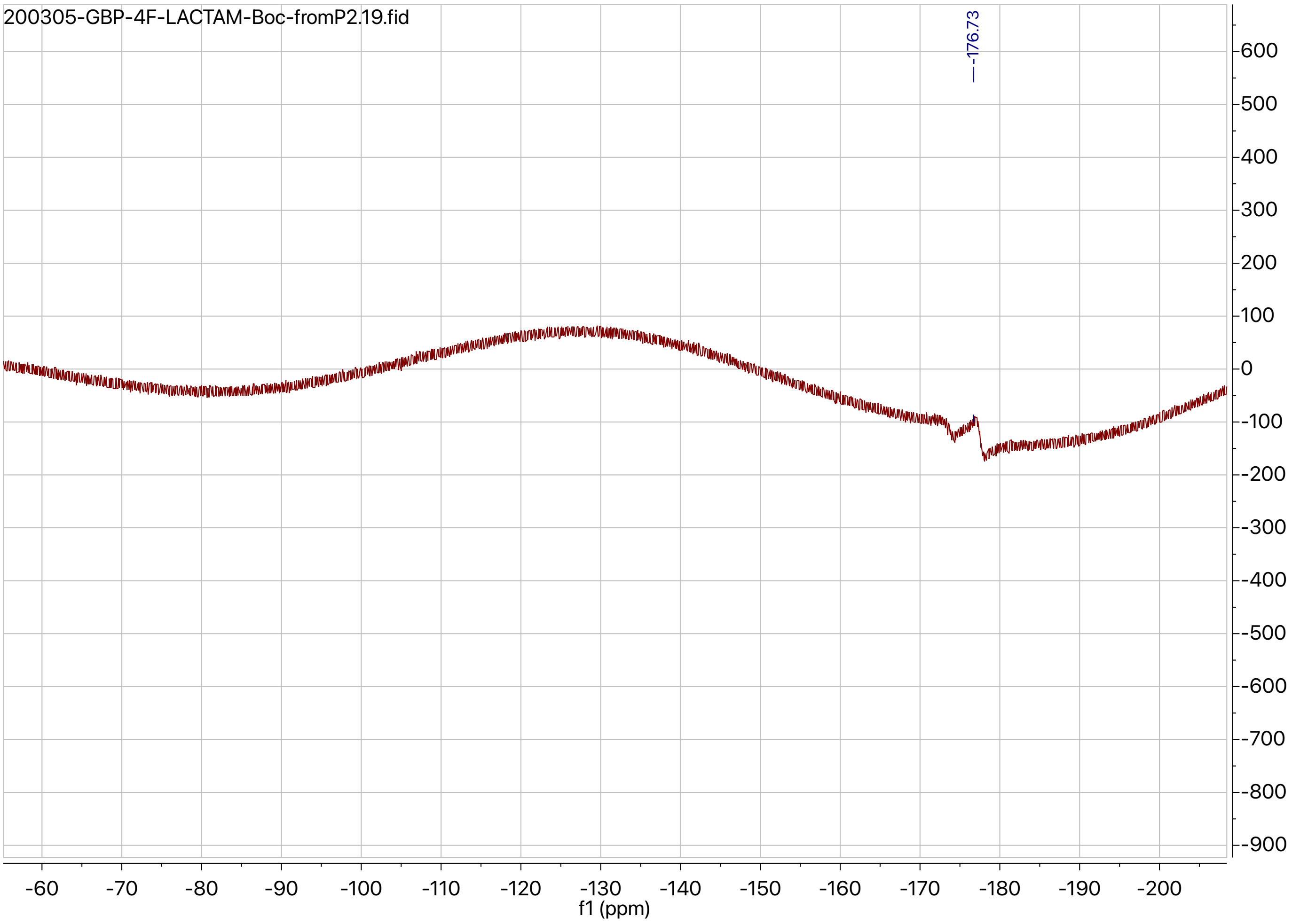


$^{19}$F NMR spectrum of **4a**.

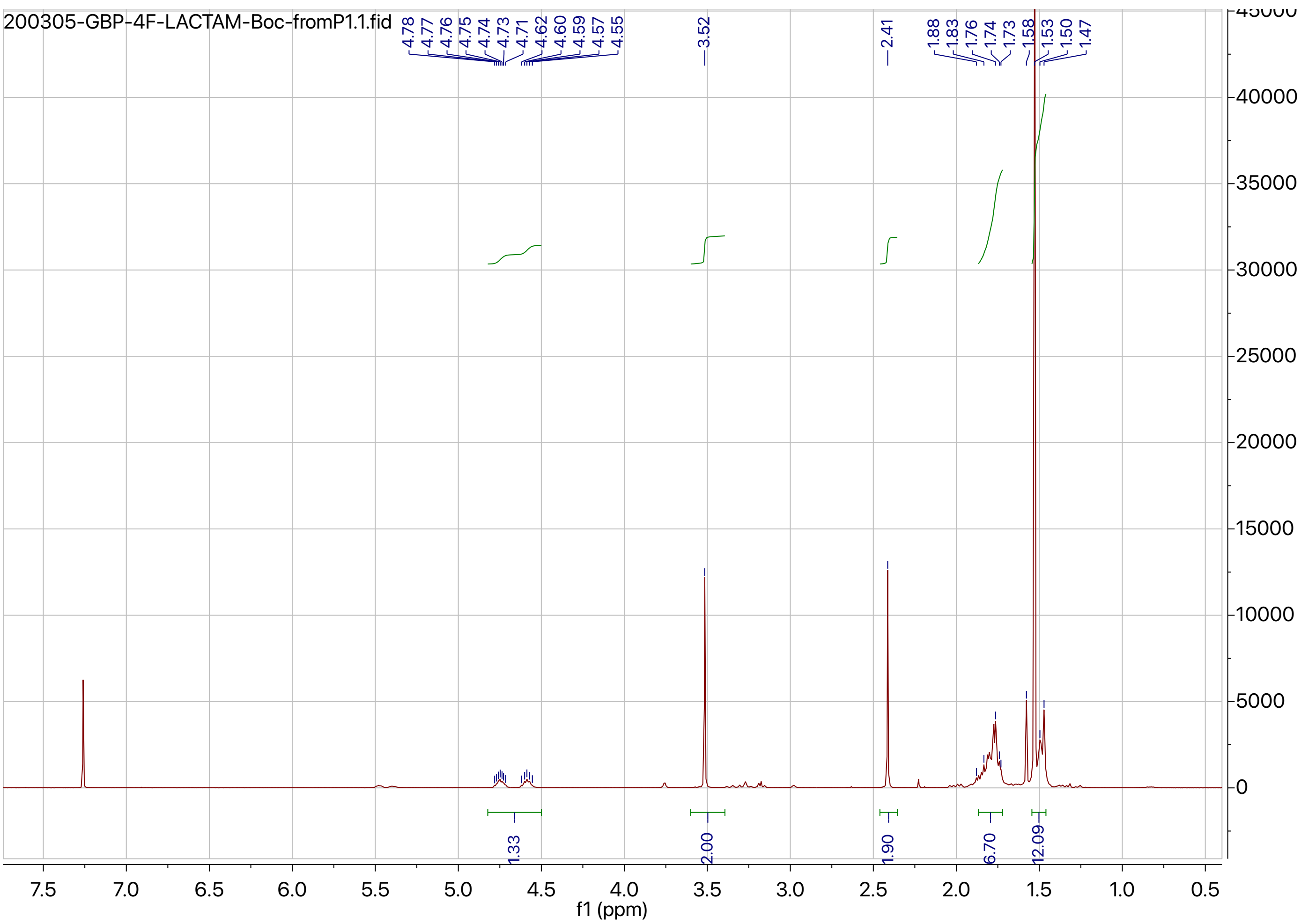


$^1$H NMR spectrum of **4b**.

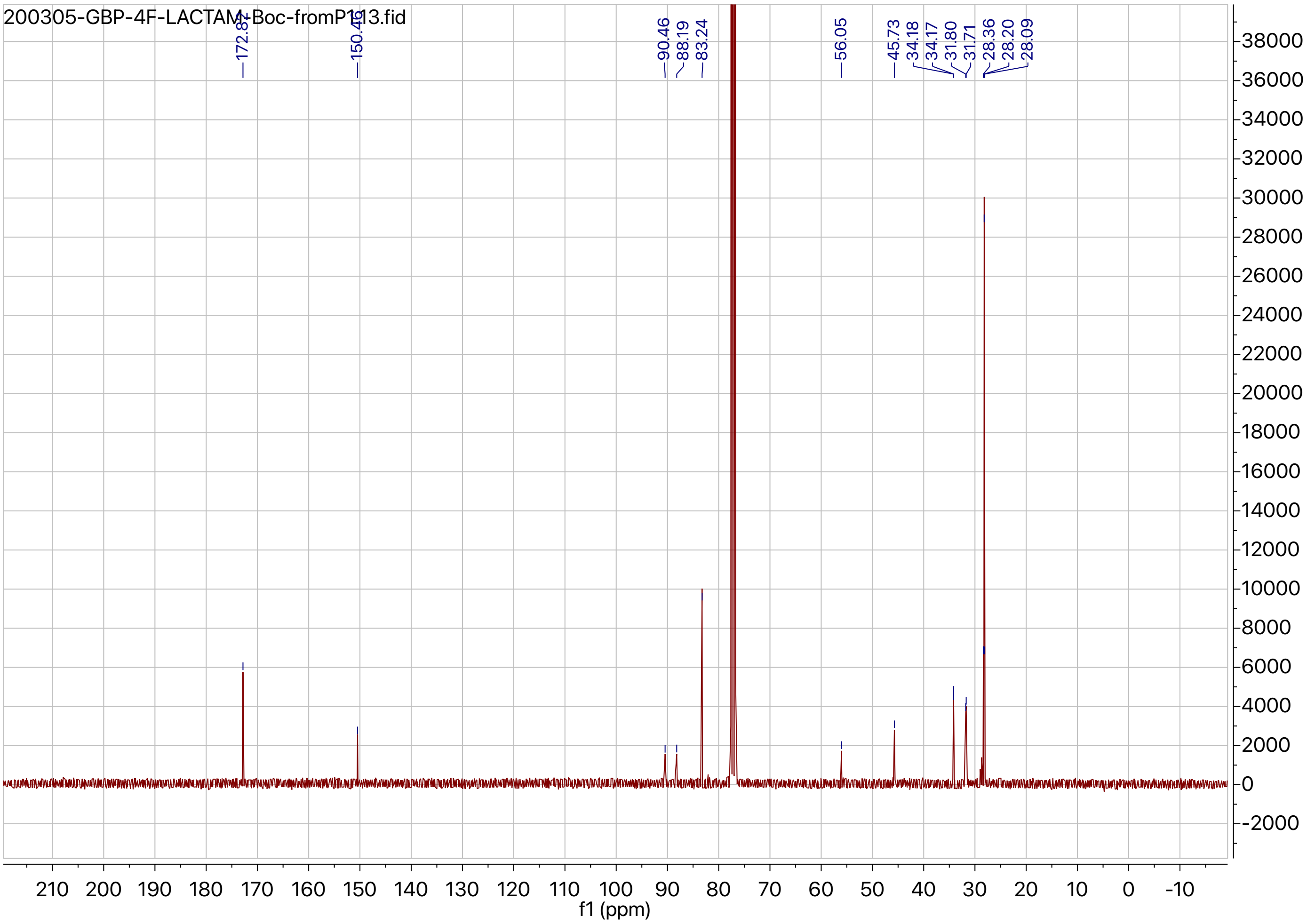


[13]C NMR spectrum of **4b**.

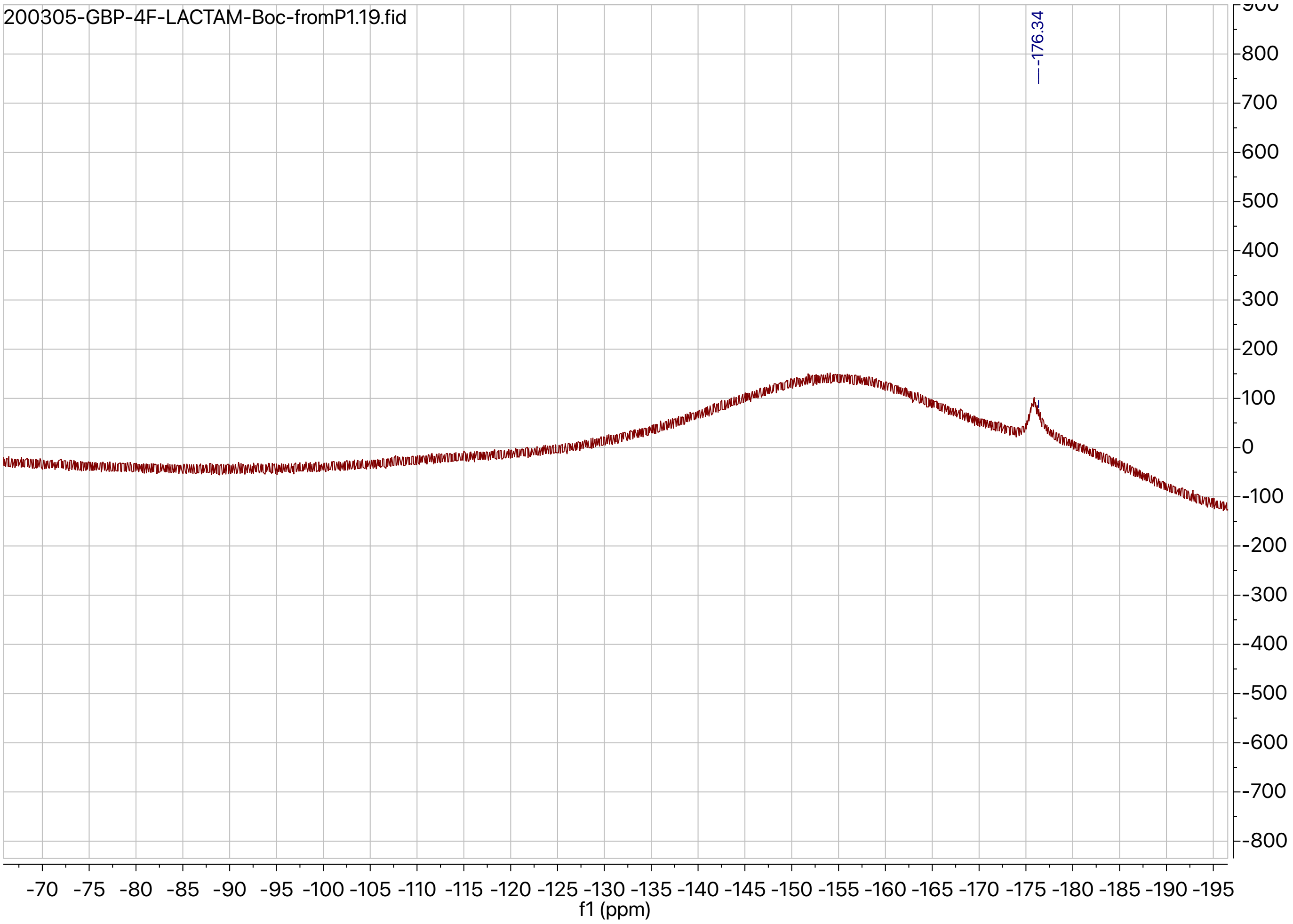


$^{19}$F NMR spectrum of **4b**.

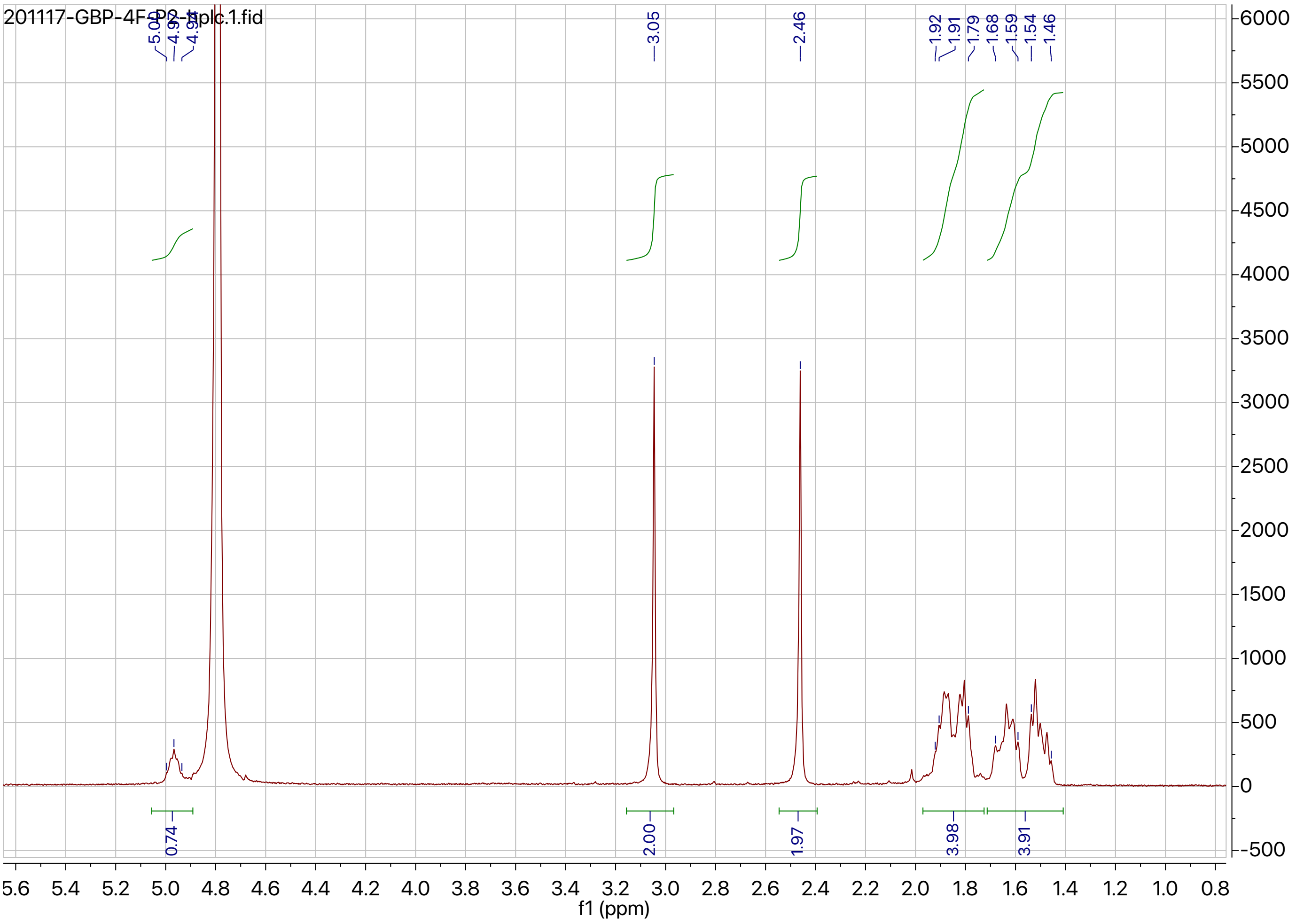


[1]H NMR spectrum of **5a**.

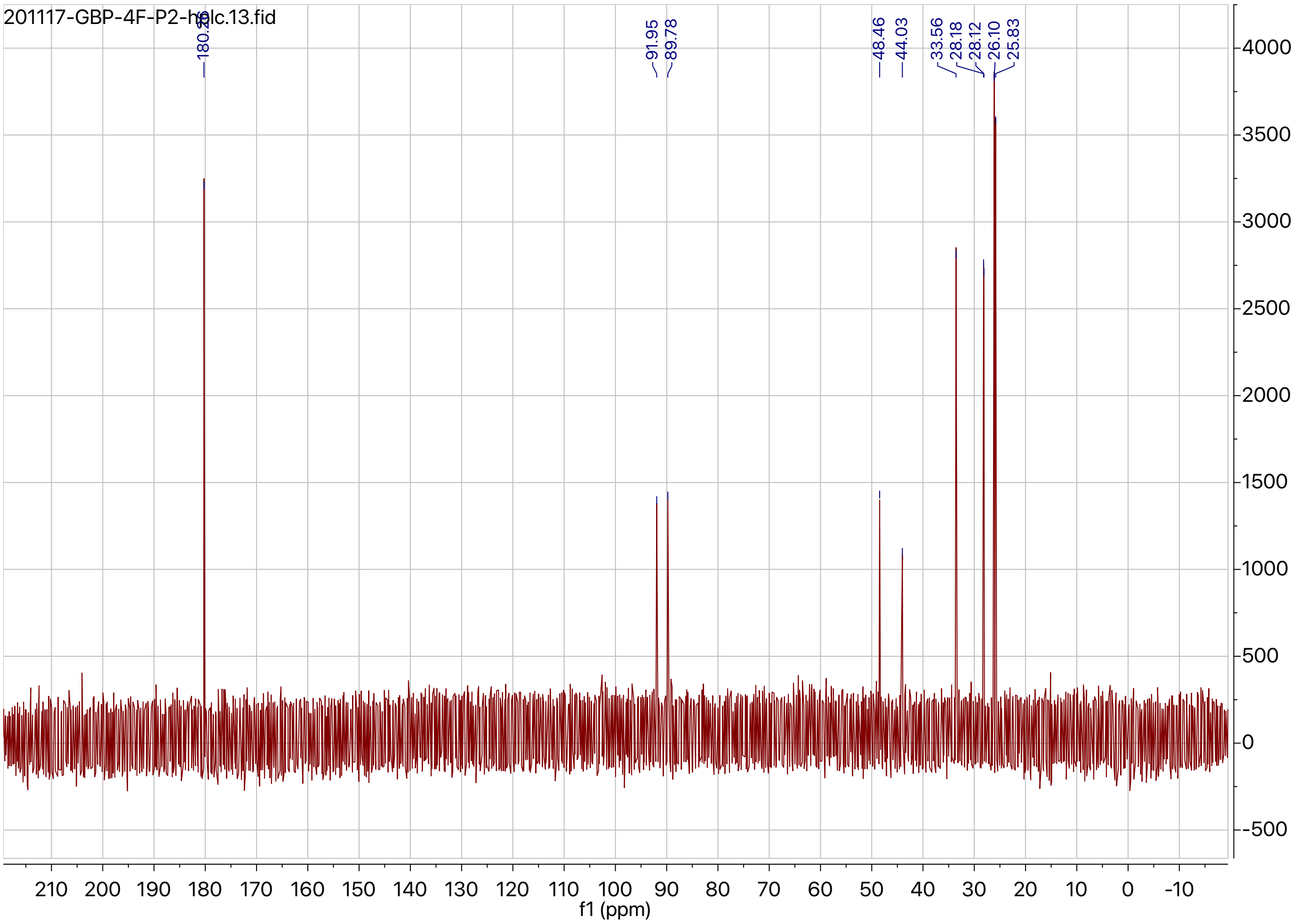


$^{13}$C NMR spectrum of **5a**.

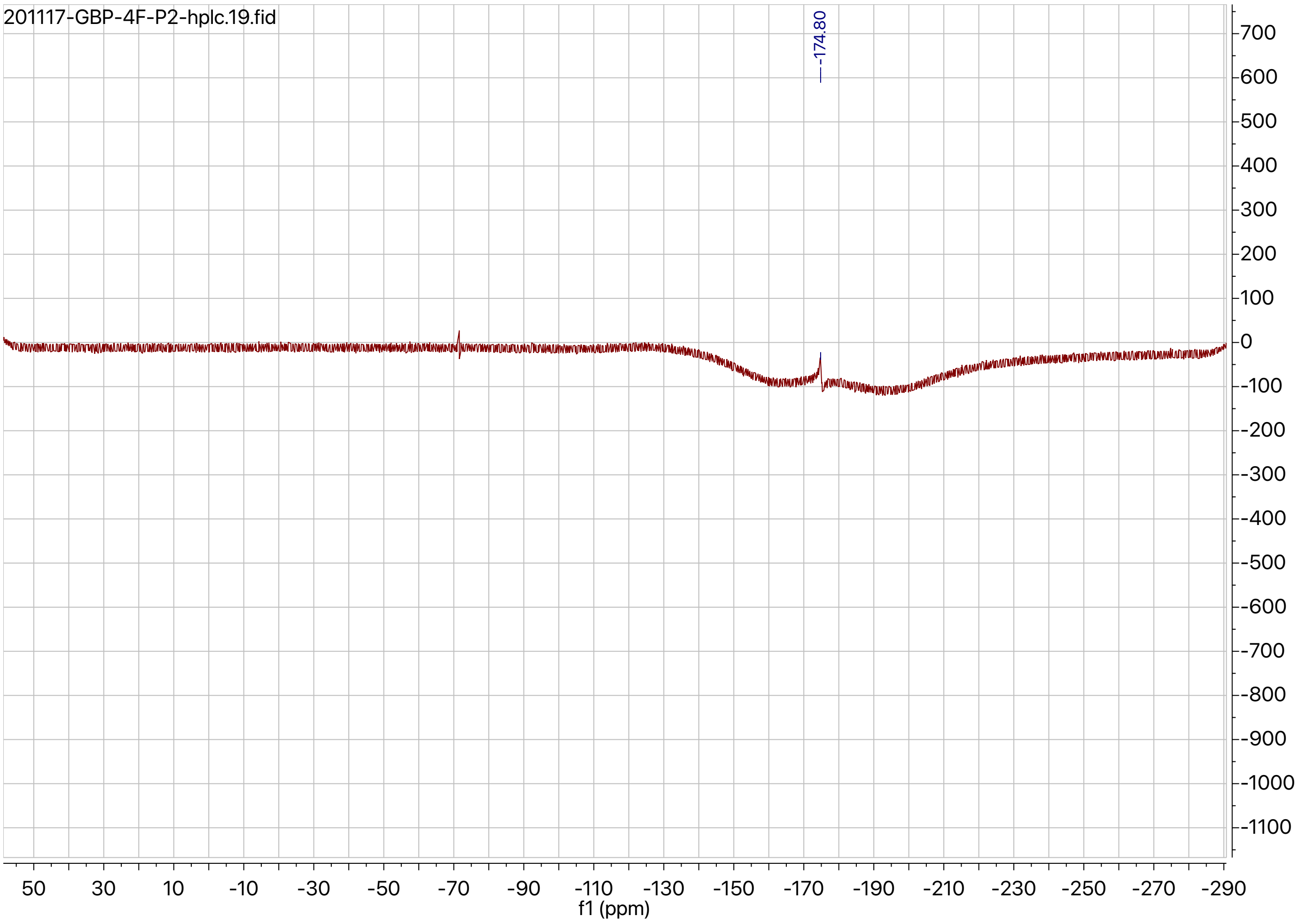


$^{19}F$ NMR spectrum of **5a**.

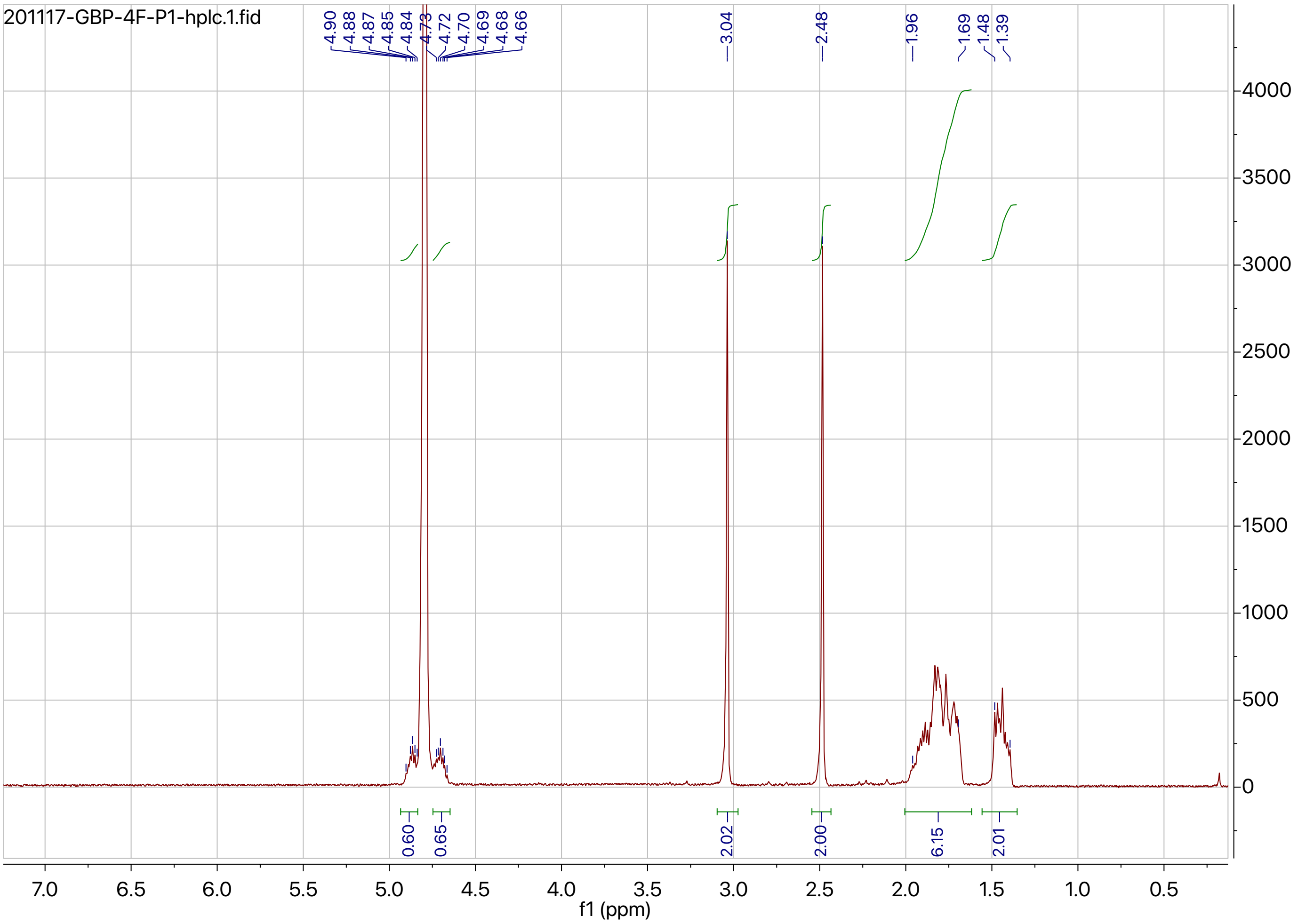


$^{1}$H NMR spectrum of **5b**.

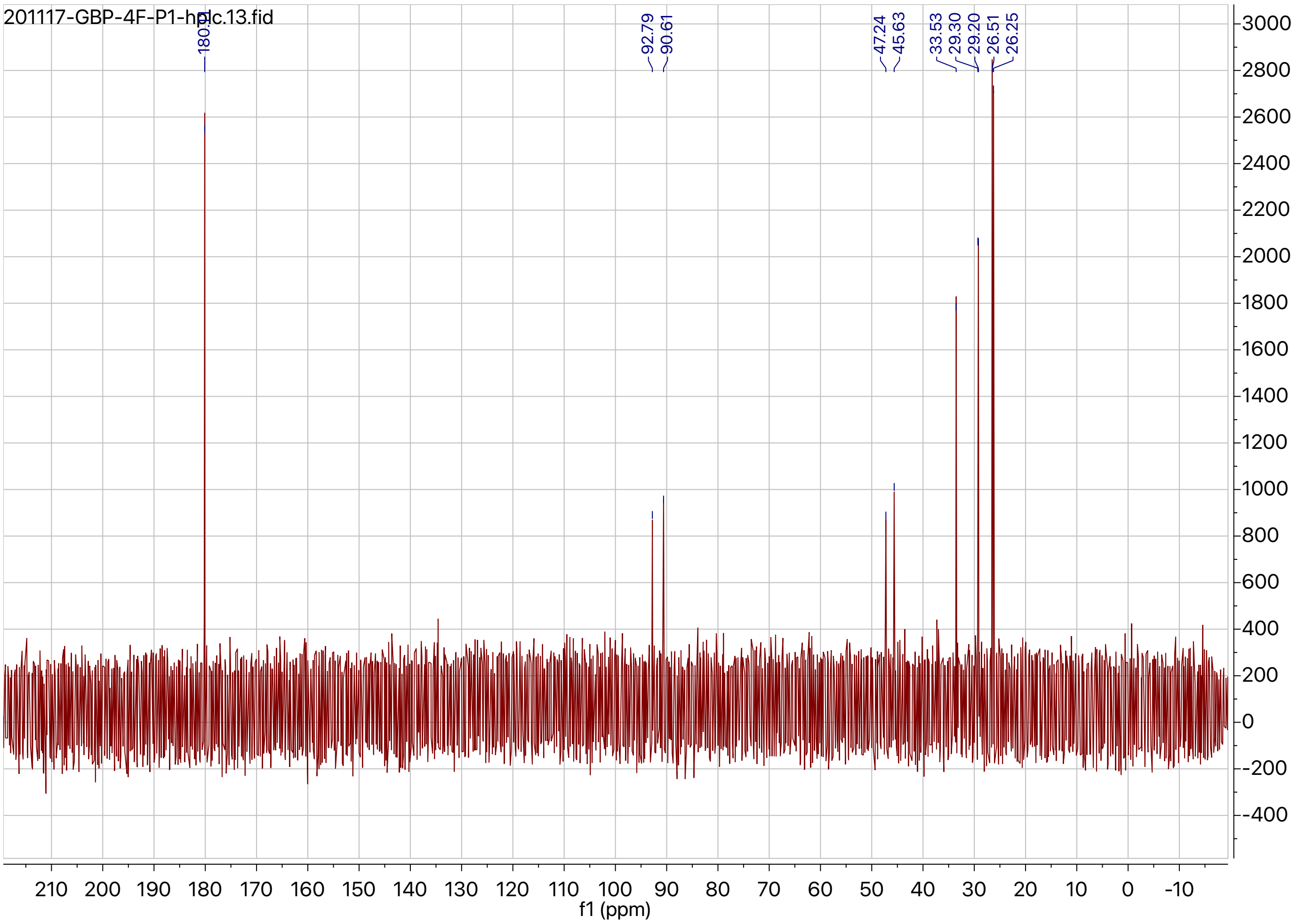


$^{13}$C NMR spectrum of **5b**.

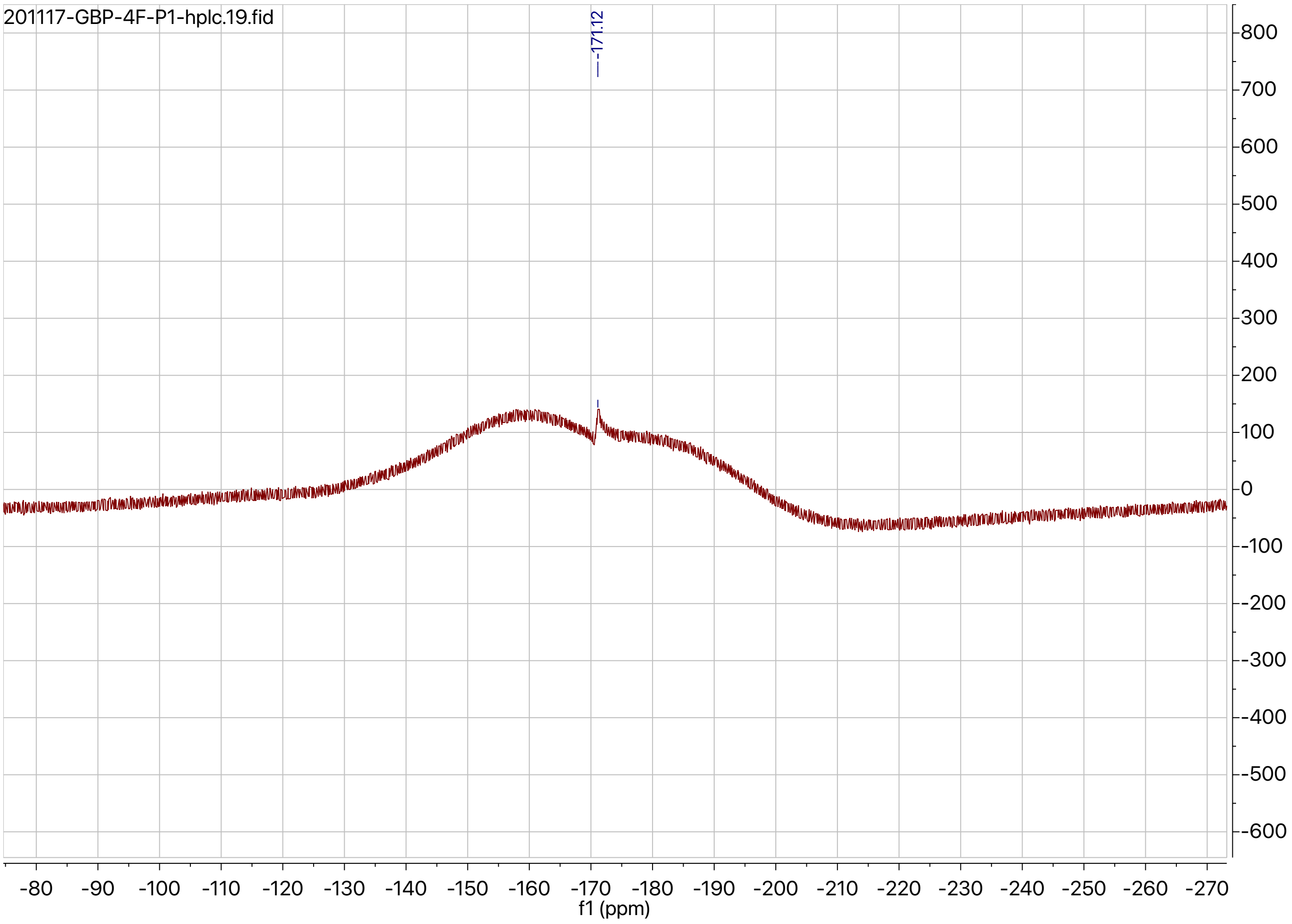


[19]F NMR spectrum of **5b**.

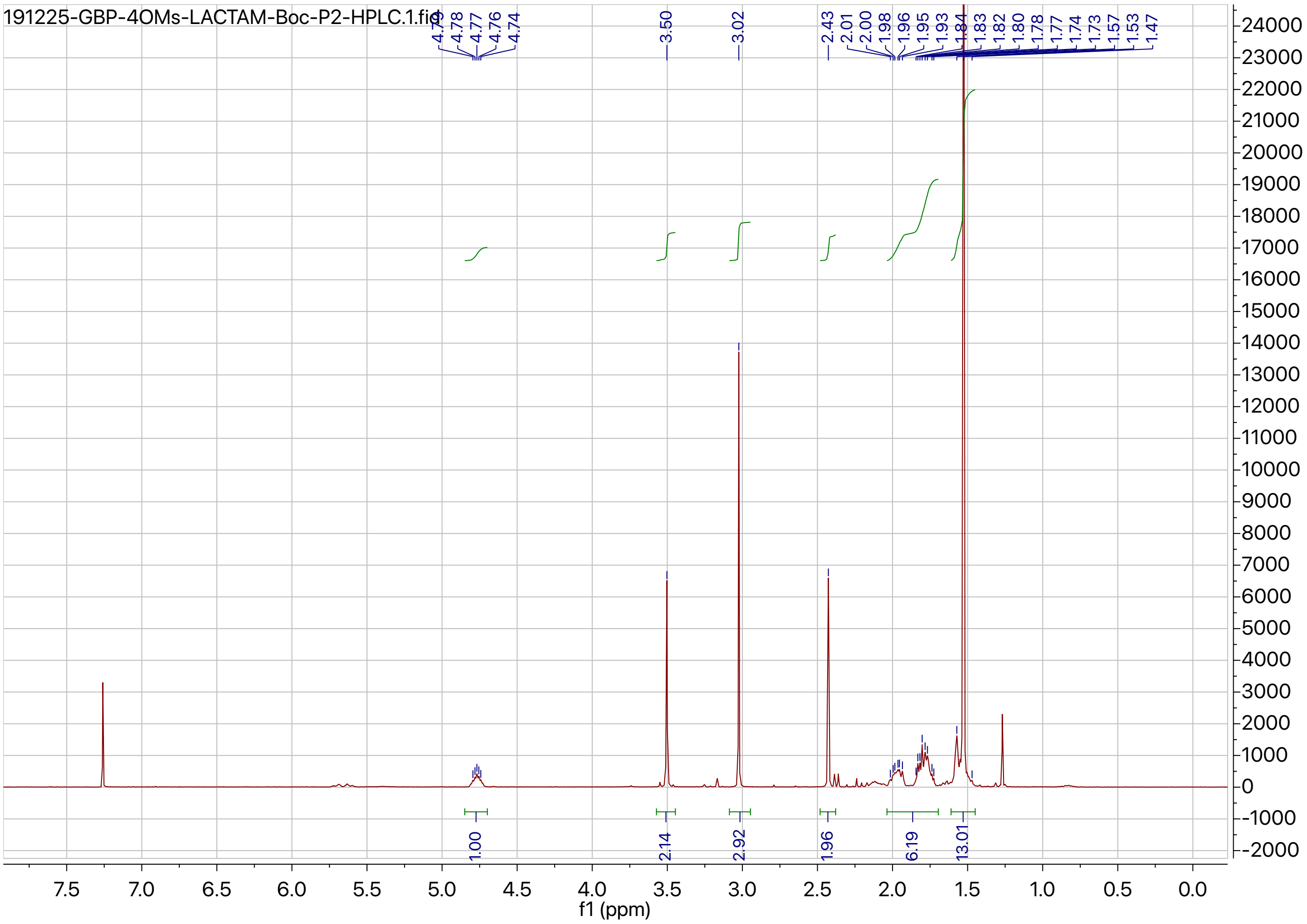


[1]H NMR spectrum of **6a**.

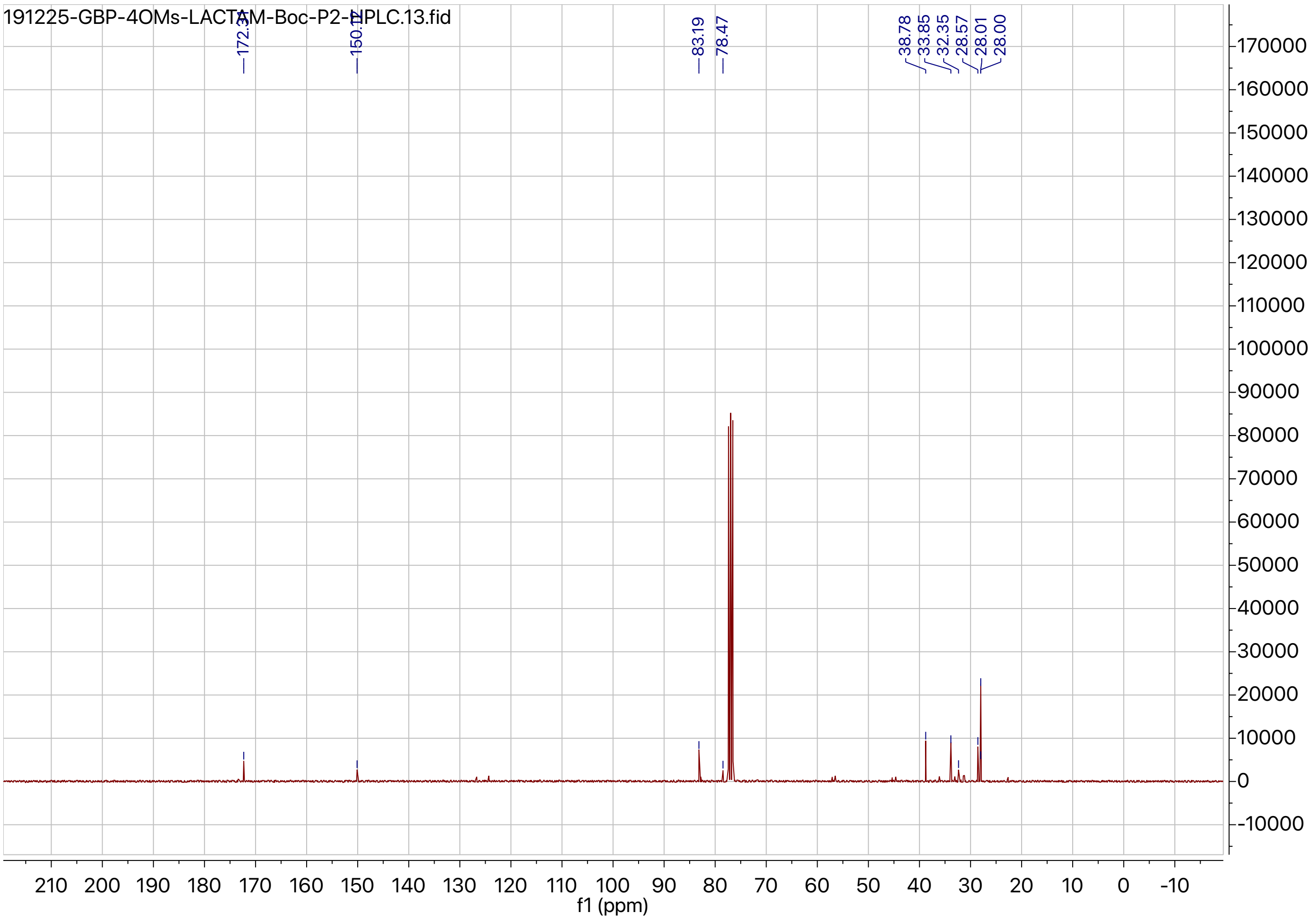


$^{13}$C NMR spectrum of **6a**.

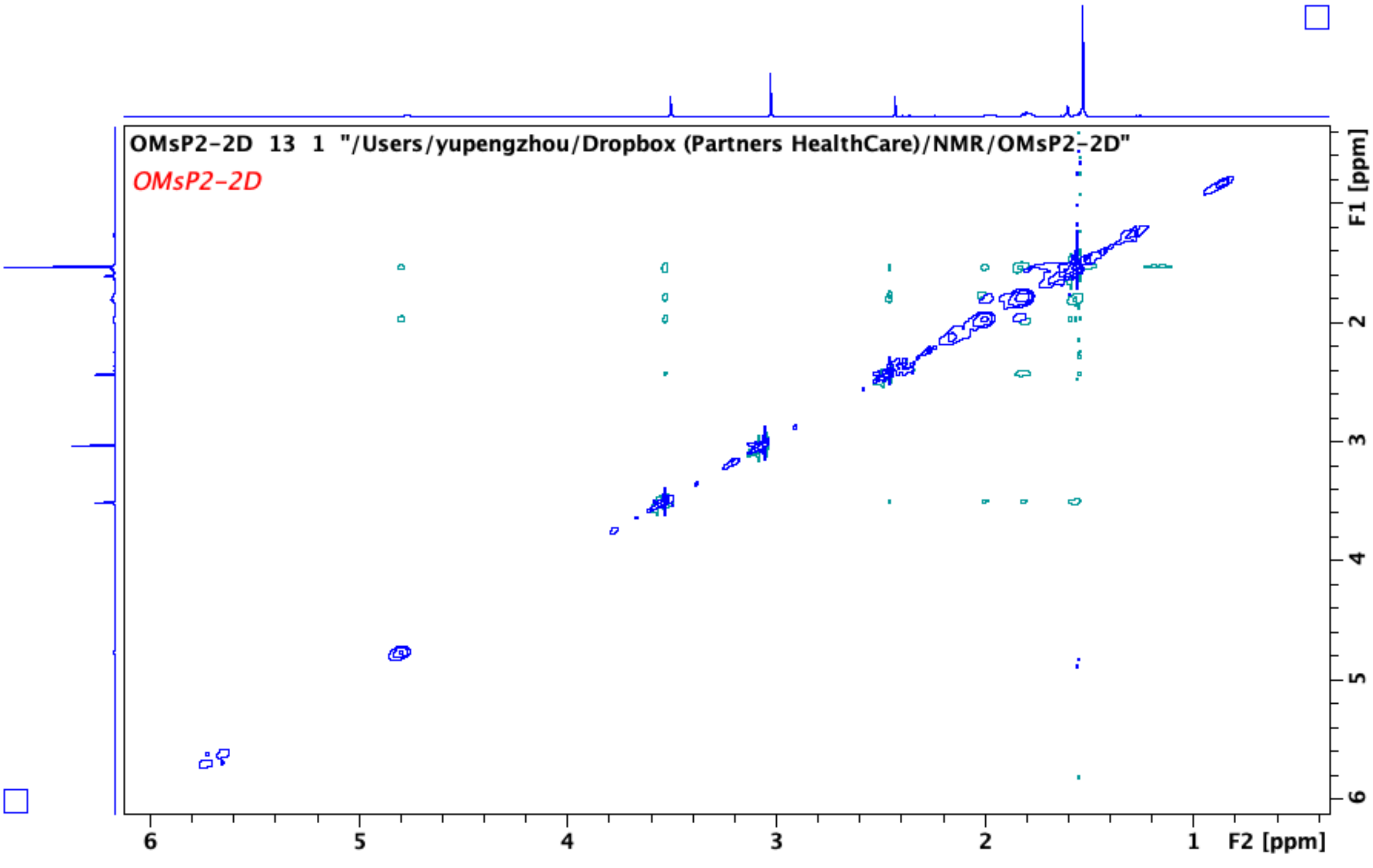


$^1$H-$^1$H NOESY NMR of **6a**.

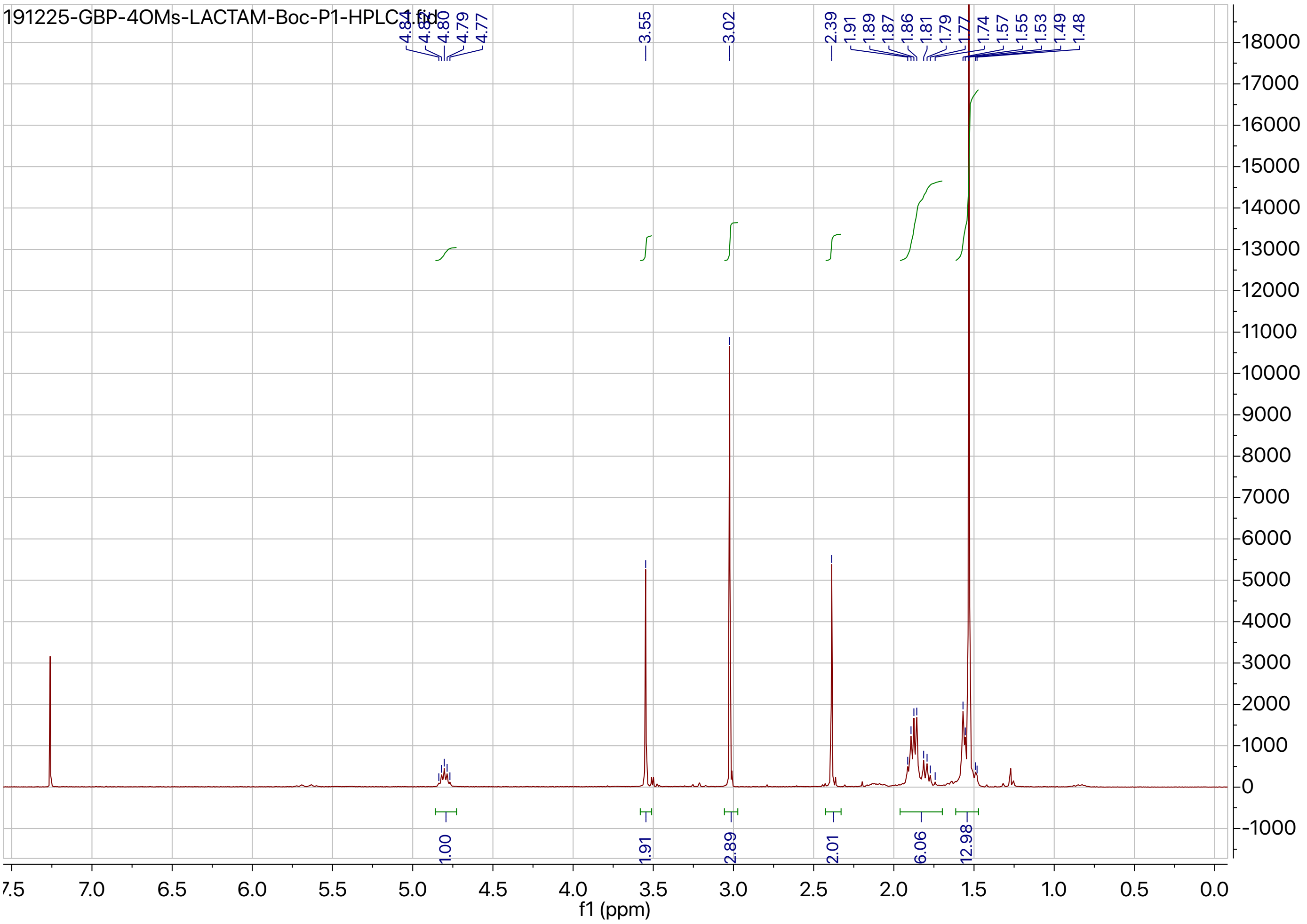


[1]H NMR spectrum of **6b**.

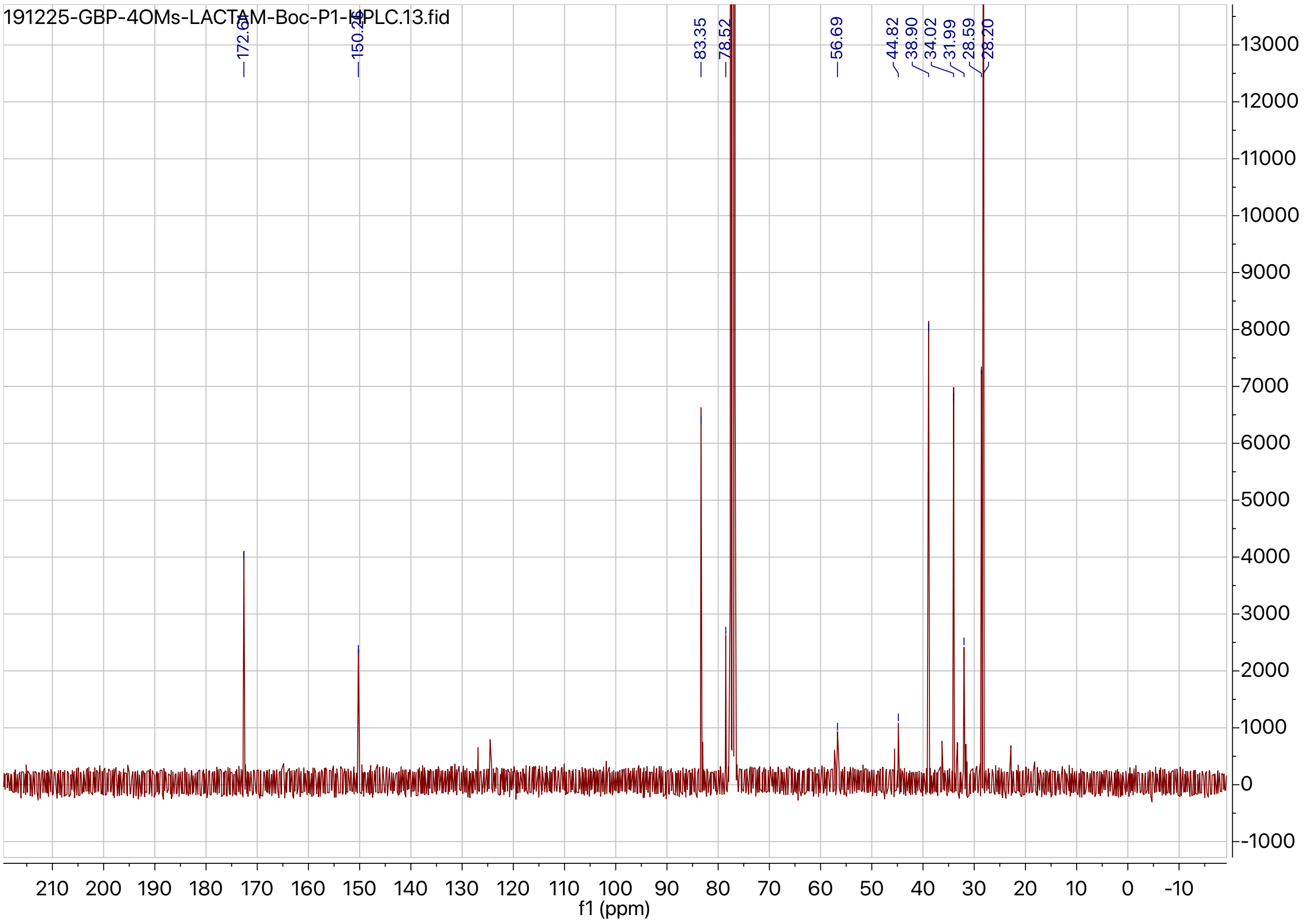


$^{13}$C NMR spectrum of **6b**.

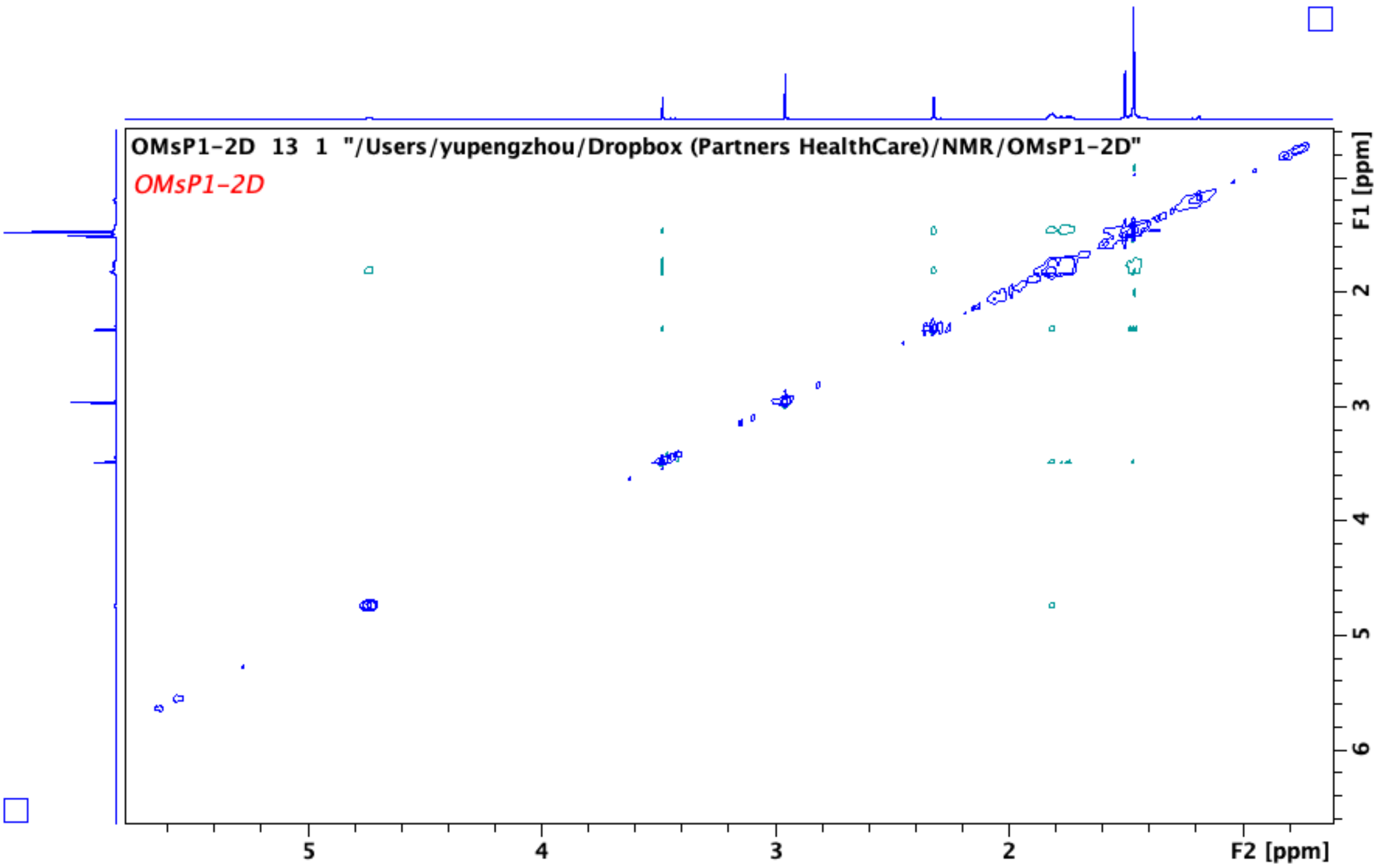


$^{1}H$-$^{1}H$ NOESY NMR of **6b**.

#### 2. High resolution mass spectra

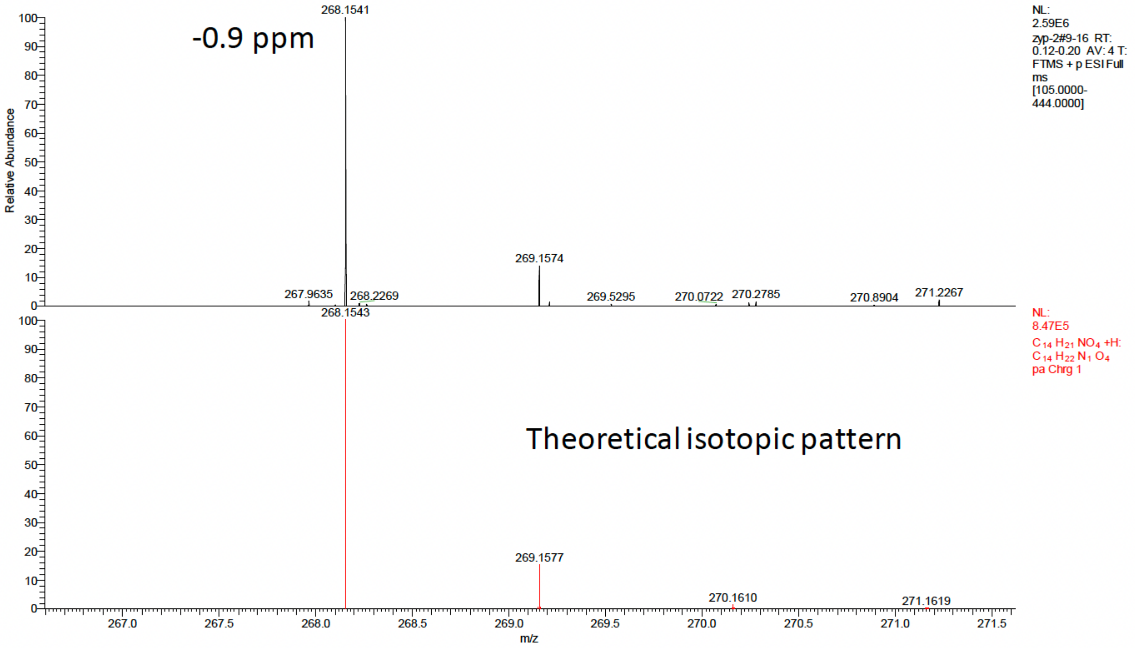


High resolution mass spectrum of **2** (top: experimental data; bottom: theoretical simulation)

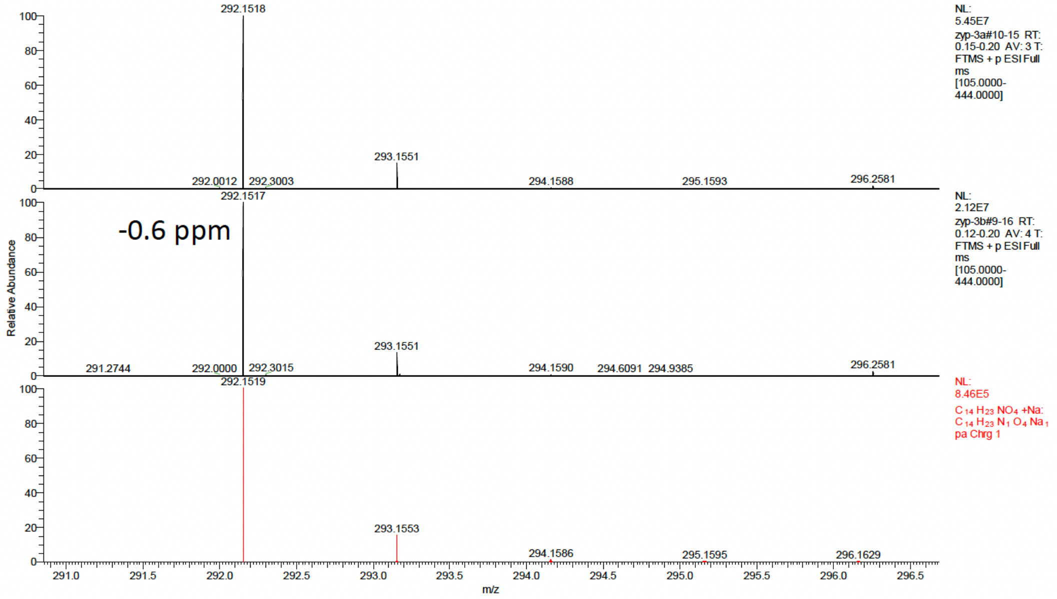


High resolution mass spectrum of **3a** and **3b** (top: experimental data of **3a**; middle: experimental data of **3b**; bottom: theoretical simulation)

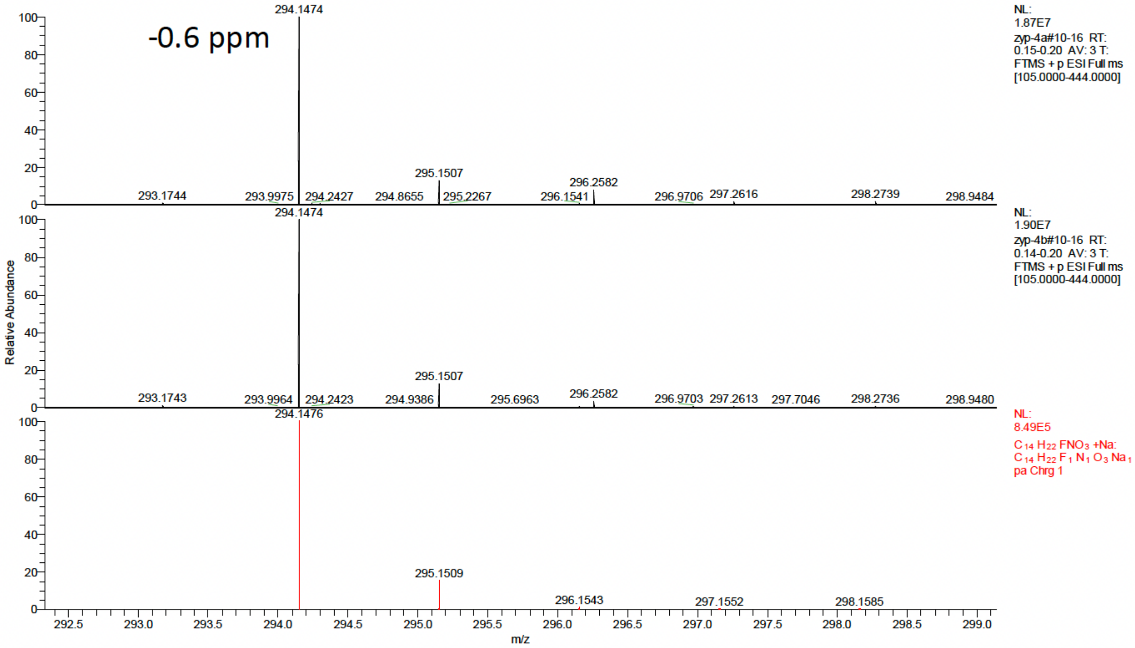


High resolution mass spectrum of **4a** and **4b** (top: experimental data of **4a**; middle: experimental data of **4b**; bottom: theoretical simulation)

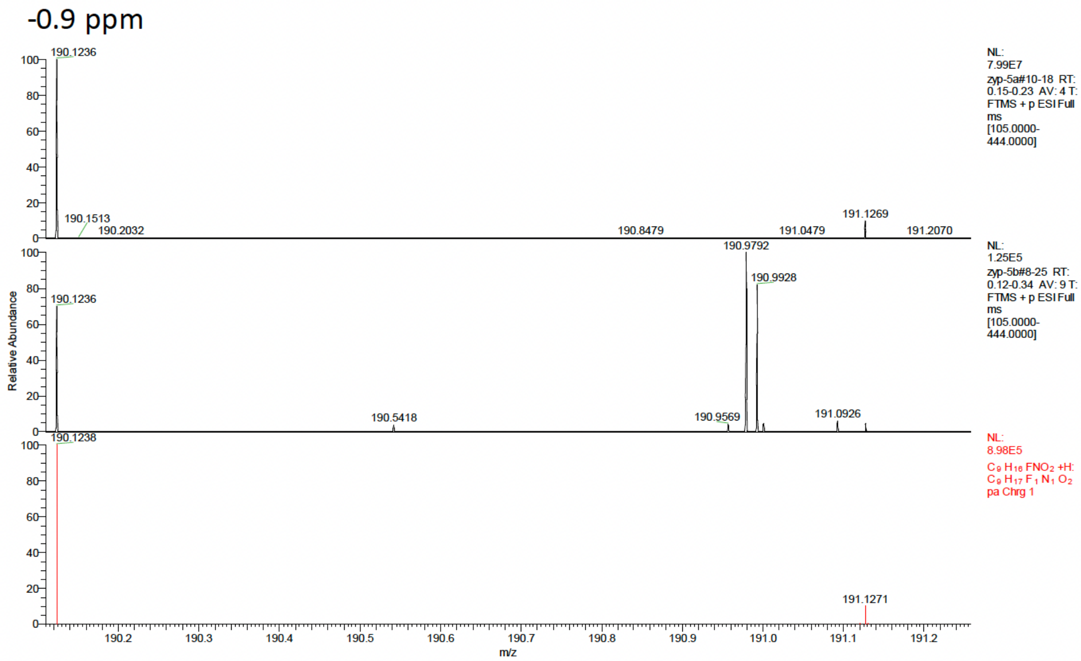


High resolution mass spectrum of **5a** and **5b** (top: experimental data of **5a**; middle: experimental data of **5b**; bottom: theoretical simulation)

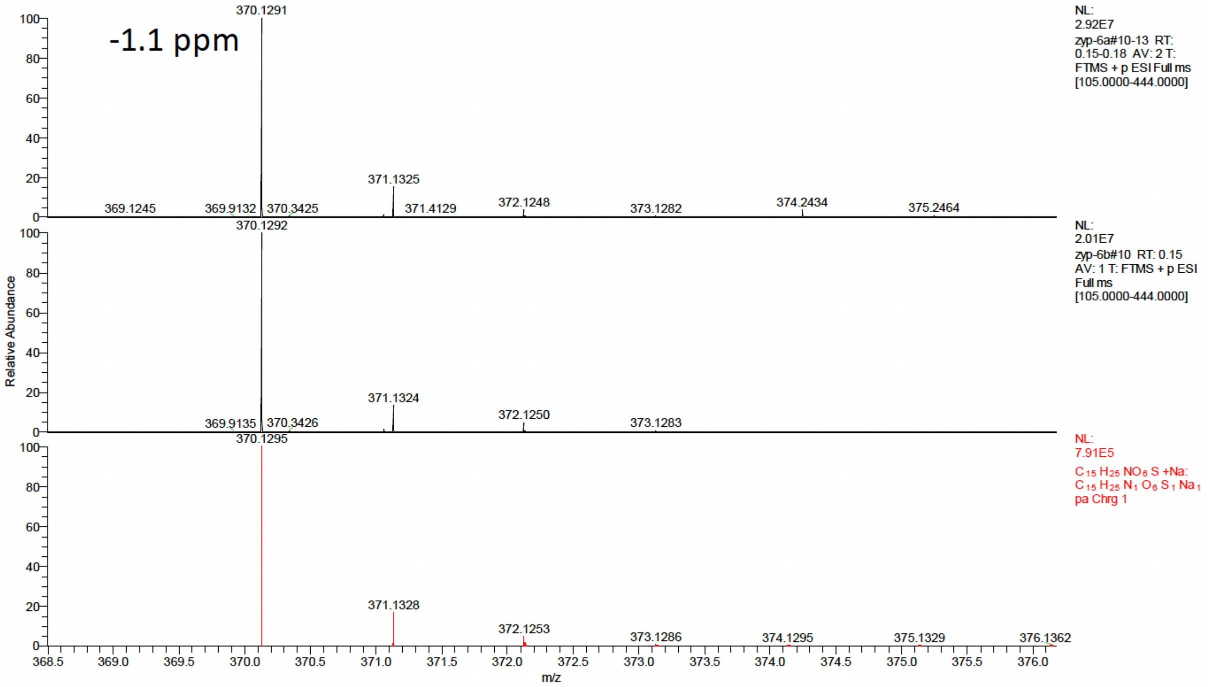


High resolution mass spectrum of **6a** and **6b** (top: experimental data of **6a**; middle: experimental data of **6b**; bottom: theoretical simulation)

### 3. HPLC chromatography

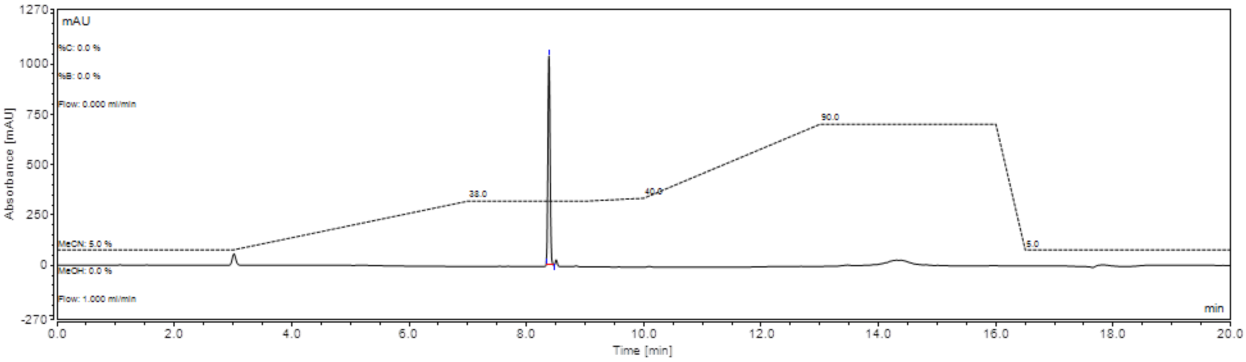

Analytic HPLC chromatography of **3a** (UV: 210 nm)

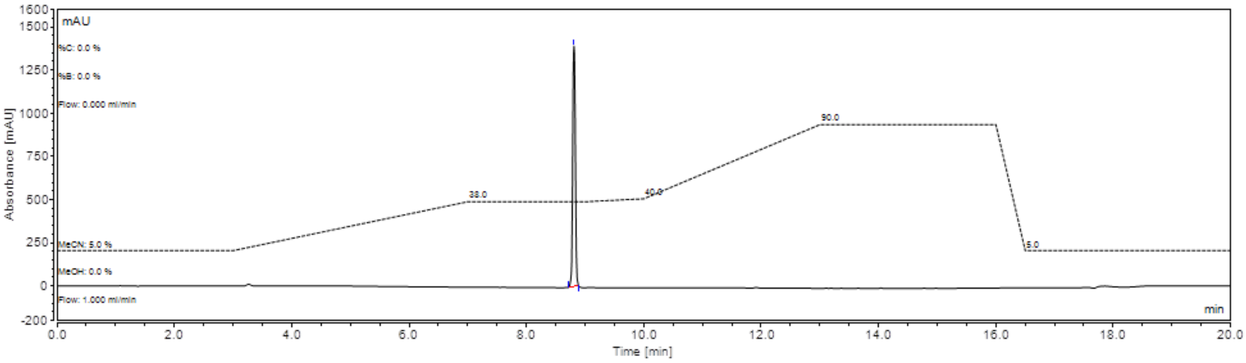

Analytic HPLC chromatography of **3b** (UV: 210 nm)

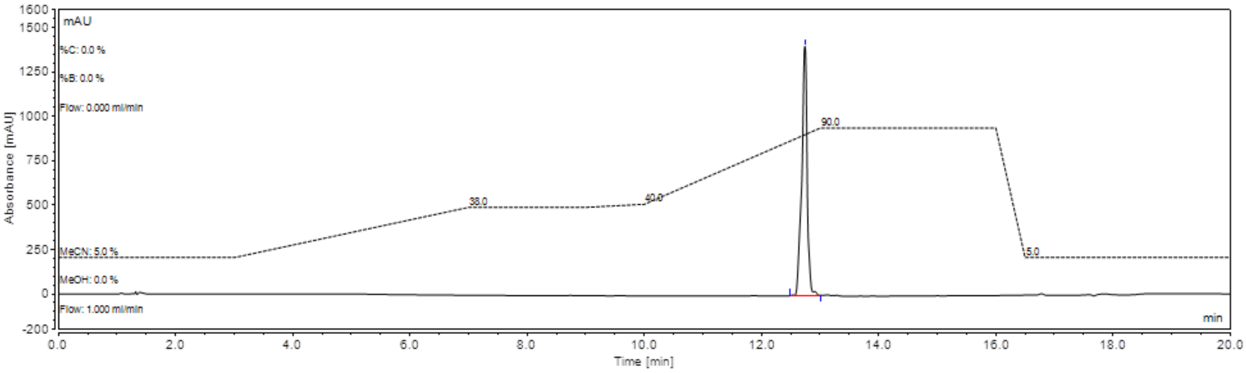

Analytic HPLC chromatography of **4a** (UV: 210 nm)

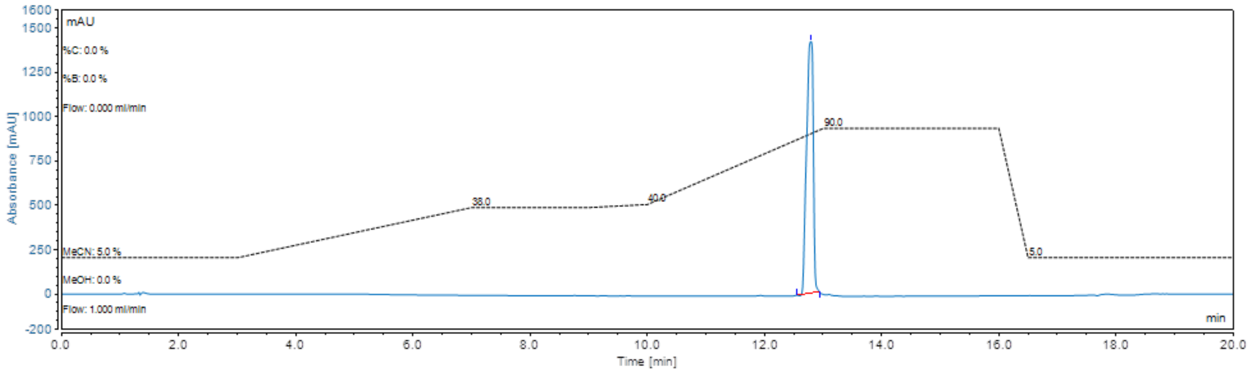


Analytic HPLC chromatography of **4b** (UV: 210 nm)

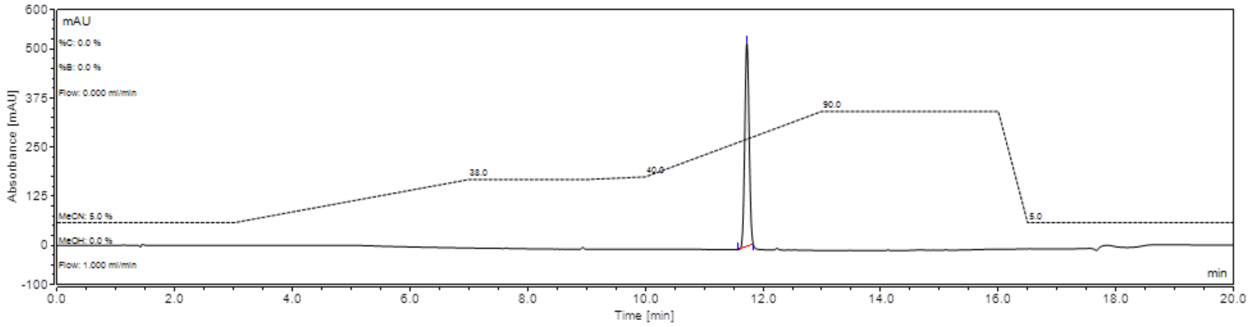


Analytic HPLC chromatography of **6a** (UV: 210 nm)

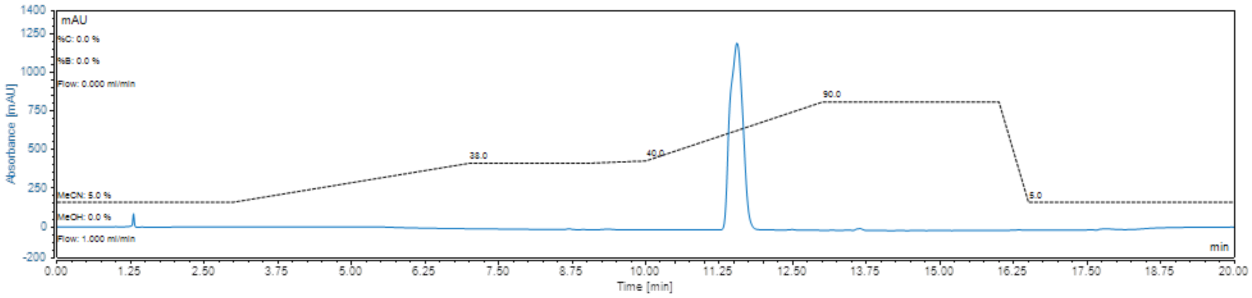


Analytic HPLC chromatography of **6b** (UV: 210 nm)

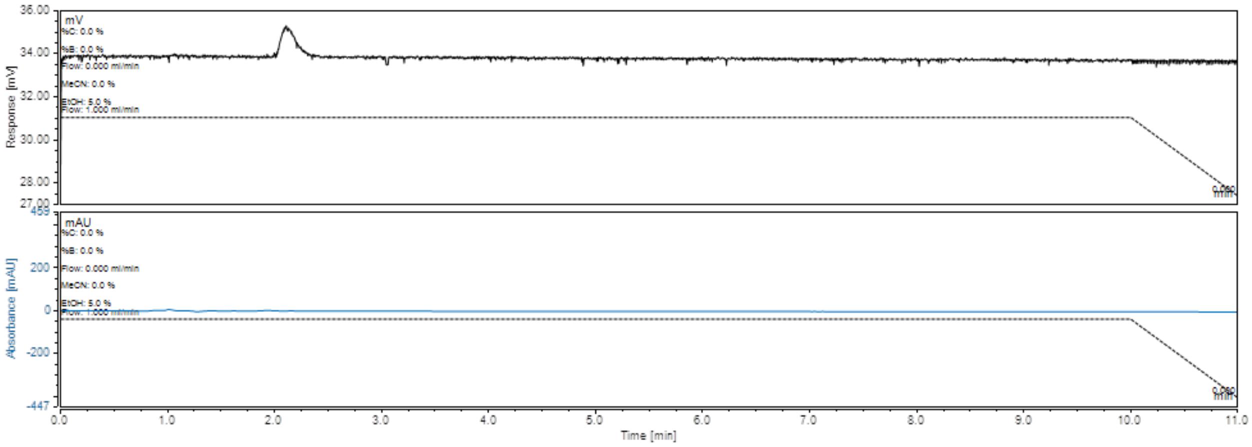


Analytic HPLC chromatography of [$^{18}$F]tGBP4F (top: radio-detector; bottom UV detector at 210 nm)

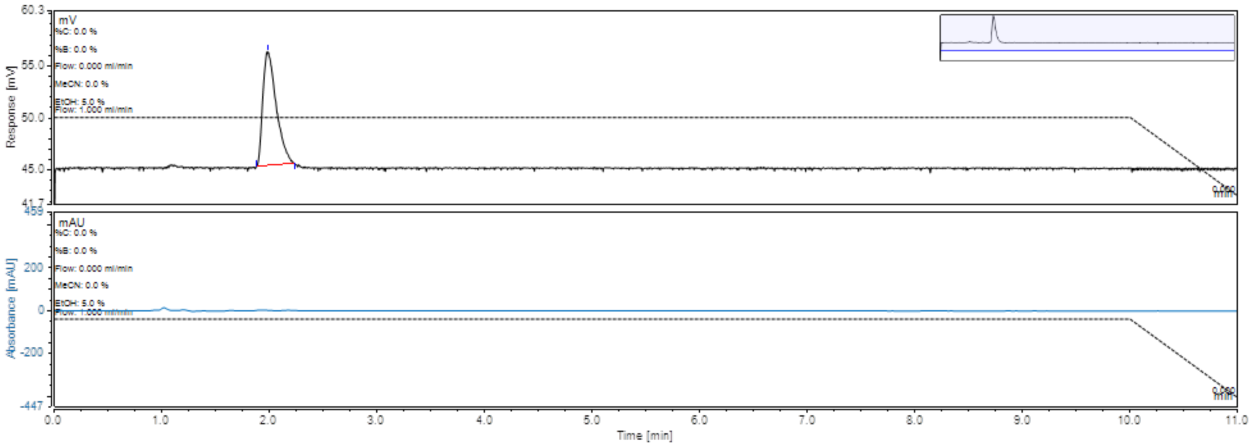


Analytic HPLC chromatography of [$^{18}$F]cGBP4F (top: radio-detector; bottom UV detector at 210 nm)

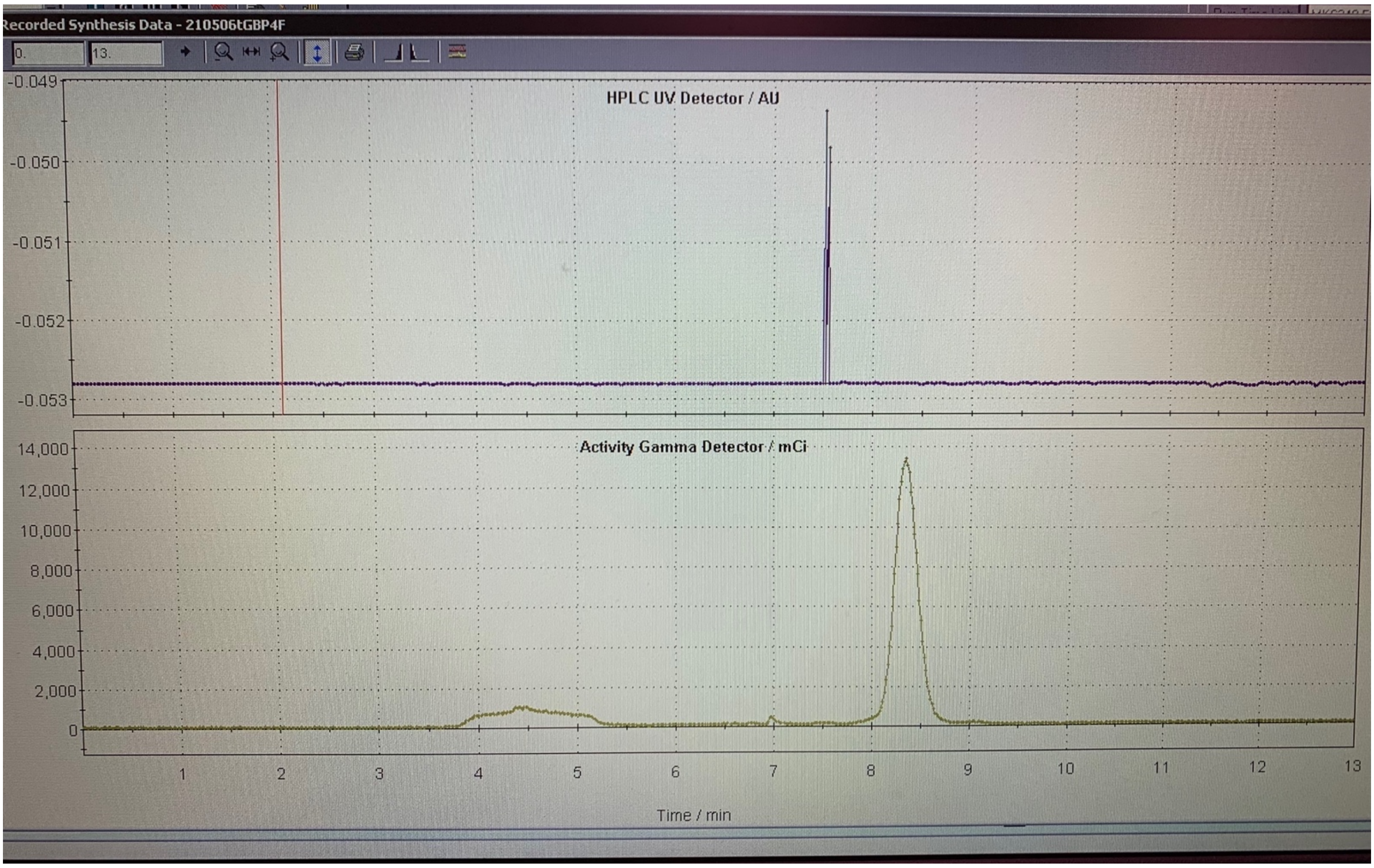


Semiprep HPLC chromatography of [$^{18}$F]tGBP4F

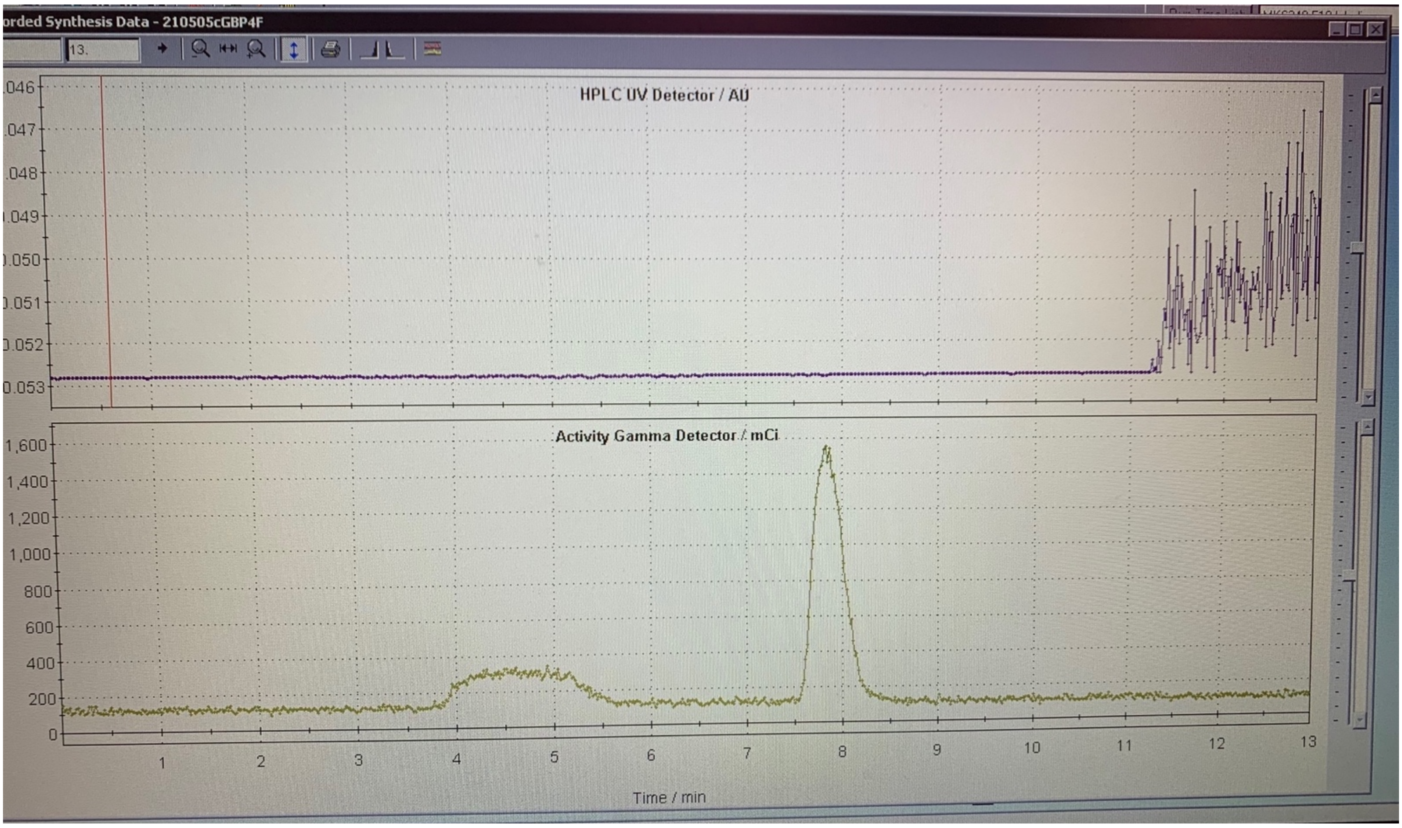

Semiprep HPLC chromatography of [$^{18}$F]cGBP4F
